# Integrative genomics reveals the polygenic basis of seedlessness in grapevine

**DOI:** 10.1101/2023.12.22.573032

**Authors:** Xu Wang, Zhongjie Liu, Fan Zhang, Hua Xiao, Shuo Cao, Hui Xue, Wenwen Liu, Ying Su, Zhenya Liu, Haixia Zhong, Fuchun Zhang, Bilal Ahmad, Qiming Long, Yingchun Zhang, Yuting Liu, Yu Gan, Ting Hou, Zhongxin Jin, Xinyu Wu, Yiwen Wang, Yanling Peng, Yongfeng Zhou

## Abstract

Seedlessness is a crucial quality trait in table grape (*Vitis vinifera* L.) breeding. However, the development of seeds involved intricate regulations, while the polygenic basis of seed abortion remains unclear. Here, we combine comparative genomics, population genetics, quantitative genetics, and integrative genomics to unravel the evolution and polygenic basis of seedlessness in grapes. We generated four haplotype-resolved telomere-to-telomere (T2T) genomes for two seedless grape cultivars, ‘Thompson Seedless’ (TS, syn. ‘Sultania’) and ‘Black Monukka’ (BM). Comparative genomics identified a ∼4.25 Mb hemizygous inversion on Chr10 specific in seedless cultivars, with seedless-associated genes *VvTT16* and *VvSUS2* located at breakpoints. Population genomic analyses of 548 grapevine accessions revealed two distinct clusters of seedless cultivars, tracing the origin of the seedlessness trait back to ‘Sultania’. Introgression, rather than convergent selection, shaped the evolutionary history of seedlessness in grape improvement. Genome-wide association study (GWAS) analysis identified 110 quantitative trait loci (QTLs) associated with 634 candidate genes, including novel candidate genes, such as three *11S GLOBULIN SEED STORAGE PROTEIN* and two *CYTOCHROME P450* genes, and well-known genes like *VviAGL11*. Integrative genomic analyses resulted in 339 core candidate genes categorized into 13 groups related to seed development. Machine learning based genomic selection achieved a remarkable 99% precision in predicting grapevine seedlessness. Our findings highlight the polygenic nature of seedless and provide novel candidate genes for molecular genetics and an effective prediction for seedlessness in grape genomic breeding.

## Introduction

The production of seedless fruits leads to tremendous success in the global fruit market^1^, such as bananas^2,3^, citrus^4,5^, watermelons^6,7^, and table grapes^8^. Seed abortion in table grape has been a major focus of breeding efforts for decades, as seedless grapes are highly preferred by consumers owing to improved tastes and convenience. There are two primary methods employed to obtain seedless grapes. One involving the application of phytohormones, applying gibberellin acid (GA) and cytokinin analogs before the full bloom stage can effectively induce seed abortion in seeded grapes^9–11^. Although this process has already become a common practice for producing seedless grapes, it raises concerns about food safety and labor costs^12^. Another one is based on genetic breeding of seedless grape cultivars. Breeders have explored diploid and triploid breeding approaches and the embryo rescue strategy to obtain new seedless varieties in recent decades^13–16^.

The development of various seed tissues in grapes involves intricate genomic regulations. Previous studies have identified specific genes associated with different tissues of seed development^17^. For instance, the formation of the seed coat (integument) has been affected by genes like *VviAGL11*^18–21^, *VvMADS28*^22^, *VviINO*^23,24^, and *VvHB63*^25–27^. Nutrient storage in the endosperm is controlled by genes such as 7S and 11S globulin-like seed storage proteins^28–30^, while normal embryo growth relies on several gibberellin (GA) genes^31,32^. Moreover, the growth of ovules (young seeds) is influenced by genes such as *VvMADS39*^33^, *VvMADS45*^34^, *VviABCG20*^35–37^, *VvFUS3*^38^, *VvNAC26*^39^, *Vv*β*VPE*^40,41^, *VviASN1*^42^. In general, multiple genes involved in tissue development of grapevine seeds. Seed abortion in grapes can occur when any of the seed tissues fail to develop properly. However, the polygenic basis of seedlessness in grapes remains unclear.

Previous studies have mapped multiple QTLs in different progenies in grapevine. In the linkage map of ‘Dominga’ and ‘Autumn Seedless’, three QTLs for seed number (SN) and six QTLs for seed fresh weight (SFW) were detected^43^. Similarly, in the ‘Muscat of Alexandria’ and ‘Crimson Seedless’ progeny, six QTLs for SN and ten QTLs for SFW were detected^44^. Recent comparative analyses, encompassing 28 different grape varieties(13 seeded and 15 seedless), have detected 34 candidate genes associated with the divergence between seeded and seedless lineages^45^. However, the restricted genetic background in progenies and limited population samples hinders the investigation of the polygenic basis of seedlessness in grapes.

In this study, we first generated haplotype-resolved T2T genomes for two seedless grapes, ‘Thompson Seedless’ (TS) and ‘Black Monukka’ (BM), and we compared these haplotype genomes with 11 other grape genomes to detect structural variations (SVs) between seeded and seedless genomes. Population genetic analysis was conducted on whole-genome sequencing (WGS) of 548 accessions to investigate the evolutionary history of seedlessness, while quantitative genetic analysis involved 444 accessions to map QTLs and key genes associated with seedlessness. Integrative genomic analysis incorporated three transcriptomes with 14 development stages, homologous genes related to 34 Gene Ontology (GO) terms, and 451 family genes and seven molecular markers previously reported with significant effects on seed development processes. Finally, genomic selection, based on polygenic model and machine learning algorithms, were applied in predicting the seedlessness trait in grapes. Collectively, we aimed to address five sets of questions. First, at genome level, how do the seedless cultivars compare with cultivars with seeds? Can we detect big SVs related to seedless/seeded cultivars? Second, what evolutionary factors has driven the origin of seedlessness during grape improvement? i.e., introgression or convergent selection? Third, based on large natural populations, can we map genetic loci and candidate genes involved in seed abortion in table grapes? Fourth, can we integrate genomic analyses to identify core candidate genes underlying seed abortion? Finally, can we employ machine learning based genome selection to enhance prediction precision in grape breeding? Overall, our work contributes to improving the understanding of the polygenic basis of seedlessness and facilitate genomic breeding of grapes.

## Results

### Comparative genomics between seeded and seedless cultivars

To study the genetic basis of seedlessness, we generated haplotype-resolved T2T assemblies for two seedless cultivars: TS and BM (**Fig. 1b, c)**, utilizing high-depth PacBio HiFi sequencing (∼120× coverage) and Hi-C sequencing (∼116× coverage; **Extended Data Fig. 2c**). The evaluation of K-mer heterozygosity in the TS and BM genomes, based on HiFi data, measured 1.51% and 1.41%, respectively. The quality of these genomes meets the assessment standards of the T2T level^46^, with all centromeres regions and mostly telomeres regions marked (**Fig. 1a and Supplementary Table 1, 3**). Statistical analysis of variants revealed that TS and BM, between their two haplotype genomes, harbor 5.35 Mb and 5.04 Mb of SNPs, 5.01 Mb and 4.54 Mb of insertions and deletions (InDels, < 50 bp), and 33.42 Mb and 31.87 Mb of SVs (≥ 50 bp), respectively (**Supplementary Table 4**). A unique heterozygous inversion region (PN_T2T, Chr15: 10.7-12.0 Mb) specific to the TS hap2 genome was detected when aligning the four haplotypic genomes to PN_T2T, and several genes were found near the inversion breakpoints, such as *AGAMOUS* (*VvAG2*), *AGAMOUS-LIKE 62* (*VvAGL62*), *OIL BODY-ASSOCIATED PROTEIN 2B* (*VvOBAP2B*), and *GDSL esterase/lipase At1g29670*, which are involved in stamen and carpel determining, early endosperm development, and oil body synthesis (**Extended Data Fig. 3b and Supplementary Table 5**). The differential chromosomes between the two haplotype genomes explains the polymorphism of alleles and the variation in the number of genes (**Supplementary Table 1**).

**Fig. 1.**
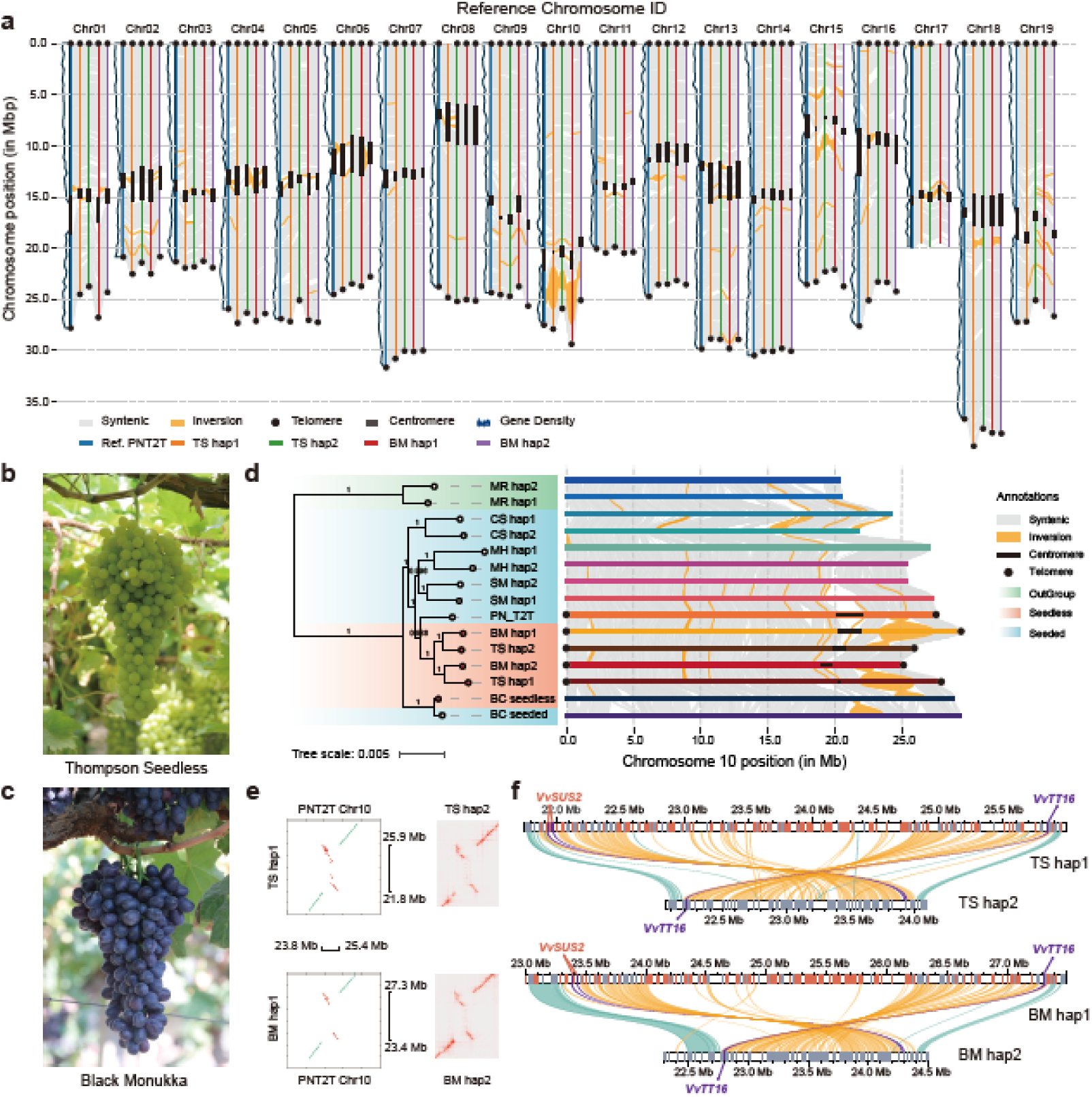
Comparative genomic between seeded and seedless grape cultivars. **a**, Visualization of the haplotype-resolved genomes aligned with the PN_T2T, with discernible inversions marked in yellow. **b** and **c**, Morphological features of TS and BM. **d**, Comparative genomic analysis of 15 grape genomes prioritized by the phylogenetic tree constructed with single-copy gene features. Included genomes: MR (Muscadinia rotundifolia), CS (Cabernet Sauvignon), MH (Muscat Hamburg), SH (Shine Muscat), PN (PN40024), BC (Black Corinth). **e**, Detailed validation of a prominent Chr10 segment inversion, with its distinctive features illustrated in the Hi-C heatmap. **f**, Gene loss in hap2 genomes within the Chr10 inversion region. *VvSUS2* and *VvTT16* are located near the inversion boundaries. Red blocks represent lost genes, while gray blocks indicate genes shared between the two haplotype genomes.

To further investigate the SVs associated with seedless and seeded grapes, we aligned a total of 15 genomes, including the five seedless genomes and ten seeded genomes, to the PN_T2T (**Extended Data Fig. 3a**). We detected a heterozygous inversion (PN_T2T, Chr10: 23.8-25.4 Mb) in seedless grape varieties (**Fig. 1d**). The authenticity of these inversions was confirmed by Hi-C heatmaps and IGV^47^ (**Fig. 1e and Extended Data Fig. 4**). A total of 210 genes (Chr10: 21.75-26.00 Mb) and 237 genes (Chr10:23.00-27.50 Mb) were identified in the inversion regions of TS hap1 and BM hap1, respectively (**Fig. 1f and Supplementary Table 7-8**). Three seed development-related genes, *TRANSPARENT TESTA 16/ ARABIDOPSIS BSISTER* (*TT16/ABS*) and two *SUCROSE SYNTHASE 2* (*SUS2*) genes, were discovered near the breakpoints of inversion region of the haplotype genomes. *TT16/ABS* controls the formation of the maternal-derived endothelial cells by interacting with *AGL11/ SEEDSTICK* (*STK*) in the previous studies^48–51^. *VvTT16* was found to be present in both TS and BM haplotype genomes, while the two *VvSUS2* tandem duplication genes were hemizygous, present only in the hap1 genome of TS and BM (**Supplementary Table 6-7**). These findings suggest that the power of comparative genomics of haplotype-resolved T2T genomes in uncovering overlooked new candidate genes underlying crucial agronomic traits.

### Introgression rather than convergent evolution underlying the evolvement of seedlessness in grapevine

To explore the evolutionary history of seedlessness in grape improvement, we used WGS data from 548 grapevine accessions, including 46 seedless grapes, for population genetic analysis (**Supplementary Table 8)**. A total of 4,462,797 SNPs, 443,812 InDels, 487,204 SVs were identified by aligning WGS data to ‘Cabernet Sauvignon’ (CS) genome^52^. The phylogenetic tree split into six primary population branches: European wild grapes (*V. vinifera* ssp. *sylvestris* EU population, EU, n = 69), Middle East and Caucasus region wild grapes (*V. vinifera* ssp. *sylvestris* ME population, ME, n = 23), domesticated grapes (*V. vinifera* ssp. *vinifera*, VV, n = 352), American fox grapes (*V. labrusca*, VL, n = 5), hybrid of VV and VL grapes (VV×VL, *V. vinifera × Vitis labrusca*, n = 92), and outgroup grapes (OG, n = 7; **Fig. 2c and Extended Data Fig. 5**). The results showed two independent lineages of seedless grapes nested in the VV×VL and VV branches, respectively, which is also supported by PCA (**Fig. 2b, c**). The seedless grapes nested in two branches, which could be driven by convergent artificial selection or introgression during grape improvement.

**Fig. 2.**
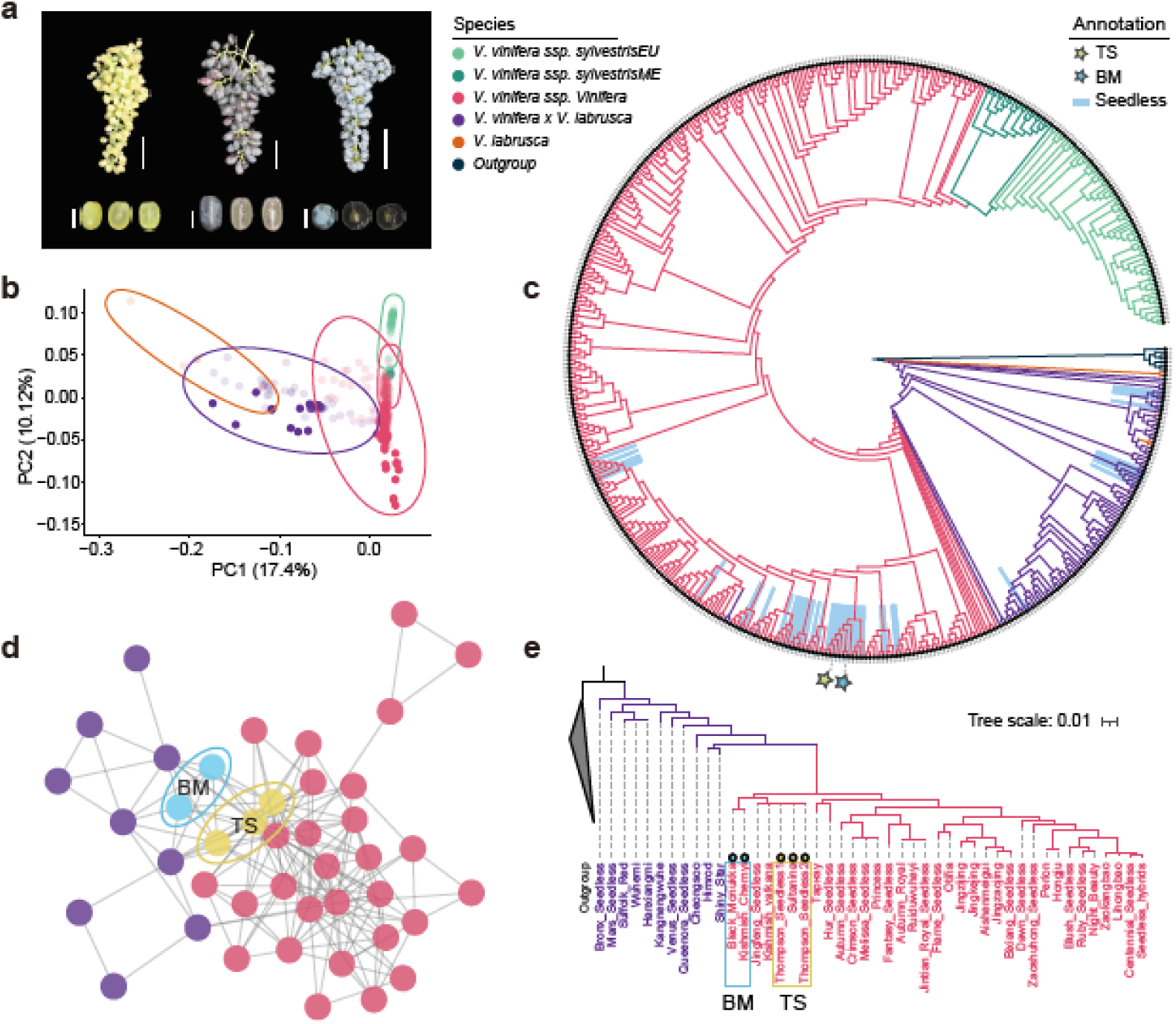
Evolutionary genomics of seedlessness in grapevine. **a**, Comparison of grape clusters and cross-sectional views of berry from three cultivated grape varieties: TS, BM, and CS. The scale bar indicates 5 cm for grape clusters and 1 cm for individual berries. **b**, PCA analysis on the whole-genome sequencing (WGS) data, except the outgroup, resulting in obviously separated five populations. Solid dots denote seedless grapes, while transparent dots represent seeded grapes. CS genome serves as the reference genome for variants calling. **c**, Phylogenetic tree analysis of the six populations. The light-blue blocks represent seedless grapes and star symbols indicate TS and BM (see Extended Data Fig. 6 for detailed information of the phylogenetic tree). **d**, IBD analysis of 46 seedless grapes, filtering the results with a threshold of 0.40. Purple points represent the *V. vinifera × V. labrusca* population, while red points represent the *V. vinifera* population. **e**, The phylogenetic tree of 46 seedless grapes.

To distinguish convergent selection and introgression in generating seedless traits in different grapevine lineages, we preformed the introgression analyses, throughout the whole genome using *f*_d_ statistics^53^, following previous studies^54,55^. Interestingly, we detected significant genomic signals of introgression at seedless associated locus (see the GWAS section) between VV and VV×VL seedless grapes (the upper 5th percentile, *f*_d_ = 0.266, *P* = 1.28e-39), including the redefined *SEED DEVELOPMENT INHIBITOR* (SDInew, 30.36-31.86 Mb) locus^20^ and the newly detected QTL on Chr07 (SDI2, 8.85-8.86 Mb; **Fig. 3c, d**). These results suggested that introgression rather than convergent artificial selection has driven the evolutionary history of seedlessness in grapes.

**Fig. 3.**
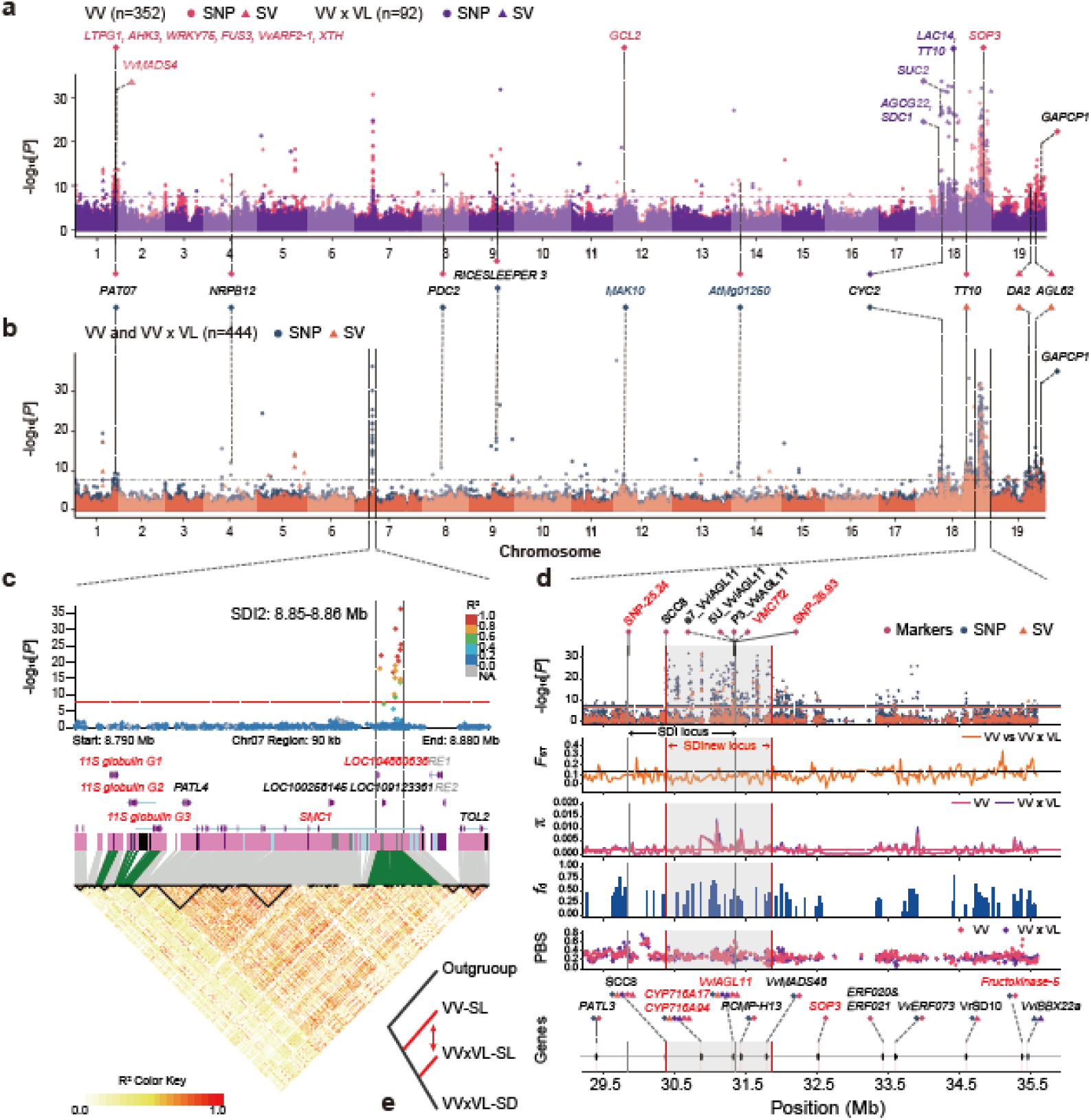
GWAS mapping of the polygenic basis of seedlessness. **a** and **b**, Seedless-associated genomic loci and genes across three populations: the VV population (in red, n=352), the VL population (in purple, n=92), and the admixed population (in yellow and dark-blue, n = 444). The horizontal dashed lines denote the Bonferroni thresholds (-log_10_[0.05/Variant Numbers]): 7.62 for VV, 7.45 for VV *×* VL, and 7.66 for admixed population, respectively. Points represent SNPs, while triangles represent SVs (and InDels). **c**, LD correlation analysis of a highly linked SDI2 locus in Chr07, and genes associated with seed abortion are highlighted in red. **d**, Admixed population analysis of a highly linked region in Chr18. The SDI locus (Chr18: 29.83-31.34 Mb) is identified based on SNP markers, and the redefined SDI locus (SDInew, Chr18: 30.36-31.86 Mb) is defined based on GWAS results, depicted in the grey block. The significant threshold: 7.61 for SNPs, and 6.66 for SVs (and InDels). The dashed line for fixation indices (*F*_ST_) indicates an average value of 0.126, with genetic diversity (π) measured 0.0019 for VV*×*VL and 0.0016 for VV. The top 5th percentile of *f*_d_ statistics is 0.277, with PBS statistics measuring 0.356 in VV and 0.415 in VV*×*VL. **e**, Gene flow pattern based on ABBA-BABA statistics, where SD represents Seeded, SL represents Seedless.

To validate the genetic relationship among seedless varieties, a network was constructed comprising 46 seedless grapes (35 VV and 11 VV×VL) based on the results of Identity-by-Descent (IBD) analysis (**Supplementary Table 8**). The result revealed that TS group and BM group serve as bridges for gene flow between VV and VV×VL clusters (**Fig. 2d, e**). In fact, BM grape traces its ancestry back to ‘Sultania’ (TS) and ‘Ichkimar’^56^, with an IBD score of 0.50 supporting this observation (**Supplementary Table 9**). Additionally, we identified three grapes belonging to the ‘Sultania’ somatic variants or synonym group (‘Jingfeng seedless’, TS1, and TS2, IBD > 0.95), seven grapes classified as parent-offspring relationship (IBD > 0.50), and 32 grapes (0.50 > IBD > 0.09) that were closely connected to ‘Sultania’ variety (**Supplementary Table 9**). These findings provide evidence that ‘Sultania’ had been extensively employed in crossbreeding with local grape varieties to enhance quality and develop new seedless grape cultivars^8,57^. Furthermore, the seed abortion could also be caused by cytoplasmic male sterility (CMS)^58^. We analyzed the chloroplast and mitochondrial genomic variation and found nuclear inheritance, rather than CMS inheritance, played a crucial role in controlling seed abortion (**Extended Data Fig. 6).** These findings corroborated IBD results that frequent introgression has facilitated the formation of seedlessness the VV and VV×VL populations (**Fig. 3e**). As a result, the origin of the seedlessness trait could be traced back to ‘Sultania’, and continuous introgression, rather than convergent evolution, led to seed abortion in seedless grape varieties.

### Genome-wide Association Study for Seed Abortion Trait

To detect the QTLs and genes associated with seed abortion, we used three population for GWAS analysis, considering the effects of population structure in GWAS analyses: VV (35 seedless and 317 seeded grapes), VV×VL (11 seedless and 81 seeded grapes), and an admixed population (46 seedless and 398 seeded grapes; **Supplementary Table 8**). We identified a total of 110 QTLs (634 genes), including 20 QTLs (126 genes) specific to the VV population, 18 QTLs (106 genes) specific to the VV×VL population, and 72 consensus QTLs (402 genes) in admixed population, respectively (**Extended Data Fig. 7 and Supplementary Table 10, 11**). GO analysis revealed that genes specific to VV×VL were enriched in defense response and lignin catabolic process (*P* < 0.05), while genes specific to VV population were enriched in embryo development ending in seed dormancy and xylan metabolic process (*P* < 0.05; **Extended Data Fig. 8 and Supplementary Table 12**).

Remarkably, two consensus regions exhibited high consistence in three populations: Chr07: 8.85-8.86 Mb and Chr18: 29.40-35.54 Mb (**Fig. 3a, b**). In Chr07 locus (SDI2), two genes, *REVERSE TRANSCRIPTASE ZINC-BINDING DOMAIN-CONTAINING PROTEIN* (*LOC104880636*, *Vitvi011893*) and *STRUCTURAL MAINTENANCE OF CHROMOSOMES PROTEIN* (*SMC1*, *Vitvi011891*), were positioned within a tightly linked region with high linkage disequilibrium (LD) values (**Fig. 3c**). The *LOC104880636* gene is annotated as the regulation of seed growth on UniProt, and the *smc1* mutants produced arrested early embryo development and blocked cellularization of the endosperm^59^ (**Supplementary Table 11**). Additionally, within the 50 kb upstream region of SDI2, we identified a closely linked cluster of three tandem-duplicated genes, *11S GLOBULIN SEED STORAGE PROTEIN*^28–30,60^ (**Fig. 3c**), and designated them as *11S globulin G1*, *G2*, and *G3* based on their genomic positions. Notably, two nonsynonymous mutations and one deletion related to seedlessness were detected across 14 grape genomes (**Fig. 4a and Extended Data Fig. 9**). Among them, both the heterozygous Asp-to-Val and Leu-to-Val mutations were specific in seedless grapes, except for the somatic mutations of ‘Black Corinth’ (BC) seeded grapes. However, the heterozygous deletion of the 18 amino acids in *11S globulin G3* was only detected in seedless grapes (**Fig. 4a**). The relative expression values of these genes showed a significant correlation with seed phenotypes, especially from 40-50 days after flowering (DAF; **Fig. 4b**).

**Fig. 4.**
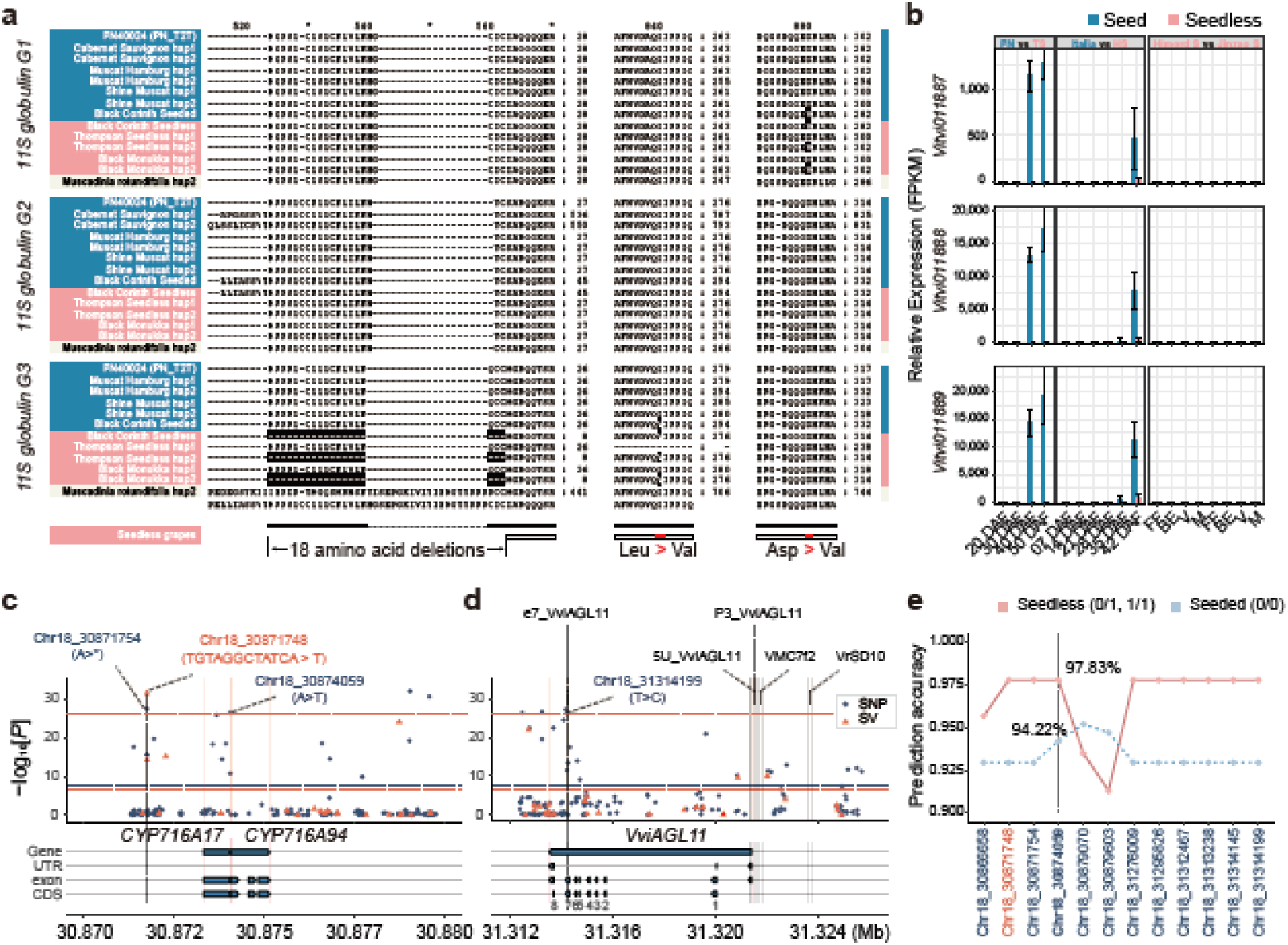
Deep mining of key loci associated with seed abortion. **a**, Sequence alignment of three *11S GLOBULIN SEED STORAGE PROTEIN* genes in Chr07 across 14 grape varieties. Black blocks represent deletion and nonsynonymous mutations. **b**, Relative expression values of these three genes at different time points across six grape varieties. Abbreviations: DAF (Day after flowering), FF (Full flowering), BE (Berry expansion), V (Veraison), M (Maturity). **c** and **d**, Visualization of crucial candidate genes associated with seed abortion within the SDInew locus region. GWAS significance thresholds (-log_10_[0.05/Variant Numbers]): 7.61 for SNPs, and 6.66 for SVs (and InDels). The top ten percentiles of significant variants are Chr18_ 31295826 (y = 26.39). **e**, Genotyping percentage of highly significant variants is within the region Chr18: 30.70-31.32 Mb. ‘Seedless’ includes cases with genotypes 0/1 and 1/1, ‘Seeded’ includes the case with genotype 0/0.

Another consensus region is located on the Chr18: 29.40-35.54 Mb, encompassing 17 QTLs and the reported SDI locus (Chr18: 29.83-31.34 Mb; **Fig. 3d**). Due to the narrow genetic background of the samples used in the previous study^20^, we redefined the SDI locus (SDInew, Chr18: 30.36-31.86 Mb) through GWAS analysis, revealing a total of eight QTLs (**Fig. 3d and Extended Data Fig. 7**). In this region, the population differentiation was lower than the genomic background between VV (n = 35) and VV×VL (n = 11) seedless grapes, as showed by the fixation indices (*F*_ST_) results, suggesting a relatively close genetic distance between the two populations (SDInew *F*_ST_ = 0.073 vs. genome-wide *F*_ST_ = 0.126; **Supplementary Table 13**). This finding was also supported by genetic diversity (π) statistics. The two seedless populations showed similar genetic diversity on the SDInew locus (**Fig. 3d**). The *f*_d_ statistics^53^ revealed numerous sites showing evidence of introgressions (*f*_d_ = 0.266, *P* = 1.28e-39), including numerous genes related to seedlessness, such as *CYP716A94*, *CYP716A17*, *VviAGL11*, etc. (**Fig. 3d**). Notably, several SNPs and InDels were highly associated with these candidate genes, especially in the promoter and coding sequence region (**Fig. 4c, d and Extended Data Fig. 10**), while several published molecular markers for seedlessness prediction, including e7_VviAGL11^44^, 5U_VviAGL11^44^, P3_VviAGL11^18^, and VMC7f2^61^, showed low predictive accuracy due to the absence of significant genotyping quality (**Fig. 4d and Supplementary Table 14**). We selected the top 10% of associated sites based on −log_10_[*P*] values within the SDInew locus (**Fig. 4e**), and the most promising site for seedlessness prediction is Chr18_30874059, exhibiting a predicted accuracy of 97.8% for seedless grapes and 94.22% for seeded grapes in natural population.

Interestingly, we also detected two genomic regions for specific to each population (**Extended Data Fig. 7**). In the VV population, a specific region Chr01: 17.85-20.42 Mb harbored 13 QTLs and 67 genes, including seven primary genes involved in seed development, such as *NON-SPECIFIC LIPID TRANSFER PROTEIN GPI-ANCHORED 1* (*LTGP1*), *ARABIDOPSIS HISTIDINE KINASE 3* (*AHK3*), *B3 DOMAIN-CONTAINING TRANSCRIPTION FACTOR FUS3* (*FUS3*), *XYLOGLUCAN ENDOTRANSGLUCOSYLASE PROTEIN* (*XTH*), *11-BETA-HYDROXYSTEROID DEHYDROGENASE B* (*SOP3*), as well as previously reported *VvMADS4*^62^ and *VvARF2-1*^63^ (**Fig. 3a**). In the VV×VL population, a specific region Chr18: 15.14-20.57 Mb harbored eight QTLs and 33 associated genes. Among these genes, *SERINE DECARBOXYLASE 1* (*SDC1*), *ABC TRANSPORTER G FAMILY MEMBER 22 ABCG22* or *VvPNWBC22.2*(TANG, 2018 #20), *SUS2*, *LACCASE-14* (*LAC14*) and *TT10*/*LAC15* were highly associated with seed development (**Fig. 3a**). Our results suggest that seed abortion can be regulated by the collaborative effects of multiple genes, and the mapped candidate genes and variable sites hold valuable potential applications in seedless grapes breeding.

### Integrative Genomic Analysis Identified 339 Seedless Candidate Genes

To elucidate the polygenic basis of seed abortion, we further utilized an integrative genomic analysis using transcriptomic analyses, seed development associated GO term genes, previously reported family genes and molecular markers, and GWAS candidate genes, to identify the core candidate genes associated with seed abortion (**Supplementary Table 17**). Among these, three transcriptomic groups, including 76 samples and 14 time points, were employed to detect differentially expressed genes (DEGs) between seeded and seedless grapes at each development stage (**Supplementary Table 15**). For the ‘Italia’ and ‘Hongju Seedless’ (HS) groups, we identified a total of 2,680 significantly upregulated genes and 1,835 significantly downregulated genes from the six time points (**Extended Data Fig. 13c**). Similarly, in the ‘Pinot Noir’ (PN) and TS groups, we identified a total of 3,969 significantly upregulated genes and 2,695 significantly downregulated genes (**Extended Data Fig. 13c**). ‘Himrod Seedless’ (Himrod) and ‘Jinzao Wuhe’ (Jinzao) were used serve as a control for cross-validation during four fruit development stage. Interestingly, we found *VviAGL11* was exclusively in the downregulated DEGs in the PN and TS comparison, but not in the ‘Italia’ and HS comparison (**Extended Data Fig. 13a, b**). In addition, 1,301 core upregulated genes and 616 core downregulated genes were only identified in transcriptomic analyses using integrative genomic analysis, such as *DORMANCY-ASSOCIATED PROTEIN HOMOLOG 4* (*DRM1 homolog 4*), *NON-SPECIFIC LIPID-TRANSFER PROTEIN 2* (*LTP2*), *VICILIN-LIKE SEED STORAGE PROTEIN*, *7S SEED STORAGE PROTEIN* (*7S GLOBULIN*), *TT10*, *LAC17*, etc. (**Fig. 5a and Supplementary Table 11**).

**Fig. 5.**
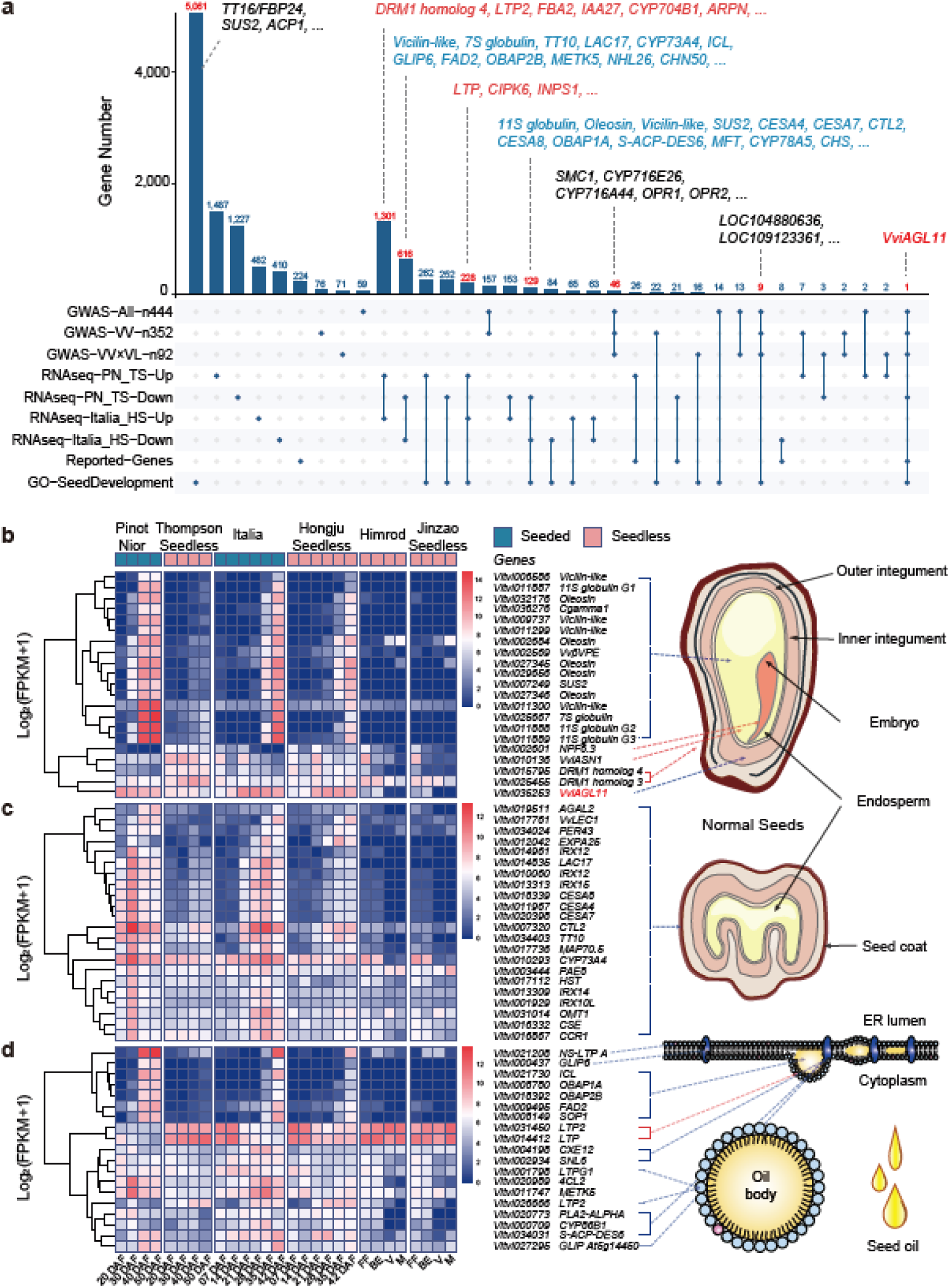
Integrative genomic analyses for grapevine seed abortion. **a**, Results from integrative genomic analyses: GWAS, transcriptomic analysis, reported genes and markers mapping, and GO homologous genes overlapping. Blue bars represent the number of genes uniquely and overlappedly through this approach. **b** and **c**, Heatmap of Log_2_(FPKM) for candidate genes related to embryo, endosperm, and seed coat development, resulting from integrative genomic analyses. The genes are indicated in relation to their expression in specific tissues and time points on schematic representation transverse profiles of seeds. **d**, Expression patterns of genes involved in lipid synthesis, degradation and transportation, which also pinpointed on the schematic representation of oil body formation. Abbreviations: DAF (Day after flowering), FF (Full flowering), BE (Berry expansion), V (Veraison), M (Maturity).

To include more important genes involved in seed development, we performed a protein sequence similarity alignment for all genes associated with 34 GO terms related to seed development against the PN_T2T reference genome (**Extended Data Fig. 12a, b**). This yielded 6,529 homologous genes (see Methods), with 5,061 genes only present in the GO pathway, including the *TT16/FBP24* gene identified in the comparative genomics (**Fig. 5a**). Notably, the intersection between the RNA-seq related genes and the GO homologous genes revealed 163 downregulated genes and 294 upregulated genes (**Extended Data Fig. 13b**), such as *LTP*, *11S GLOBULIN*, *OLEOSIN*, *CELLULOSE SYNTHASE A CATALYTIC SUBUNIT* (*CESA4*, *CESA7* and *CESA8*), etc. (**Supplementary Table 11**). The consensus GWAS genes and GO homologous genes exhibited nine candidate genes, such as previously mentioned *LOC104880636* (**Fig. 3c, 5a**). Furthermore, we integrated a total of 451 family genes and seven molecular markers related to seed abortion from previous studies (**Supplementary Table 16**). All these elements were aligned against the PN_T2T reference genome (e-value < 0.1) and exhibited overlap with candidate genes in other analyses, including previously reported genes such as *Vv*β*VPE*^40^, *HD-ZIP PROTEINS ATHB-1/HAT5*, *ATHB-12/VvHB56* and *ATHB40/VvHB18*^25^, *VviASN1*^42^, *VvMJE1*^64^, *VvLEC1*^65^, *VvMADS2/VvSEP1*^66^, etc. (**Extended Data Fig. 13d and Supplementary Table 11**).

Through integrative genomic analysis, we screened 339 core candidate genes, categorized into 13 groups, by condition-based filtering that exhibited significant differential expression between seedless and seeded grape cultivars (see Methods, **Supplementary Table 11**). Among them, 77 genes were directly associated with seed development-related GO homologous genes, and three groups deserve our attention: Firstly, the differential expression of candidate genes in the endosperm development impacts nutrient storage in seedless grapes (**Fig. 5b**). Nutrient deficiency could be primary factor leading to seed abortion in later embryo development; Secondly, genes involved in the regulation of lignin and cellulose synthesis/degradation in seed coat exhibit higher activity levels in seeded grapes (**Fig. 5c**); Thirdly, candidate genes manage the synthesis and transport of oil bodies ensuring efficient lipid accumulation and utilization during seed development (**Fig. 5d**). Overall, our findings indicate that multiple genes have an accumulative effect in the process of seed abortion, resulting varying degrees of seedlessness^67^. This complexity highlights the challenge of accurately distinguish seed abortion using a single gene or variant site.

### Machine Learning based Genomic Selection for Seedless Grape Breeding

Given the polygenic nature of the seedlessness trait, genomic prediction could greatly improve the speed and accuracy for seedless grape breeding. We extracted the information of all 794 high-quality variant sites from GWAS analysis, including 77 InDels and 717 SNPs (**Supplementary Table 18**). Using these variations, an unrooted phylogenetic tree was constructed based on admixed populations (n = 444), revealing that the majority of seedless individuals clustered together (**Fig. 6a**). However, some seeded samples, like seeded hybrid progeny^20^ and Rizamat^44^ were mixed in seedless grapes, as well as seedless samples, such as ‘Dawn Seedless’, ‘Bronx Seedless’, ‘Cheongsoo’, ‘Ruby Seedless’, and ‘Jingkejing’, were mixed in seeded grapes. Interestingly, the mutations in the top two QTLs were heterozygous: Chr07: 8.85-8.86 Mb (SDI2 locus) and Chr18: 30.36-31.86 Mb (SDInew locus; **Fig. 6d**). These results suggest the complexity of seedless in grapes.

**Fig. 6.**
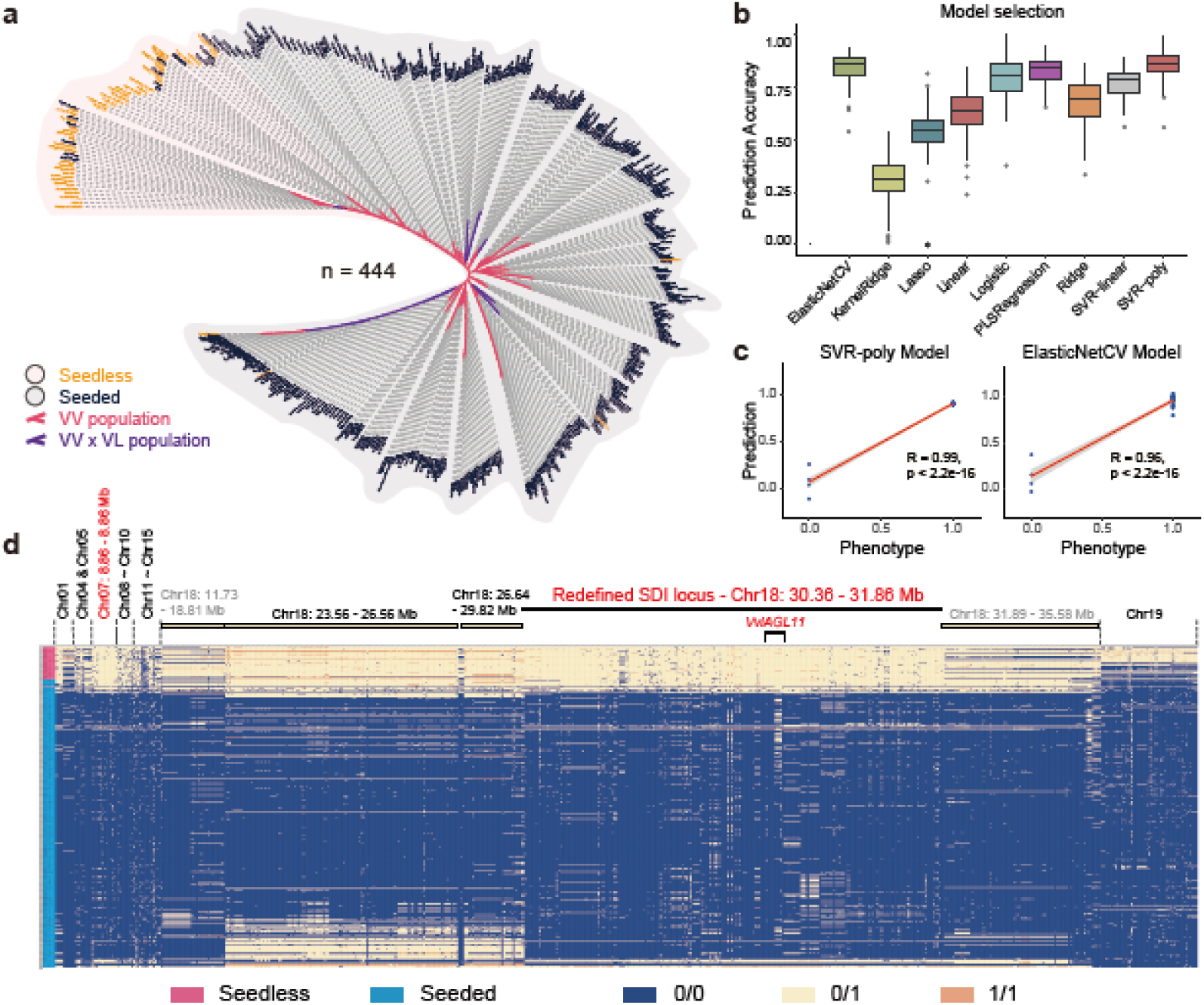
Machine learning based genomic selection on seedlessness in grapevine breeding. **a**, Phylogenetic clustering based on 794 significant variants (77 InDels and 717 SNPs) derived from the GWAS analysis. **b**, Comparison of seedlessness prediction accuracy across nine classical models for genome selection. The training set comprises of 444 grape samples and their phenotypes. **c**, Prediction results of the two best-performing models. The testing set includes 39 samples, distinct from the 444 samples used for model training. **d**, Genotyping visualization of the 794 variants, sorted based on prediction results of the SVR-ploy model.

Therefore, to address this problem, we employed genomic selection based on machine learning to enhance predictive accuracy (**Extended Data Fig. 14**). We used 794 variant sites and phenotypic data from the admixed populations (444 samples) as the training dataset, and evaluated the ability of nine different classical models to predict the seedlessness trait based on 100 rounds of random cross-validations (**Fig. 6b**). Among these models, machine learning based methods, including SVR-poly and ElasticNetCV, demonstrated a strong performance in predicting the phenotype, yielding prediction accuracies of 85.36% and 84.57%, respectively (**Fig. 6b**). As a result, we applied SVR-poly and ElasticNetCV to genomic prediction on the testing set data (39 samples not included in the model building) (**Supplementary Table 8**). We observed that the SVR-poly model and the ElasticNetCV model yielded high levels of accuracy, with correlation coefficient R values of 0.99 (*P* < 2.2e-16) and 0.96 (*P* < 2.2e-16), respectively (**Fig. 6c**), suggesting the efficiency of machine learning based genomic selection of seedless in grapes.

## Discussion

Seedless is an important quality trait in table grape breeding. Previous genetic investigated the functions of multiple genes, including *VvAGL11*, however, quantitative genetic analyses of natural population were not conducted in grapes. In this study, we conducted integrative genomic analyses to investigate the polygenic basis of seedlessness in grapes: (1) comparative analysis of 15 genomes allowed us to discover a heterozygous inversion in Chr10 associated with the seedless trait; (2) the evolutionary genomic analyses showed that seedless grapes were closely related with an origin from the well-known seedless grape ‘Sultania’ (TS). Introgression rather than convergent evolution was associated with the evolution of seedlessness in grapes; (3) a total of 110 QTLs associated with 634 candidate genes were identified through GWAS analysis within a large natural population, including four significant linkage regions such as Chr01: 18.49-19.96 Mb (specific to VV population), Chr07: 8.85-8.86 Mb (shared, SDI2 locus), Chr18: 11.7-20.0 Mb (specific to VV×VL population), and Chr18: 23.9-35.5 Mb (shared, SDInew locus); (4) a total of 339 core genes associated with seedlessness was detected through integrative genomics analyses; (5) machine learning based genome selection were built to accurately predict the seedless phenotypes. Importantly, these findings could efficiently save the cost and time in table grape breeding.

### The polygenic nature for seedlessness in grapes

The occurrence of seedlessness phenotypes from a cumulative polygenic effect associated with different tissues and development stages^17^. The limited observations employing single methods, such as transcriptomic analysis, are insufficient. As a result, numerous important candidate genes were ignored by previous studies. For example, *TT16/ABS*, found at the inversion boundary through comparative genomics (**Fig. 1d**), is one of promising candidate genes. Due to the redundant function between *STK* and *SHATTERPROOF* (*SHP1* and *SHP2*), the double *abs stk* mutants and triple *tt16 shp1 shp2* mutants all induced fewer seeds and exhibited defects in seed coat formation^17,50^. Candidate genes mapped by GWAS, such as *SMC1*, *11S globulin G1/2/3*, *LTG1*, *SUS2*, *LAC14*, *TT10/LAC15*, *MANNAN ENDO-1,4-BETA-MANNOSIDASE 5* (*MAN5*), *PROBABLE FRUCTOKINASE-5*, *E3 UBIQUITIN-PROTEIN LIGASE* (*DA2*), *AGL62*, etc., and well-known gene *VviAGL11*, play crucial roles in pollen, endosperm and seed coat development **(Fig. 3a, b and Supplementary Table 11**). Additionally, three transcriptomic analyses, GO homologous genes, and previously reported family genes provide us with numerous significant candidate genes (**Fig. 5a and Supplementary Table 11**).

Importantly, to define the interconnections among multiple genomic analyses, integrative genomic analysis was applied in this case, screened out 339 core genes with high significance from thousands of candidate genes (**Fig. 5a and Supplementary Table 17**). Most of these genes are associated with three main tissues related to seed coat, endosperm, and embryo, as shown in **Fig. 5b-d**, and ten other development processes (**Supplementary Table 11**). For example, we identified numerous candidate genes involving in seed hormone regulation, such as *MOTHER OF FT AND TFL1* (*MFT*) in ABA and GA pathways, *VvABI3-1*^68^ in ABA pathway, *VvGH3.9*^69^ in auxin pathways, and *VvMJE1*^64^ in jasmonate pathway. Additionally, genes controlling the development of floral organs, especially pollen and stigma, significantly influence the subsequent ovule development. This includes candidate genes like *CYP78A5*, *VvMADS27*, and metal ion transport genes *METALLOTHIONEIN-LIKE PROTEINS MT1* and *MT3*^70^. Except for lipid accumulation (**Fig. 5d**), the processes of sugar and amino acid synthesis also contribute to seed development, such as *BGLU15*, *VviGAPDH*^71^, glycine-rich protein (*Vitvi021557*, *Vitvi036421*), and 36.4 kDa proline-rich protein (*Vitvi002605*). Overall, these findings not only emphasize the polygenic nature of seedlessness but also provide novel candidate genes for functional genetics, highlighting the complexity of seed abortion regulation.

### The implications of genomic breeding of seedless table grapes

Both individual markers for Marker-Assisted Selection (MAS)^44,72–74^ and marker set for genomic selection generated in this study could be efficiently used in table grape breeding (**Fig. 6d**). Interestingly, in the variation map around *VviAGL11*, we observed that all the previously developed molecular markers based on lineages with a narrow genetic background for marker-assisted in table grape breeding were filtered away, due to either the low genotyping quality or a high missing rate, such as SCF27^75^, e7_VviAGL11^44^, 5U_VviAGL11^44^, P3_VviAGL11^18^, VMC7f2^61^, and VrSD10^76^ (**Fig. 4d**). Luckily, we designed a set of 12 markers, based on our GWAS analyses of natural population with species-wide genetic diversity. These markers could accurately delimit seeded and seedless grapes, achieving a precision rate >90% in nature populations (**Fig. 4e**).

Quantitative genetics analysis revealed 110 high-quality QTLs associated with seedlessness in grapevines, including 634 candidate genes (**Supplementary Table 11**). Through extracting GWAS significant variants from admixed population, we obtained detail information on all 794 significant variant sites for training models. Genome selections, employing machine learning algorithms, achieved an impressive precision of 99%. This approach could facilitate early genomic selection of natural germplasms and hybrid progeny. Many crops, such as tomato^77^, potato^78,79^, cereal^80^, rice^81,82^, wheat and maize^83,84^, have successfully applied genomic selection and prediction in their breeding programs. However, genomic selection has rarely been used on grape breeding. In the future, numerous agronomic traits like fruit aroma, disease resistance, soluble solids content, and so on, could be integrated into a single GS chip, offering a powerful genomic tool for genomic design of grapevine breeding.

## Methods

### Plant materials and genome sequencing

To enrich the genetic diversity of seedless grape varieties, we collected fresh tissues of two seedless grape varieties, ‘Thompson Seedless’ (TS) and ‘Black Monukka’ (BM), from the Anningqu Experimental Station (87°28′00″E, 45°56′00″ N) at the Xinjiang Academy of Agricultural Sciences in China. Genomic DNA was extracted from grape leaves using CTAB method, followed by purification with the QIAGEN Genomic kit (CAT#13343). For each sample, a total of 15 μg DNA was utilized for HiFi (SMRTbell) library preparation. The sheared DNA fragments (gTUBEs, Covaris, USA) underwent treatment with the SMRTbell Enzyme Cleanup Kit (Pacific Biosciences, CA, USA) and purification using AMPure PB Beads. The resulting libraries were employed for HiFi sequencing on a PacBio Sequel II instrument (CCS mode) with Sequencing Primer V2 and Sequel II Binding Kit 2.0 in Grandomics, yielding 59.78 Gbp and 60.35 Gbp of sequencing data for TS and BM, respectively.

For Hi-C library preparation, the fresh leaves were cut into 2 cm pieces and vacuum infiltrated with nuclei isolation buffer supplemented with 2% formaldehyde. The isolated nuclei were then digested with 100 units of restriction enzyme DpnII. The resulting Hi-C sequencing data amounted to 59.39 Gbp (TS) and 57.67 Gbp (BM) via the Illumina Novaseq/MGI-2000 platform. Additionally, the genomic DNA of 29 grape samples was extracted from fresh leaves, and the high-depth WGS sequencing was carried out using the PE150 mode of the Illumina Navaseq 6000 platform. For genome annotation, 1 μg total RNA was extracted from mixed tissues, such as roots, buds, and leaves. cDNA library was used TruSeq RNA Library Preparation Kit (Illuminia, USA) and sequenced with 150 bp pair-end reads on the Illuminia Navaseq 6000 platform.

### Genome assembly

The detailed workflow of haplotype-resolved genome assembly and annotation is described in **Extended Data Fig. 1**. The full pipeline for genome assembly and gap filling can be found on our lab GitHub@zhouyflab (see Code availability). In brief, we utilized the Hi-C and HiFi data integrated assembly algorithm to generate contig reads by Hifiasm (v. 0.16.1-r375)^85^. The contig reads were oriented and ordered to scaffold level using RagTag (v. 2.1.0)^86^ with default parameters. Hi-C reads were also employed to anchor scaffolds onto chromosomes by Juicer 2.0^87^ and Juicebox (v. 1.11.08)^88^. The adjusted genome at the chromosome level was generated using 3D-DNA (v. 201008)^89^. The detailed workflow of haplotype-resolved genome assembly and annotation is described in **Extended Data Fig. 1**. The full pipeline for genome assembly and gap filling can be found on our lab GitHub@zhouyflab (see Code availability). The chloroplast genome was de novo assembly using GetOrganelle toolkit (v1.7.7.0)^90^ with K-mer parameters set to 21, 65, 105, and 127. For the high-quality mitochondrial genome, de novo assembly was performed using Flye (v2.9.2)^91^ with HiFi long reads. Furthermore, using PN_T2T genome^46^, we utilized RagTag to assemble three chromosomes-level genomes based on scaffold level: ‘Cabernet Sauvignon’ (CS)^52^, ‘Black Corinth Seedless’ (BC seedless)^92^, and ‘Black Corinth Seeded’ (BC seeded)^92^.

### Assembly assessment

HiFi data from TS and BM were performed to assess genome heterozygosity based on k-mer using GenomeScope2.0^93^. Meanwhile, basic genome statistics were calculated using seqkit (v. 2.2.0)^94^, which included genome length, N50, and GC content. The completeness of the haplotype genomes was evaluated using BUSCOs (v. 5.3.0)^95^ with the embryophyta_odb10 database. Merqury (v. 1.3, best_k=19)^96^ was employed to evaluate the quality value (QV) and completeness of the haplotype genomes based on whole-genome sequencing data. The Hi-C interactive signals on grape genome were visualized using Juicebox.

### Genome annotation

The genome-wide annotation pipeline, identification of telomeres and centromeres, and annotation of transposable elements (TEs) were referred from pervious study^46,97^. The statistic results of TEs classification through Pan-genome TE annotation^46^ can be found in **Supplementary Table 2**. Additional details on telomere regions, telomere copy numbers, and centromere regions for each chromosome are provided in **Supplementary Table 3**. The CS genome was annotated based on sequence similarity using Liftoff (v. 1.6.3)^98^ and PN_T2T annotation files.

### Comparative genomics

For genome level variant calling, we selected a total of 15 grape genomes, including ten seeded accessions: ‘PN40024’ (PN_T2T), ‘Cabernet Sauvignon’ (CS, hap1 and hap2)^92^, ‘Muscat Hamburg’ (MH, hap1 and hap2), ‘Shine Muscat’ (SM, hap1 and hap2), ‘Muscadinia rotundifolia’ (MR, hap1 and hap2) ^99^, BC Seeded^92^, as well as five seedless: TS (hap1 and hap2), BM (hap1 and hap2), BC Seedless^92^. Among them, TS and BM were newly sequenced in this study. Four recently synchronized haplotype-resolved genomes of MH and SM will be published in another study.

All chromosome-level genomes were aligned with PN_T2T genome using Mummer4 (v. 4.0.0rc1)^100^, and the results were visualized using Plotsr (v. 0.5.4)^101^ and Linux based Gunplot. R script was used for visualizing the gene density (line) of the reference genome (see Code availability). To validate the authenticity of SVs, the raw reads of HiFi and Hi-C were mapped to their haplotype genomes using Minimap2. The corresponding BAM files were extracted using SAMtools (v. 1.13)^102^ and then inputted into the IGV software for inversions validation. In addition, the genome consensus phylogenetic tree was constructed using OrthoFinder (v. 2.5.4)^103^ based on the single-copy orthologous genes from the whole genomes.

### WGS variation detection

To genotype SNPs, InDels and SVs in 548 accessions, low-quality resequencing reads were removed using Fastp (v. 0.23.2)^104^ with default parameters. The filtered data were then mapped to the CS reference genome using BWA (v. 0.7.17-r1188)^105^. Non-uniquely mapped and duplicated reads were excluded using SAMtools and GATK (v.4.2.3.0)^106^. Subsequently, SNPs and InDels calling were performed using GTX (v. 2.1.11, http://www.gtxlab.com/product/cat), followed by the merging of genotyping of the gVCF files into a VCF file. Delly (v. 1.1.6)^107^ was used to SVs calling with default parameters. Basic filtering of the VCF file was performed using VCFtools (v. 0.1.16)^108^ with the following parameters: --max-missing 0.8, --minGQ 20, --min-alleles 2, --max-alleles 2, --minDP 4, --maxDP 1000, and --maf 0.0005. PLINK (v. 1.90b6.21)^109^ was utilized to reduce the size of the VCF file and improve computational efficiency. Finally, we obtained a total of 4,462,797 SNPs, 443,812 InDels, and 487,204 SVs from 548 grape accessions in the nuclear genome, 38,267 variation sites (SNPs and InDels) from 314 grape accessions in the MT genome, and 2,247 variation sites (SNPs and InDels) from 314 grape accessions in the Pltd genome.

### Population genetics analysis

Phylogenetic analysis was conducted using 250,821 high-quality SNPs filtered for LD by PLINK, with parameters: --indep-pairwise 20 5 0.2 --geno 0.1. The phylogenetic tree was inferred by Iqtree (v. 2.1.4-beta)^110^ with parameters: -m GTR+I+G -bb 1000 -bnni -alrt 1000 -st DNA, and the result were visualized using iTOLs^111^. Similarly, the phylogenetic analysis of the MT and PT genomes yielded 9,647 and 758 high-quality SNPs using PLINK (--geno 0.2 --maf 0.001), respectively. The PCA and IBD analysis were preformed using PLINK and visualized using R scripts and Cytoscape (v. 3.9.1)^112^, respectively. VCFtools was employed to calculate the *F*_ST_ and π statistics at whole genome level with 20 kb window size. Population introgression analysis was calculated using the "ABBABABAwindows.py" script from the "general_genomics" tool (https://github.com/simonhmartin/genomics_general) with 20 kb window size. The population branch statistic (PBS) assesses differentiation in the same branch of the phylogenetic tree with 50 SNP per window between seeded and seedless grapes in both VV and VV×VL populations^113^. PBScan was utilized to estimate population differentiation using *D_xy_* (-div 2).

### Genome-wide association study

We selected three populations for GWAS analysis, including VV population (n = 352), VV×VL population (n = 92), and the admixed population (n = 444). High-quality variants were obtained using PLINK with MAF ≥ 0.05 and missing < 0.2, resulting in 2,086,600 variants (1,881,457 SNPs, 173,547 InDels, and 31,596 SVs), 1,419,982 variants (1,274,130 SNPs, 124,054 InDels, and 21,798 SVs), and 2,292,404 variants (2,065,307 SNPs, 192,536 InDels, and 34,561 SVs), respectively. GWAS analysis utilized the mixed linear model in GEMMA (v. 0.98.3)^114^ with the first three PCAs as a random effect matrix. Variant positions and p-wald test statistics (*P-*value) were extracted to generate Manhattan plots and Q-Q plots by R scripts. A Python script was utilized to identify the associated genes within 5 kb windows of the significant variants, which were above the significance threshold of −log_10_(0.05/Variant Numbers).

GO enrichment analysis was employed using the web tool DAVID^115^, and the results were visualized with R script. The SDI locus was identified through molecular markers SNP-25.24 and SNP-26.93^20^. The LD linkage heatmap was visualized using LDBlockshow (v. 1.40)^116^. Moreover, DIAMOND (v. 2.0.15)^117^ was employed for sequence comparisons of whole-genome homologous protein from 14 genomes (Seeded: Seedless: Outgroup = 8:5:1). Sequence alignments of candidate proteins were conducted using ClustalW in MEGA11 ^118^, and sequence visualization was accomplished using GeneDoc (v. gd322700)^119^. The visualization of FPKM values, gene structures, and mutation ratios was achieved using R scripts (see Code availability).

### Multi-transcriptome analysis

Three independent transcriptome datasets were downloaded from the NCBI database (see **Supplementary Table 15**). The 76 samples encompassed six grape varieties: PN and TS, which underwent repeated testing from 20 DAF to 50 DAF over a two-year period; ‘Italia’ and HS, continuously evaluated from 7 to 42 DAF within a single year; and Himrod (VV×VL population) and Jinzao (VV population), continuously monitored from full flowering stage to grape maturity within a single year. These raw fastq data underwent quality control and data cleaning using Fastp with default parameters. Using the PN_T2T genome as a reference, transcriptomic assembly was conducted on the processed data using STAR (v. 1.5.2)^120^. The R package DESeq2 (v.1.36.0)^121^ was utilized for PAC analysis and pairwise comparisons between seeded and seedless grape samples at each development time point. The thresholds for differential expression genes were |Log2FoldChange| ≥ 2 and P-adjust < 0.05.

### Integrative genomic analysis

Homologous protein alignment identified 14,650 genes within the 34 seed development-related GO terms (EMBL-EBI, QuickGO database, 2023-01-03), and 6,529 genes with high sequence similarity were screened (identity ≥ 50%, e-value ≤ 1e-5). Additionally, the primer sequences collected from previous studies in the past decade, including 451 genes and 7 molecular markers associated with seed abortion processes. All of them were then mapped to the PN_T2T genome based on sequence alignment using BALST in TBtools (v. 1_113)^122^. The results were visualized using Venn diagrams and whole-genome density plots using the R package ggvenn (v0.1.10) and RIdeogram^123^, respectively. The integrated results in this study, including GWAS results, RNA-seq results, GO enrichment results, and previously reported family genes, were overlapped and filtered manually. The core candidate genes were screened based on an average expression value across all time points (AVE FPKM ≥ 100) and Fold-Changes (Fold = Seeded AVE FPKM / Seedless AVE FPKM, Fold ≤ 0.5 or Fold ≥ 2.0). Data visualization was performed using an R script, which included UpSet plot analysis and gene expression heatmaps (see Code availability).

### Genome selection based on machine learning

For genome selection, we utilized the admixed population (n = 444) as training set, and additional 39 samples as testing set for phenotype prediction, as pipeline showed in **Extended Data Fig. 14**. Beagle (v. 5.2_21Apr21.304.jar)^124^ was used to impute the VCF file. Subsequently, the 794 variants, above the significance threshold of −log_10_(0.05/Variant Numbers) (∼ 7.66), were extracted from imputed VCF using BCFtools (v. 1.13)^125^, including 717 SNPs and 77 InDels. For model selection, we utilized the Python package ‘sklearn’ (https://scikit-learn.org/stable/install.html) and selected nine classical models: Cross-validated Elastic Net model (ElasticNetCV), Kernel Ridge Regression (KernelRidge), Lasso Regression (Lasso), Linear Regression (Linear), Logistic Regression (Logistic), PLS Regression, Linear Ridge Regression (Ridge), Linear Support Vector Regression (SVR-Linear), and Polynomial Support Vector Regression (SVR-poly). After 100 rounds of random cross-validation with the training set, we chose the models with best performance, SVR-poly and ElasticNetCV, for the prediction of testing set. Moreover, the Iqtree was employed to construct an unrooted tree, while iTOLs was used for the visualization of the phylogenetic tree. The data visualization used R scripts.

## Supplementary Data

**Extended Data Fig. 1 | Complete workflow for haplotype-resolved genome assembly and annotation.** Additional details can be found on our lab website GitHub@zhouyflab.

**Extended Data Fig. 2 | Evaluation of four haplotype-resolved genomes**. **a**, BUSCO assessment of genome completeness using the embryophyta_odb10 database. **b**, Evaluation of quality value (QV) and haplotype completeness using Merqury based on k-mer. **c**, Visualization of the diploid genome via Hi-C heatmap using Juicebox. Most single contig (green rectangle) are directly composed of chromosomes (blue rectangles).

**Extended Data Fig. 3 | Comparative genomics results. a**, Sequence alignment of 15 grape genomes with the PN_T2T genome. Red blocks represent seedless samples, green blocks indicate seeded samples, and gray blocks denote outgroup samples. Red stars highlight inversions associated with seed abortion. The Chr07 of ‘*Muscadinia rotundifolia*’ is composed of Chr07 and Chr20. **b**, Sequence alignment results of Chr15 for the 15 genomes, as well as the inversion Hi-C heatmap in inversion boundary. The phylogenetic tree was constructed using single-copy genes from the whole genome proteins.

**Extended Data Fig. 4 | Reads mapping at the inversion breakpoints in seedless haplotype genome. a**, **c**, and **e**, represent the start points of the inversions, while **b**, **d**, and **f**, represent the end points of the inversions. Coverage depth is halved before and after the breakpoint junctions, revealing transitions between heterozygous and homozygous states of reads sequences are observed.

**Extended Data Fig. 5 | Detailed phylogenetic tree of 548 grapevine accessions.** This figure provides a full zoomed-in version of Fig. 3c, which includes the six populations. Light-blue blocks represent seed abortion samples and black star symbols indicate TS and BM.

**Extended Data Fig. 6 | Phylogenetic tree of mitochondrial and chloroplast genomes in 314 grapevine accessions. a-c**, represent the consensus phylogenies constructed based on the mitochondrial genomes, nuclear genomes, and chloroplast genomes, respectively. **d-e**, depict the complete phylogenetic trees of the mitochondrial and chloroplast, encompassing six populations.

**Extended Data Fig. 7 | Visualization of whole-genome analyses.** From top to bottom, these results included QTL peaks (red: VV, purple: VV × VL, black: consensus peaks), GWAS analyses within three populations (red: VV, purple: VV×VL, dark-blue and yellow: admixed population; points: SNPs, triangles: InDels and SVs), population analyses of fixation indices (*F*_ST_), nucleotide diversity (π), introgression (*f*_d_), and divergent selection (PBS) (refer to Supplementary Table 13), and 339 core candidate genes (refer to Supplementary Table 11).

**Extended Data Fig. 8 | Quantile-Quantile (Q-Q) plot and Gene Ontology (GO) enrichment analysis.** Genome-Wide Association Study (GWAS) Q-Q plot for the three populations, along with GO enrichment analysis for biological processes in genes specific to VV×VL and VV, as well as those consensus gene for admixed populations (refer to Supplementary Table 12).

**Extended Data Fig. 9 | Sequence alignment of *11S globulin G1*-*G3* homologous genes from 14 grape genomes:** includes ‘PN_T2T’ (PN40024), ‘Cabernet Sauvignon’ (CS), ‘Muscat Hamburg’ (MH), ‘Shine Muscat’ (SM), ‘Black Corinth Seeded’ (BCsd), ‘Black Corinth Seedless’ (BCsl), ‘Thompson Seedless’ (TS), ‘Black Monukka’ (BM), and ‘Muscadinia Rotundifolia’.

**Extended Data Fig. 10 | Three candidate genes related to seedlessness in Chr18. a-c**, Candidate genes were identified through GWAS analysis using the admixed population, with a significant threshold of 7.61 for SNPs and 6.66 for SVs (and InDels).

**Extended Data Fig. 11 | Principal Component Analysis (PCA) analysis of the three transcriptomic datasets, and relative expression values (FPKM) for *VviAGL11*.**

**Extended Data Fig. 12 | Enrichment analysis of GO homologous genes related to grape seed development. a**, Enrichment analysis of 14,650 seed development genes from the GO database. **b**, Enrichment analysis of 6,529 GO homologous genes associated with grape seed development.

**Extended Data Fig. 13 | Visualization of multiple seed development-related datasets. a** and **b**, Results from two independent transcriptomic analyses, ‘Italia’ vs ‘Hongju Seedless’ (HS) and ‘Pinot Noir’ (PN) vs ‘Thompson Seedless’ (TS), overlapping with GO homologous gene. Red indicates up-regulated DEGs, blue represents down-regulated DEGs, and yellow denotes GO homologous genes associated with seedlessness. **c**, Results from integrative genomic analyses: GWAS, transcriptomics, reported genes mapping, and GO homologous genes. Green bars represent the total number of genes identified through this approach. **d**, Visualization of three datasets: GO homologous genes, reported gene families, and reported molecular markers.

**Extended Data Fig. 14 | Genome selection workflow.** The 794 significant variants extracted from GWAS results, including 77 InDels and 717 SNPs. More detail code can be found on our lab GitHub@zhouyflab.

**Supplementary Table 1. Assessment of genome quality.** A comparison between four T2T haplotype-resolved genomes and the completed reference genome PN_T2T.

**Supplementary Table 2. Pan-genome TE annotation results in the four haplotype-resolved genomes.**

**Supplementary Table 3. Centromere and telomere regions in the four haplotype-resolved genomes.** This information includes the start position, end position, copy numbers, and TRF ID.

**Supplementary Table 4. Comparative genomic statistics.** Genome alignment was conducted using the SyRI (v. 1.5.4) with input from the output file generated by Mummer4 (v. 4.0.0rc1).

**Supplementary Table 5. Summary of genome annotation in the TS hap2 genome region Chr15: 8.72-9.90 Mb.** This region contains a total of 111 genes, including 73 shared genes and 38 genes that are gained in the TS hap1 genome.

**Supplementary Table 6. Summary of genome annotation in the TS hap1 genome region Chr10: 21.75-26.00 Mb.** This region contains a total of 210 genes, including 79 shared genes and 131 genes that are exclusively lost in the TS hap2 genome.

**Supplementary Table 7. Summary of genome annotation in the BM hap1 genome region Chr10: 23.00-27.50 Mb.** This region contains a total of 237 genes, including 69 shared genes and 168 genes that are exclusively lost in the BM hap2 genome.

**Supplementary Table 8. Population information for grape resequencing.** This table provides details, such as NCBI accessions, the full name of sample, species, seed conditions, population used in GWAS, samples used in phylogenetic trees, samples for population analyses (*F*_ST_, π, *f*_d_, and PBS), the training set and testing set for genome selection, as well as prediction results of best-performance models.

**Supplementary Table 9. Identity by Descent (IBD) matrix for 46 seedless individuals.** In this matrix, the VV population is denoted by purple, while VV×VL population is indicated by red. Detailed information for all samples used can be found in Supplementary Table 8.

**Supplementary Table 10. GWAS results of three populations.** This table includes the variant positions, allele changes, −Log10 (*P* value), and associated genes (within ± 5 kb of the variant sites).

**Supplementary Table 11. Integrative genomic analysis across all study results.** This table consolidates GWAS analysis results in different populations, differentially expressed genes from three transcriptomic analyses, grape seed development associated GO homologous genes, reported gene families and molecular markers, as well as 339 core candidate genes identified through integrative genomic analysis. The reference genome utilized ‘Cabernet Sauvignon’ (CS), and the homologous proteins was aligned with the UniProt database and PN40024 12X (GCF_000003745.3), respectively.

**Supplementary Table 12. Distinct and consensus genes, and GO enrichment analyses in three populations.** This table presents candidate genes specific to the VV population, those specific to VV×VL population, and those consensus in the admixed population as identified through GWAS. GO enrichment analysis was conducted using the online toolkit DAVID (https://david.ncifcrf.gov/tools.jsp).

**Supplementary Table 13. Population analyses results.** This table includes the genome-wide fixation indices (*F*_ST_) analysis for 11 VV and 35 VV×VL seedless samples, genetic diversity (π) analysis for two population (11 VV and 35 VV×VL seedless samples), *f*_d_ statistics analysis for gene introgression, and population branch statistic (PBS) analysis for VV and VV×VL populations. Samples were selected from a branch in the phylogenetic tree, with wild grapes (ME) as the outgroup. The statistical window size is 20 kb for *F*_ST_, π and *f*_d_, while 50 SNPs per window size for PBS analysis. Population information can be found in Supplementary Table 8.

**Supplementary Table 14. Mutation ratio statistics of significant variants identified from GWAS analysis within Chr18:30.70-31.32 Mb in the admixed population.**

**Supplementary Table 15. Information on three transcriptomic groups, including NCBI accessions, full name of samples, time points, species and so on.**

**Supplementary Table 16. Primer sequences and data sources for 451 family genes and 7 molecular markers, mapping proteins with the PN_T2T genome.**

**Supplementary Table 17. Overlapped genes statistics used for upset plot.** Further gene details can be found in Supplementary Table 11.

**Supplementary Table 18. Genotyping information of 794 high-quality variants (77 InDels and 717 SNPs).** This table also includes prediction results using genome selection based on the SVR-poly model. ‘0’ represents 0/0, ‘1’ represents 0/1, and ‘2’ represents 1/1.

## Acknowledgements

We express our gratitude to Zhiwu Zhang of Washington State University, USA, for providing invaluable suggestions regarding genome selection and prediction. We thank members of the Zhou lab at Agricultural Genomics Institute at Shenzhen (AGIS) for discussions and comments on the project. We thank Lin Cheng, Hongbo Li, Zhigui Bao, and Qing Zhang at AGIS for giving guidance and scripts to haplotype genome assembly.

## Author contributions

Y. Z. conceived and designed the study. H. Z., F. Z. and X. W. collected the plant materials. H. Z. and Z. L. performed experiments and genome sequencing. X. W., S. C., Y. S., T. H., and Y. Z assembled and annotated the haplotype-resolved T2T genomes. X. W., Z. L., F. Z., X. H., W. L., Z. L., and Y. G. performed data analysis. Y. Z., X. W., Z. L., F. Z., H. X. wrote the original draft of the manuscript. All authors provided critical feedback and revised the manuscript. X. W. and Z. L. contributed equally to this work.

## Data availability

The raw sequencing data, comprising PacBio HiFi long-reads, Illumina Hi-C reads, RNA-seq reads, and 29 WGS grape accessions, is accessible on NCBI under BioProject ID PRJNA1021353 and on the National Genomics Data Center (NGDC) under BioProject ID PRJCA022010. The genome assembly and their annotations have been deposited into in Zenodo: https://doi.org/10.5281/zenodo.8278185.

## Code availability

All scripts performed in this study are available on GitHub: https://github.com/zhouyflab/Polygenetic_Basis_Seedless_Grapes

