## Extended Data Fig. 1-14 for "Integrative genomics reveals the polygenic basis of seedlessness in grapevine"

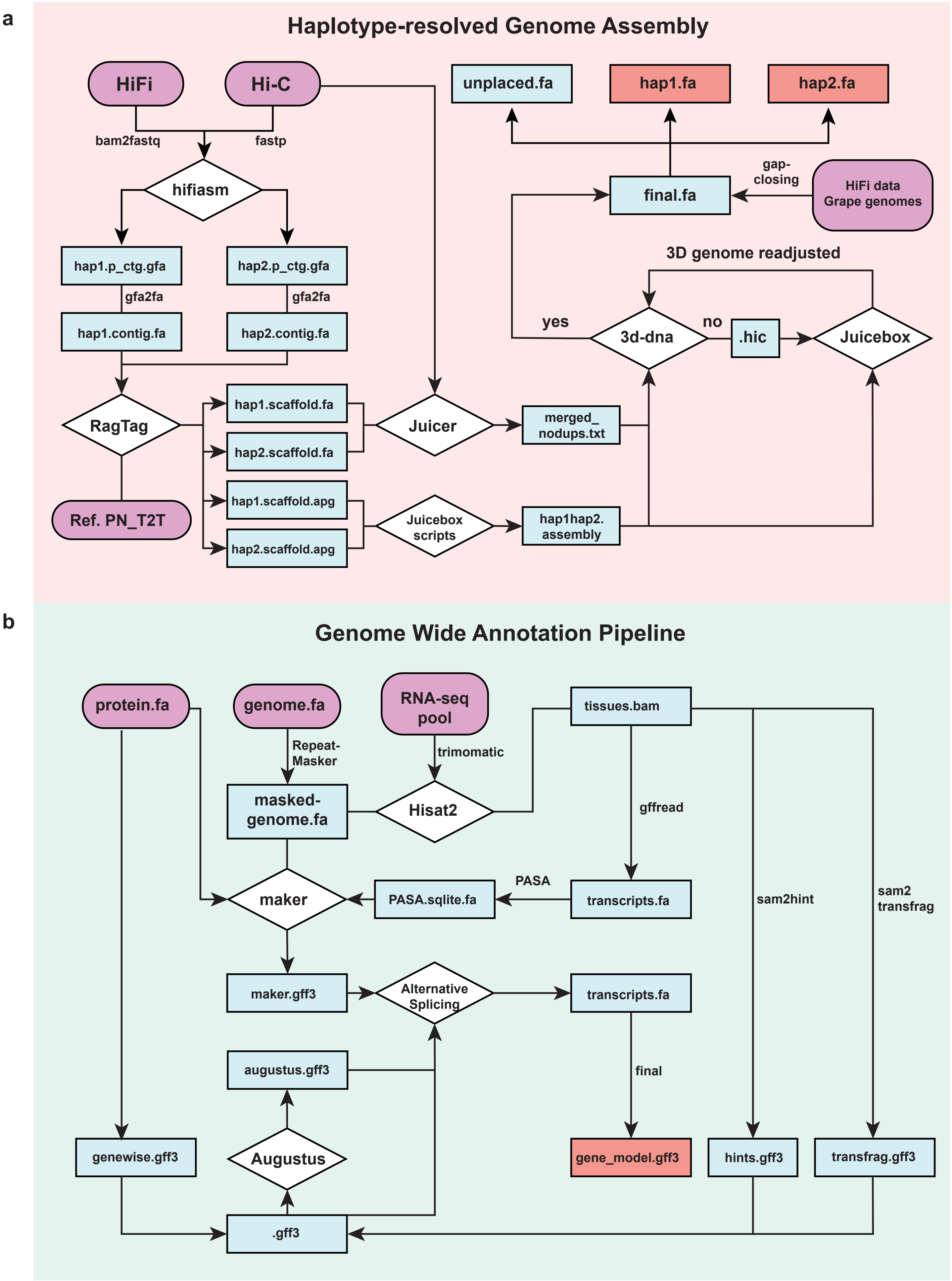


**Extended Data Fig. 1 | Complete workflow for haplotype-resolved genome assembly and annotation.** Additional details can be found on our lab website GitHub@zhouyflab.

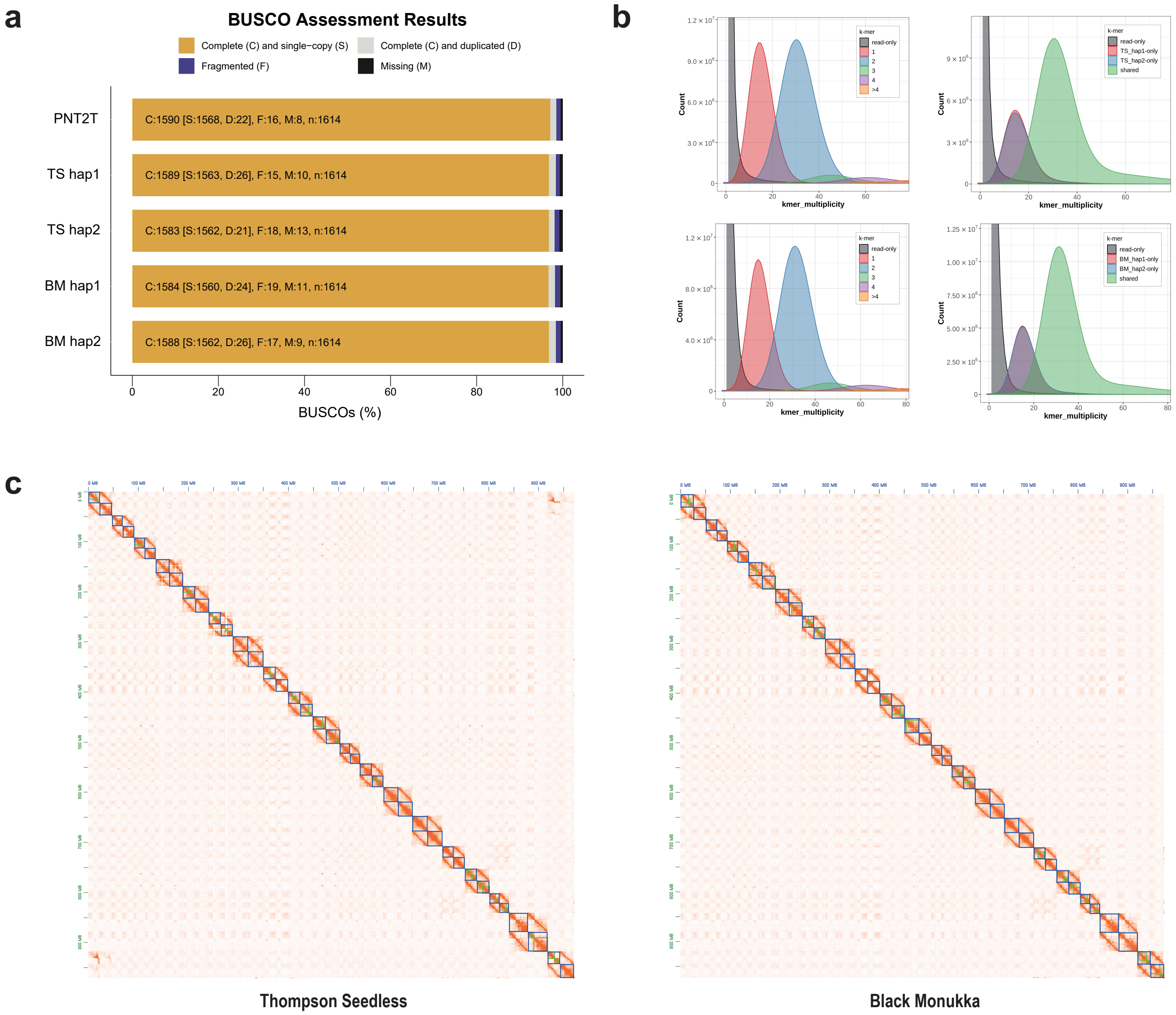


**Extended Data Fig. 2 | Evaluation of four haplotype-resolved genomes. a**, BUSCO assessment of genome completeness using the embryophyta_odb10 database. **b**, Evaluation of quality value (QV) and haplotype completeness using Merqury based on k-mer. **c**, Visualization of the diploid genome via Hi-C heatmap using Juicebox. Most single contig (green rectangle) are directly composed of chromosomes (blue rectangles).

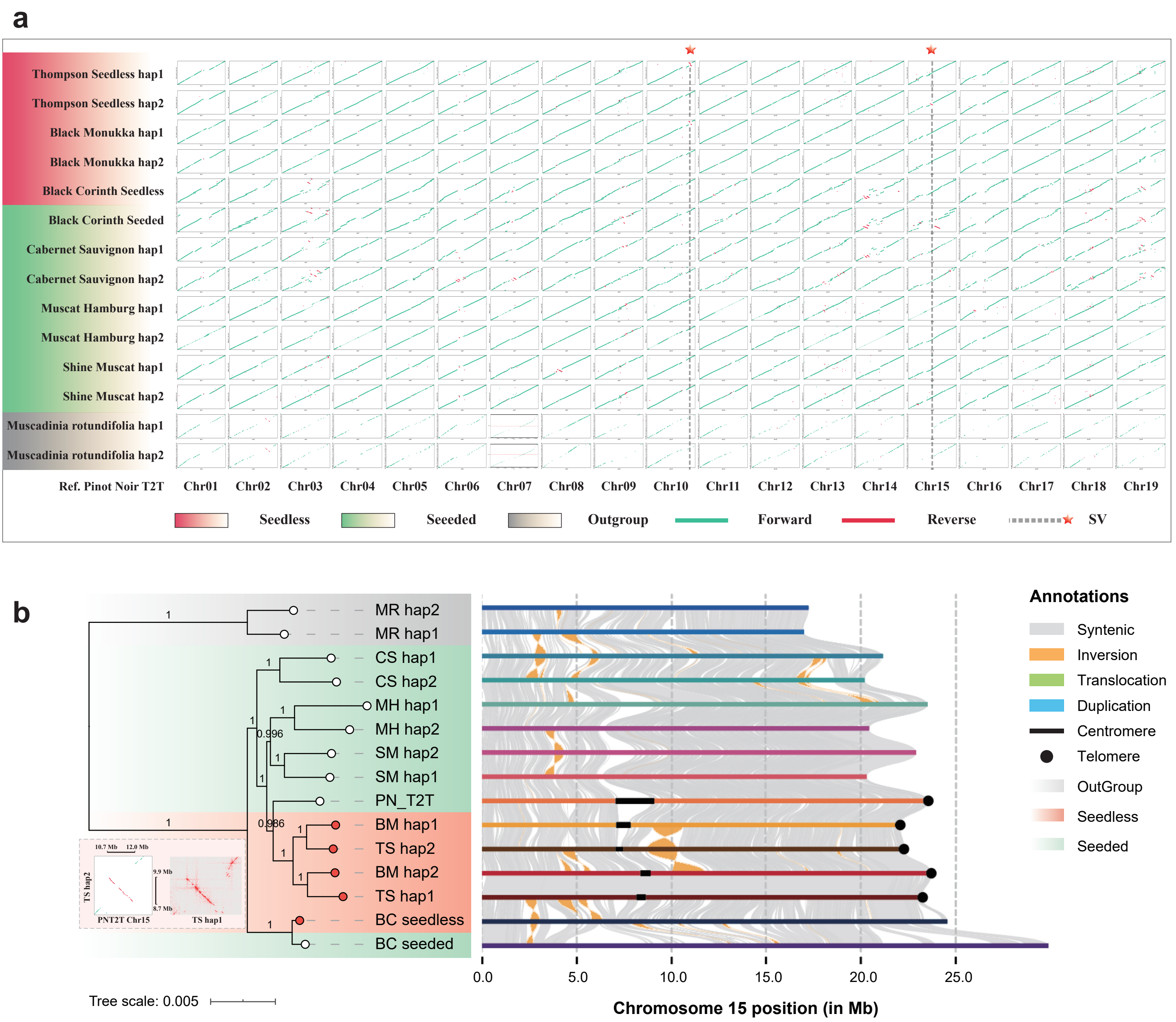


**Extended Data Fig. 3 | Comparative genomics results. a**, Sequence alignment of 15 grape genomes with the PN_T2T genome. Red blocks represent seedless samples, green blocks indicate seeded samples, and gray blocks denote outgroup samples. Red stars highlight inversions associated with seed abortion. The Chr07 of '*Muscadinia rotundifolia*' is composed of Chr07 and Chr20. **b** Sequence alignment results of Chr15 for the 15 genomes, as well as the inversion Hi-C heatmap in inversion boundary. The phylogenetic tree was constructed using single-copy genes from the whole genome proteins.

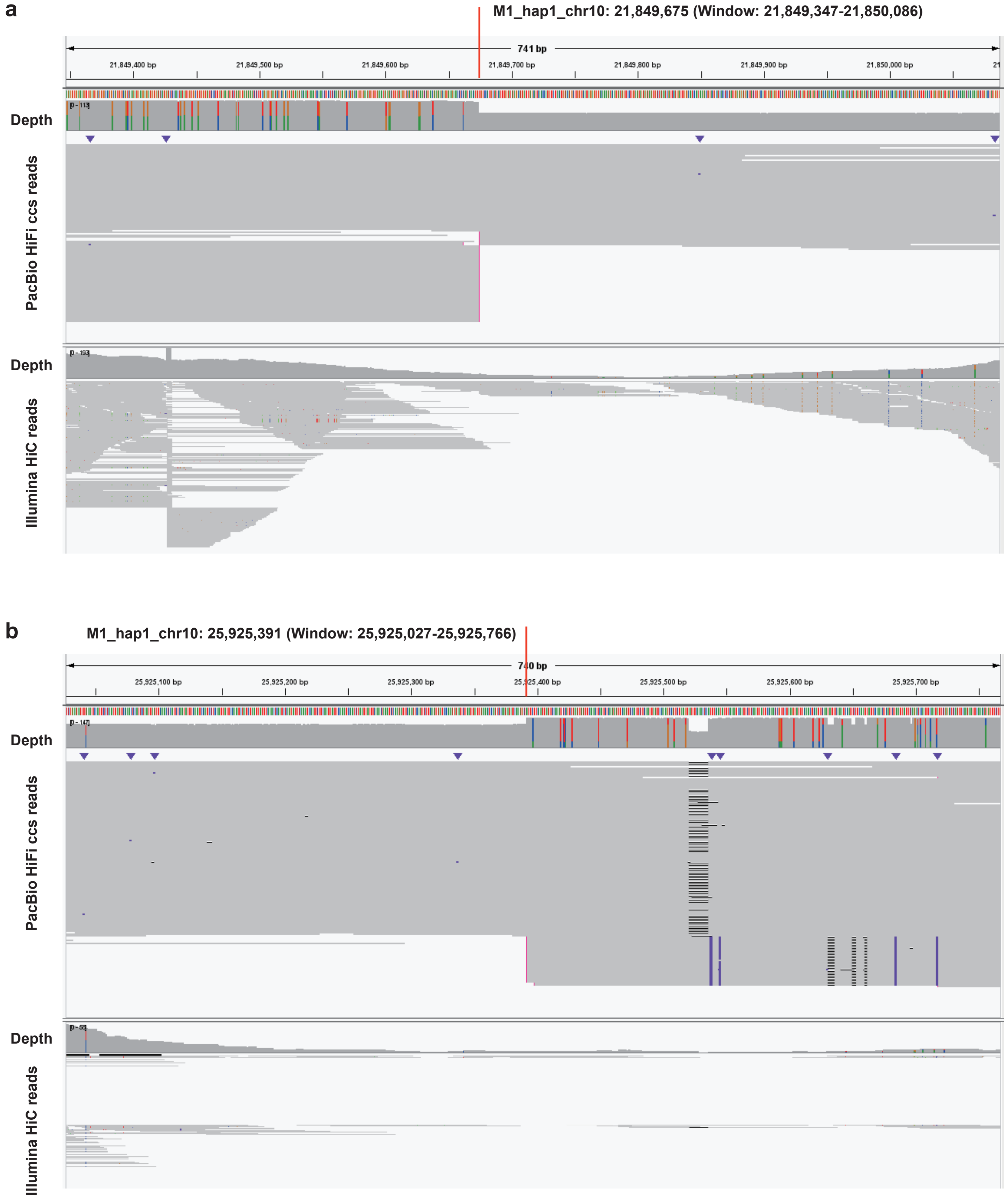
a
M1_hap1_chr10: 21,849,675 (Window: 21,849,347-21,850,086)
741 bp
21,849,400 bp
21,849,500 bp
21,849,600 bp
21,849,700 bp
21,849,800 bp
21,849,900 bp
21,850,000 bp
Depth
PacBio HiFi ccs reads
Depth
Illumina HiC reads
b
M1_hap1_chr10: 25,925,391 (Window: 25,925,027-25,925,766)
740 bp
25,925,100 bp
25,925,200 bp
25,925,300 bp
25,925,400 bp
25,925,500 bp
25,925,600 bp
25,925,700 bp
Depth
PacBio HiFi ccs reads
Depth
Illumina HiC reads

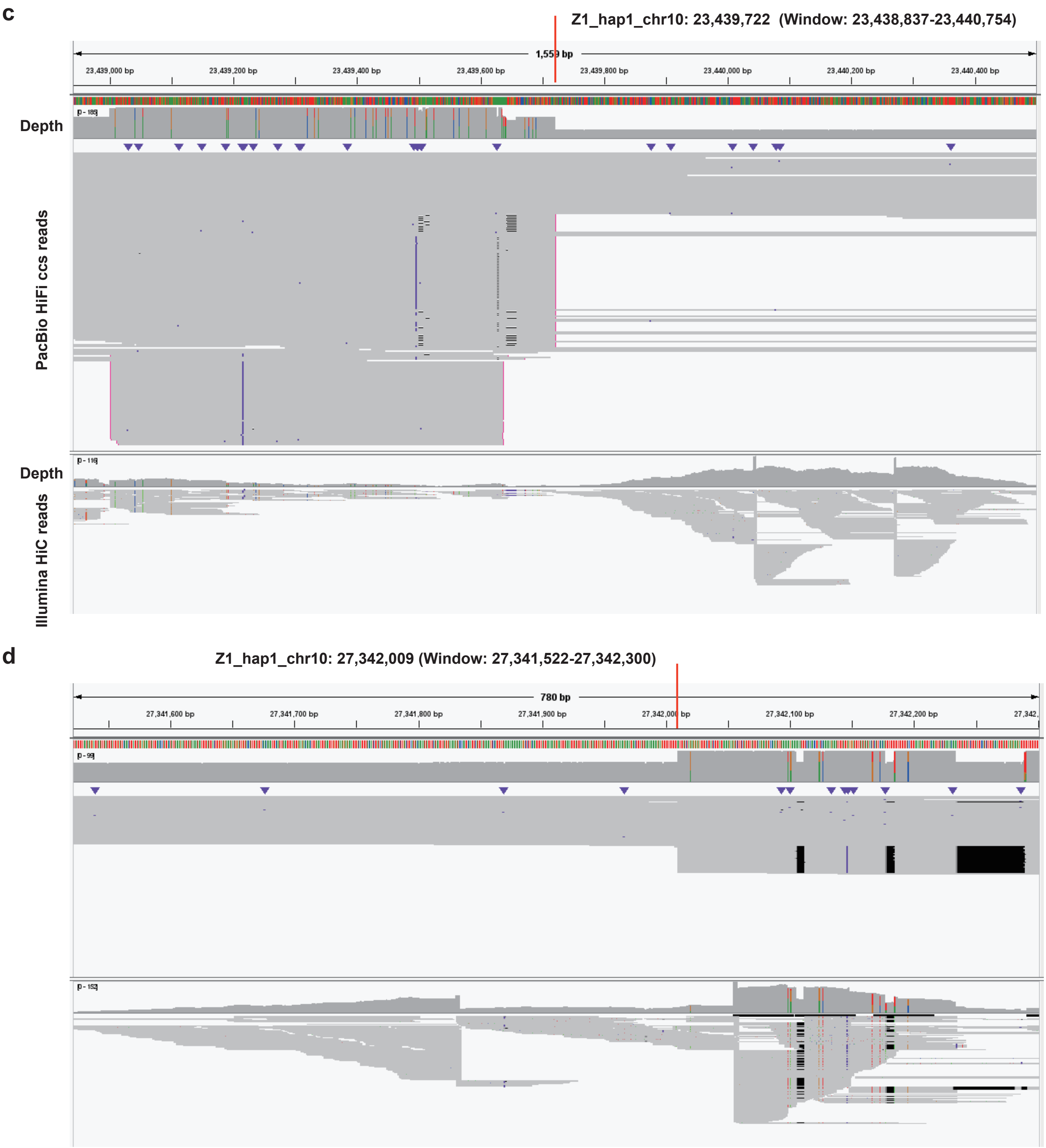
c
Z1_hap1_chr10: 23,439,722 (Window: 23,438,837-23,440,754)
1,559 bp
23,439,000 bp
23,439,200 bp
23,439,400 bp
23,439,600 bp
23,439,800 bp
23,440,000 bp
23,440,200 bp
23,440,400 bp
Depth
PacBio HiFi ccs reads
Depth
Illumina HiC reads
d
Z1_hap1_chr10: 27,342,009 (Window: 27,341,522-27,342,300)
780 bp
27,341,600 bp
27,341,700 bp
27,341,800 bp
27,341,900 bp
27,342,100 bp
27,342,200 bp

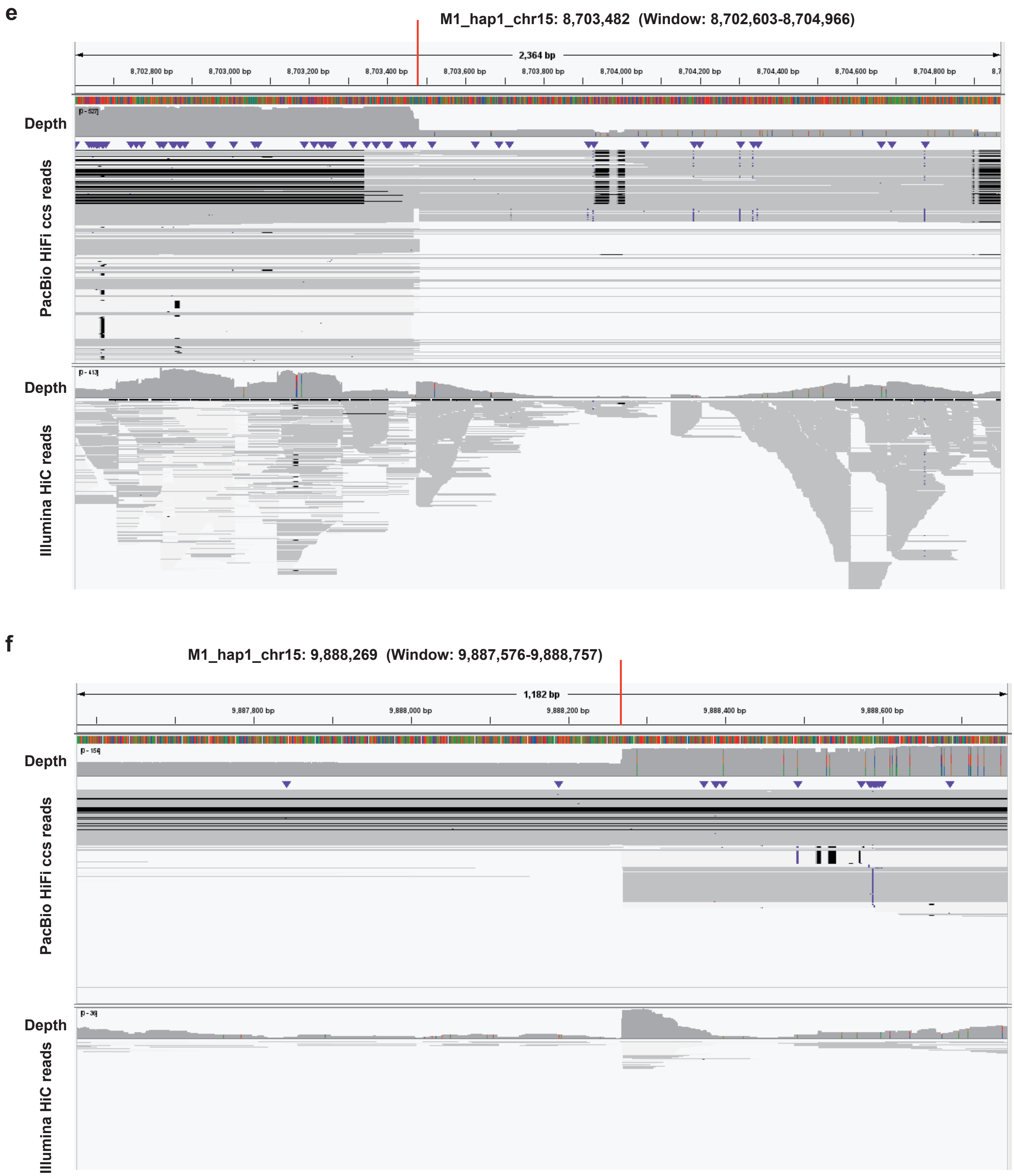


**Extended Data Fig. 4 | Reads mapping at the inversion breakpoints in seedless haplotype genome.** **a**, **c**, and **e**, represent the start points of the inversions, while **b**, **d**, and **f**, represent the end points of the inversions. Coverage depth is halved before and after the breakpoint junctions, revealing transitions between heterozygous and homozygous states of reads sequences are observed.

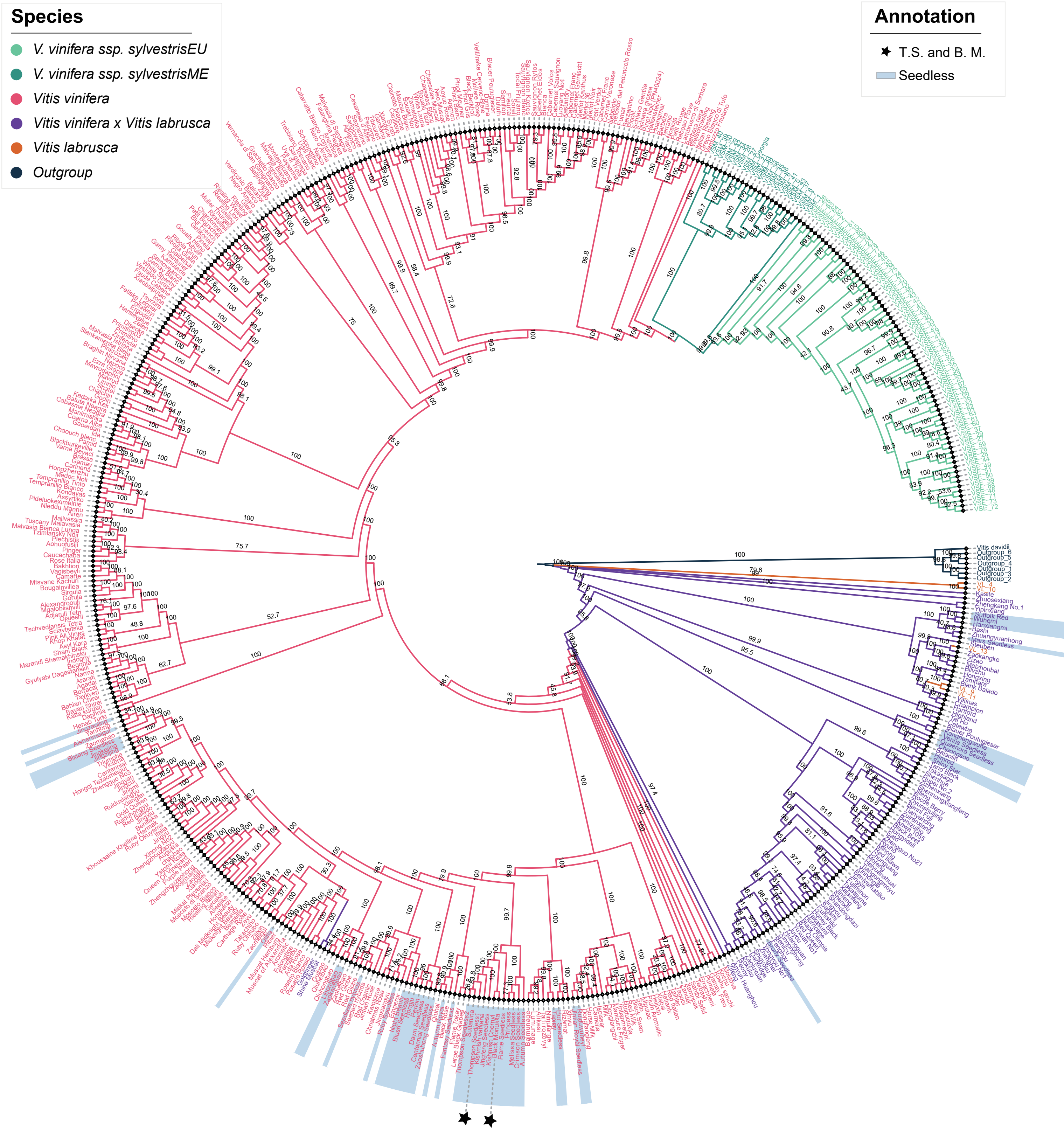


**Extended Data Fig. 5 | Detailed phylogenetic tree of 548 grapevine accessions.** This figure provides a full zoomed-in version of **Fig. 3c**, which includes the six populations. Light-blue blocks represent seed abortion samples and black star symbols indicate TS and BM.

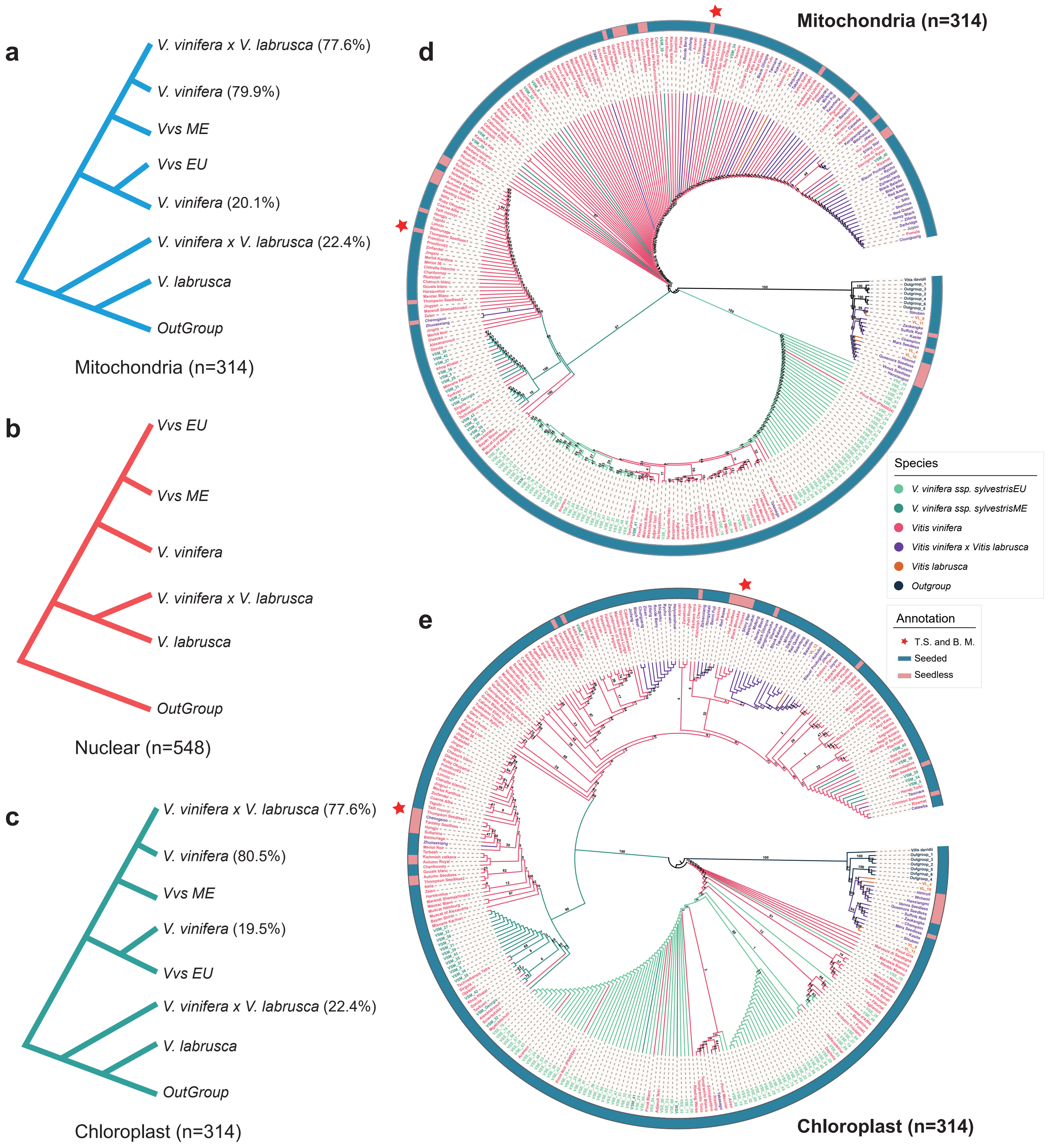


**Extended Data Fig. 6 | Phylogenetic tree of mitochondrial and chloroplast genomes in 314 grapevine accessions. a-c**, represent the consensus phylogenies constructed based on the mitochondrial genomes, nuclear genomes, and chloroplast genomes, respectively. **d-e,** depict the complete phylogenetic trees of the mitochondrial and chloroplast, encompassing six populations.

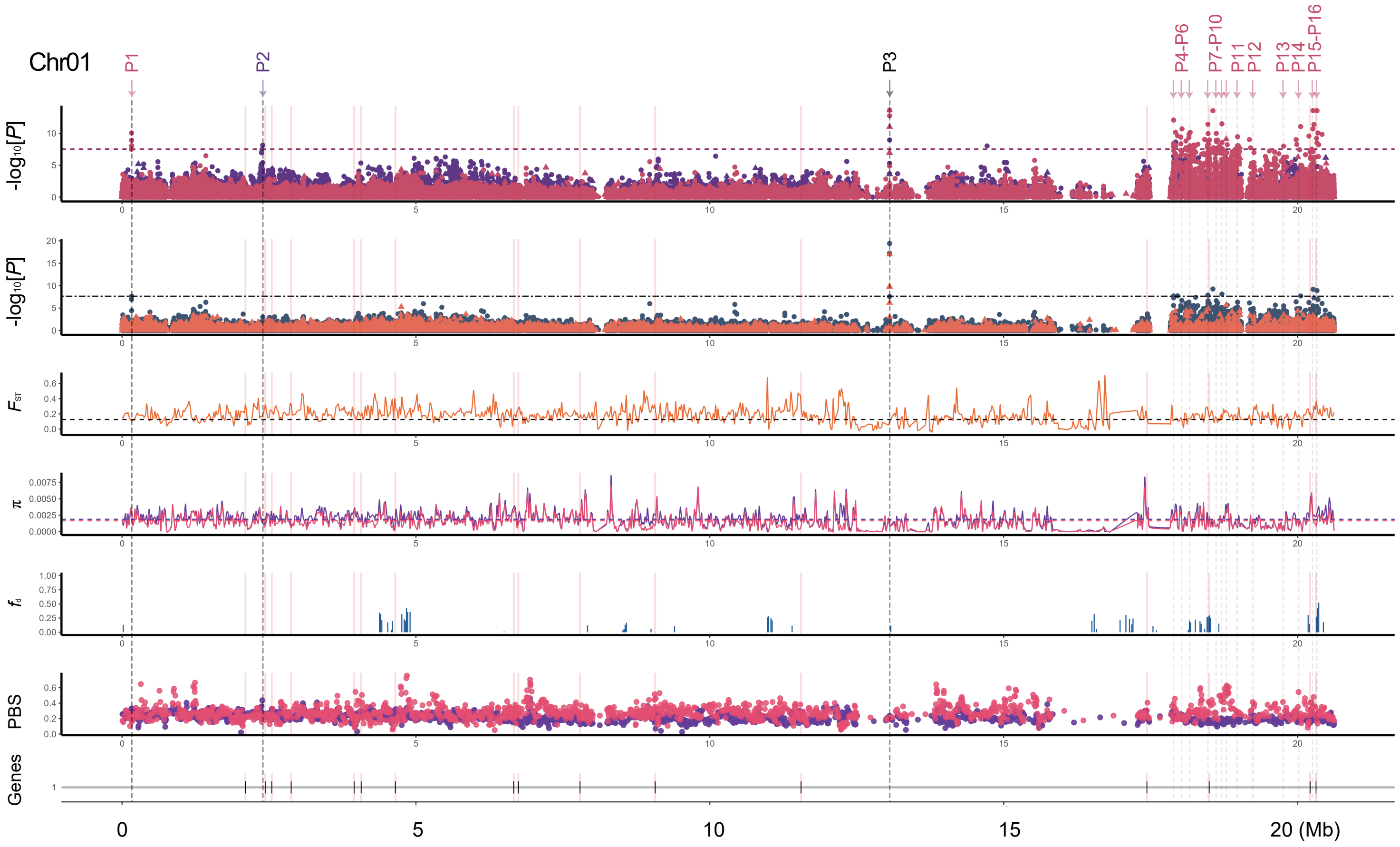
Chr01
P1
P2
P3
P4-P6
P7-P10
P11
P12
P13
P14
P15-P16
-log10[P]
FST
π
fd
PBS
Genes
0
5
10
15
20 (Mb)

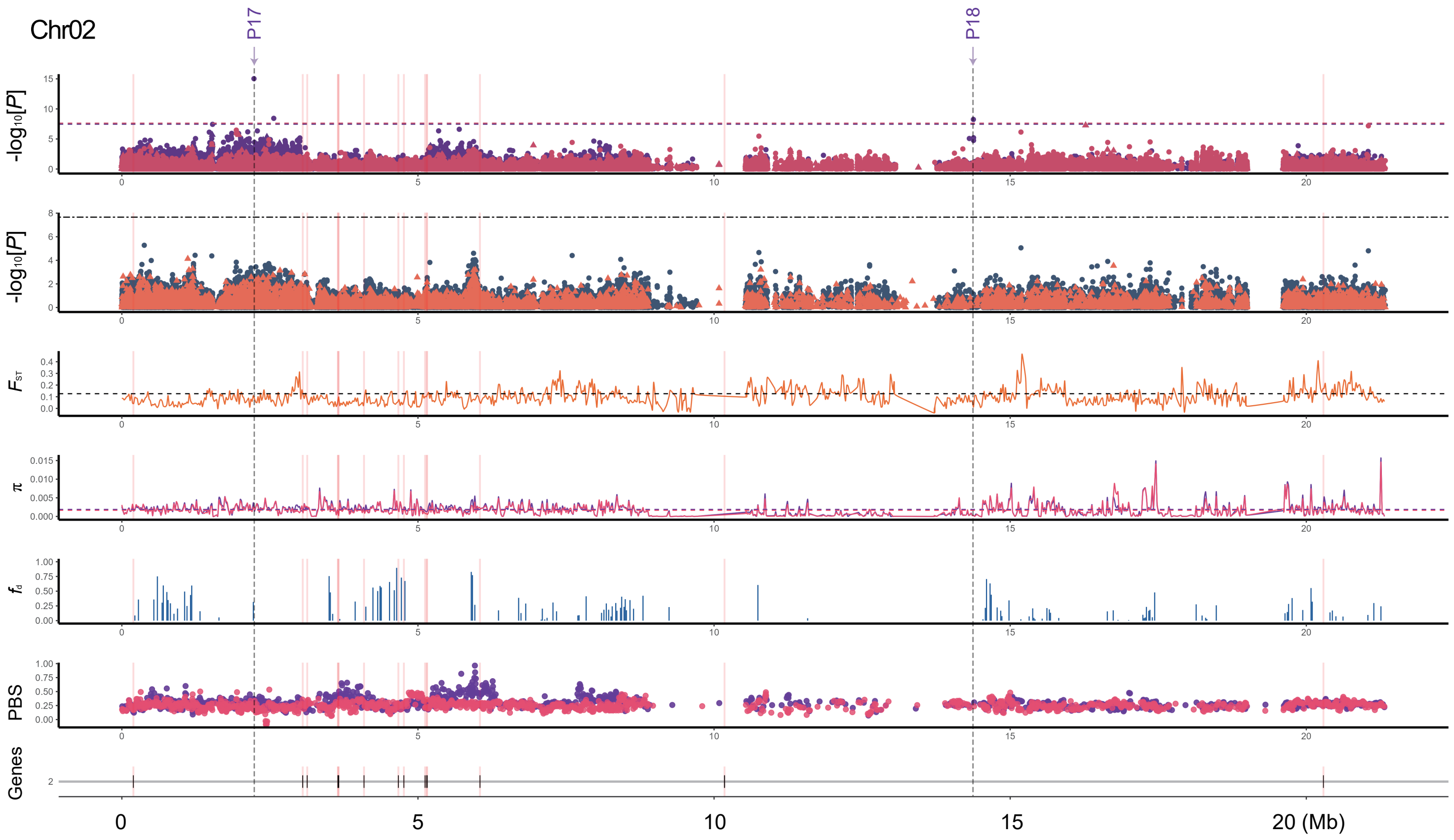
Chr02
P17
P18
-log10[P]
FST
π
fd
PBS
Genes
0
5
10
15
20 (Mb)

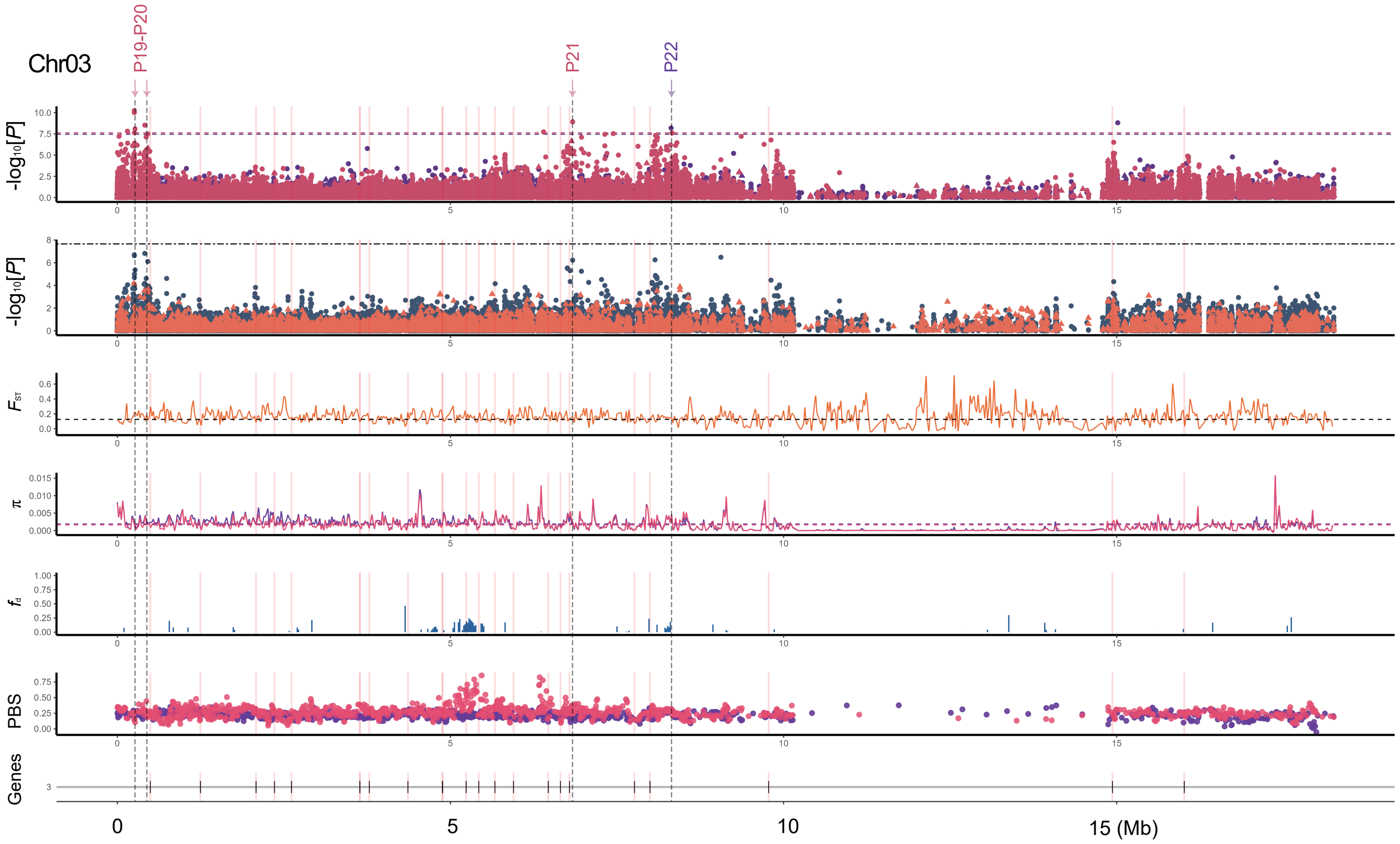
Chr03
P19-P20
P21
P22
-log10[P]
-log10[P]
FST
π
fd
PBS
Genes
0
5
10
15 (Mb)

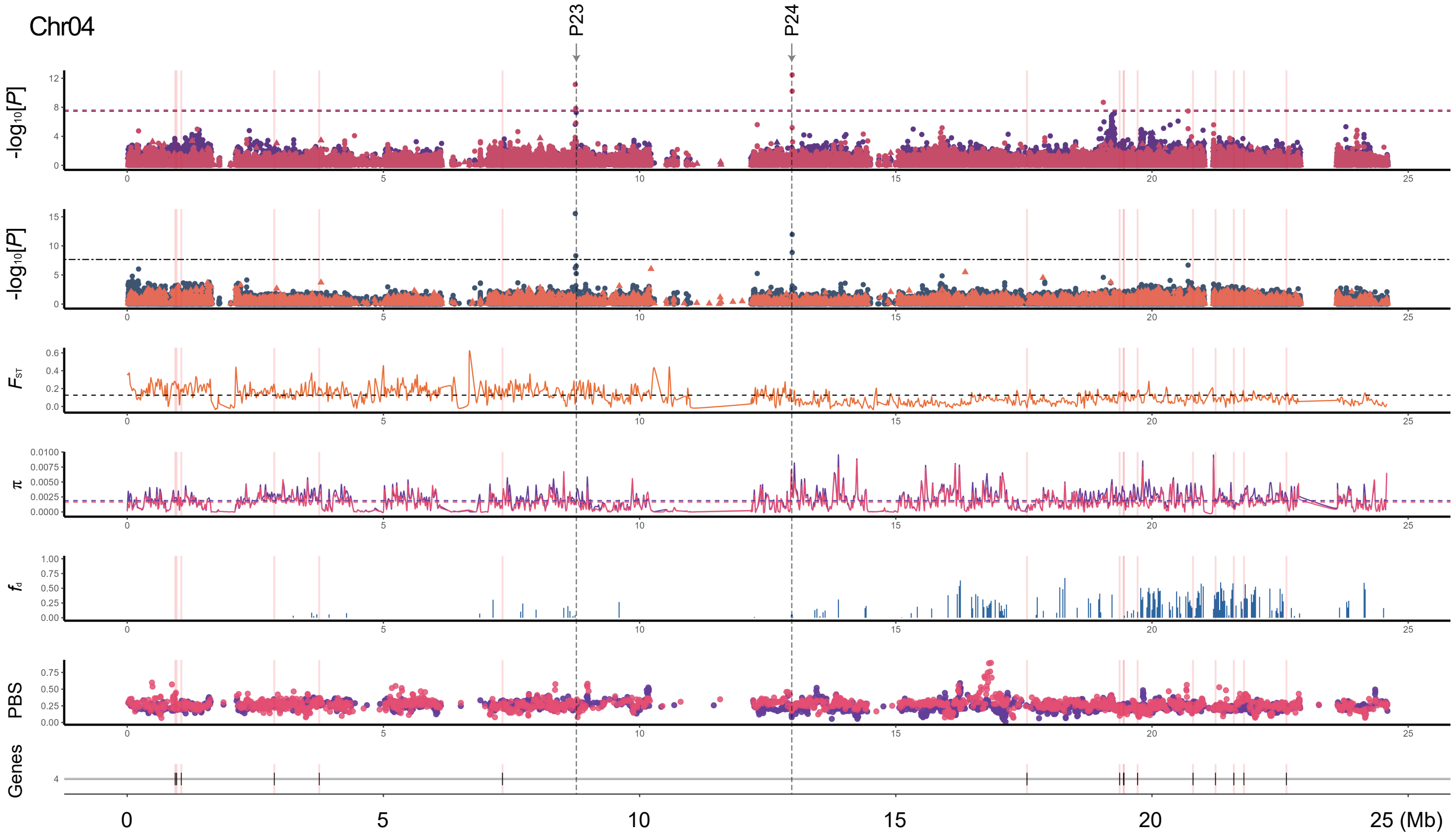
Chr04
P23
P24
-log10[P]
-log10[P]
FST
π
fd
PBS
Genes
0
5
10
15
20
25 (Mb)

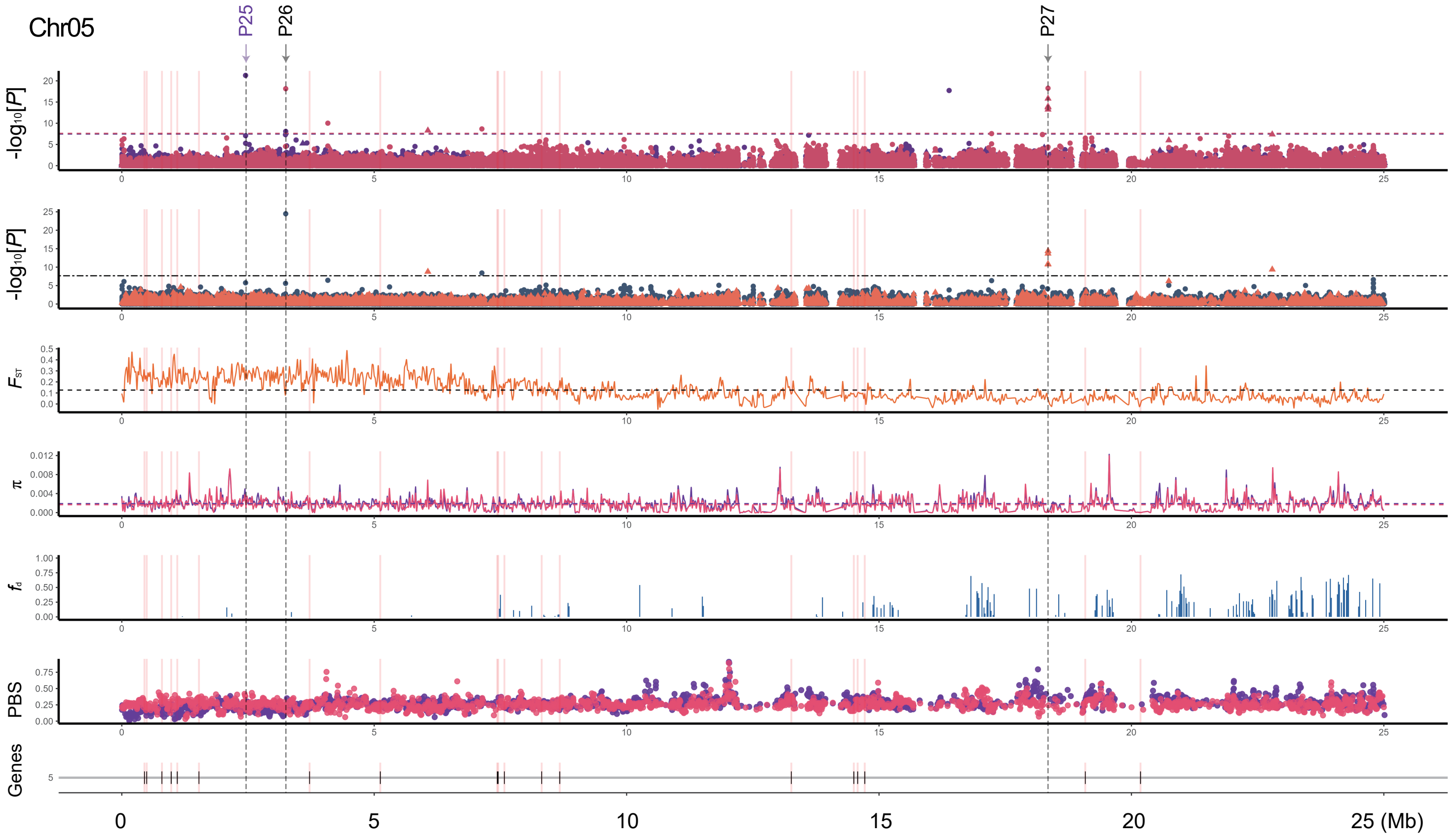
Chr05
P25
P26
P27
-log10[P]
-log10[P]
FST
π
fd
PBS
Genes
0
5
10
15
20
25 (Mb)

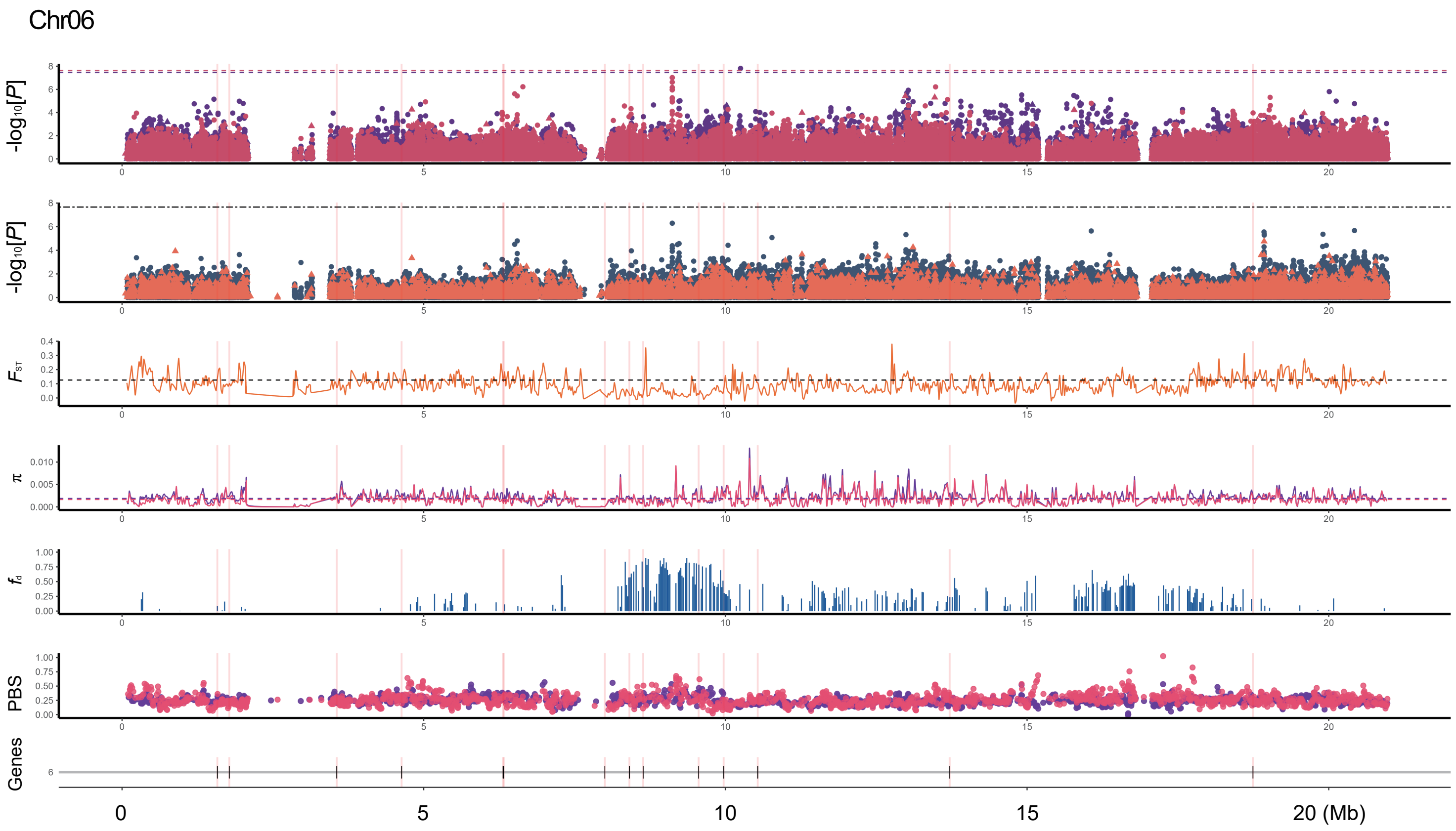
Chr06
-log10[P]
-log10[P]
FST
π
fd
PBS
Genes
0
5
10
15
20 (Mb)

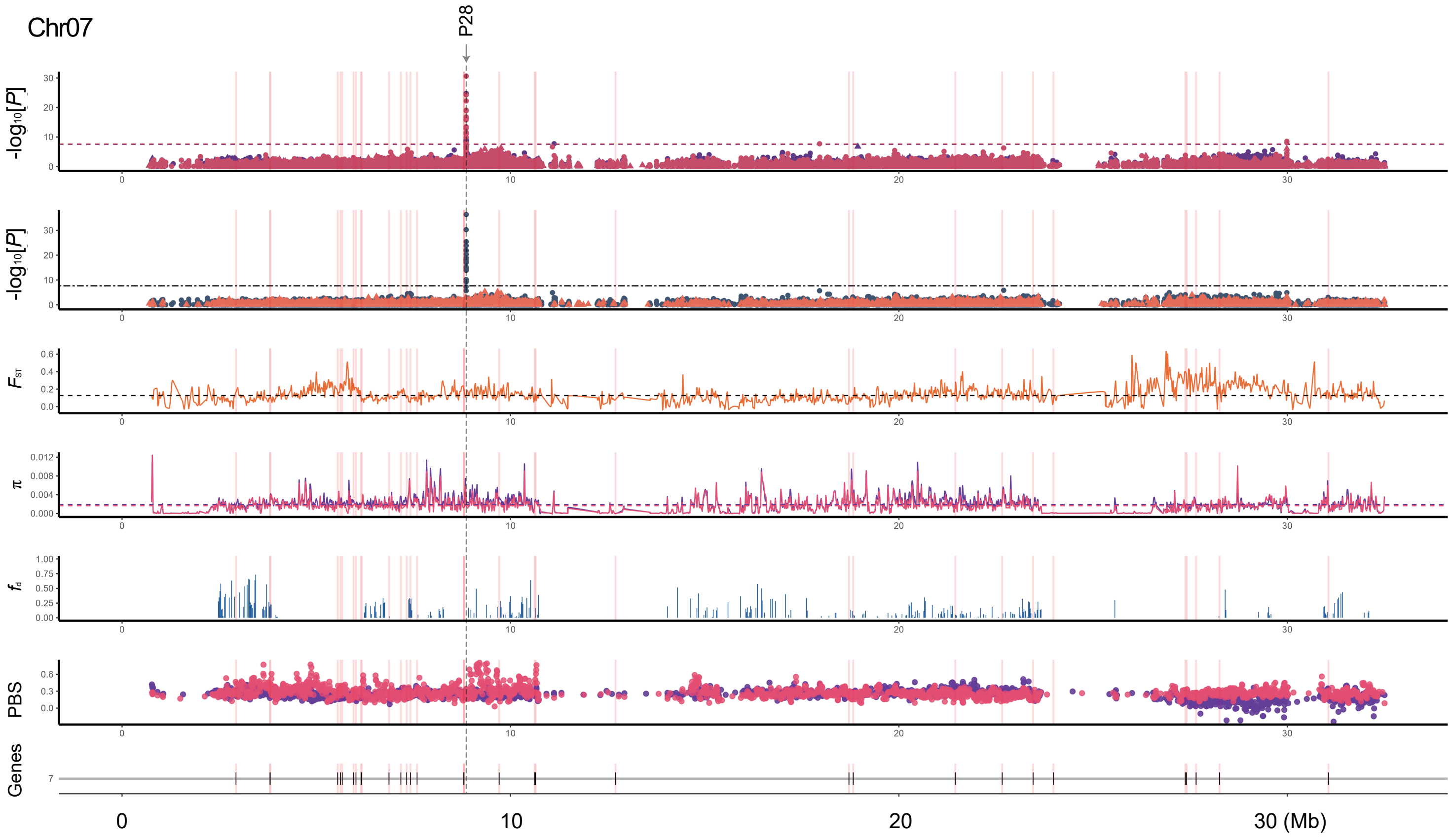
Chr07
P28
-log10[P]
-log10[P]
FST
π
fd
PBS
Genes
0
10
20
30 (Mb)

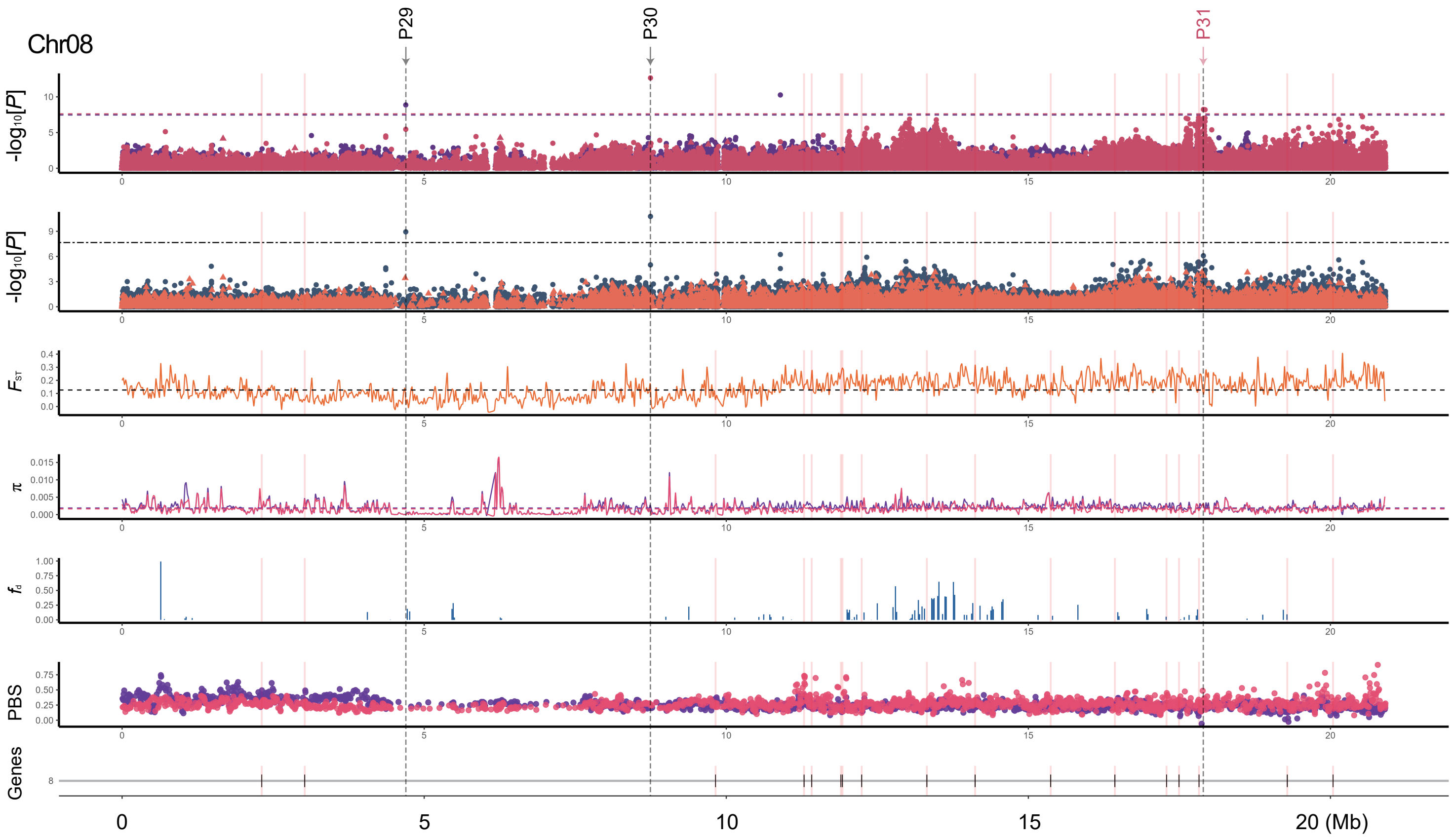
Chr08
P29
P30
P31
-log10[P]
-log10[P]
FST
π
fd
PBS
Genes
0
5
10
15
20 (Mb)

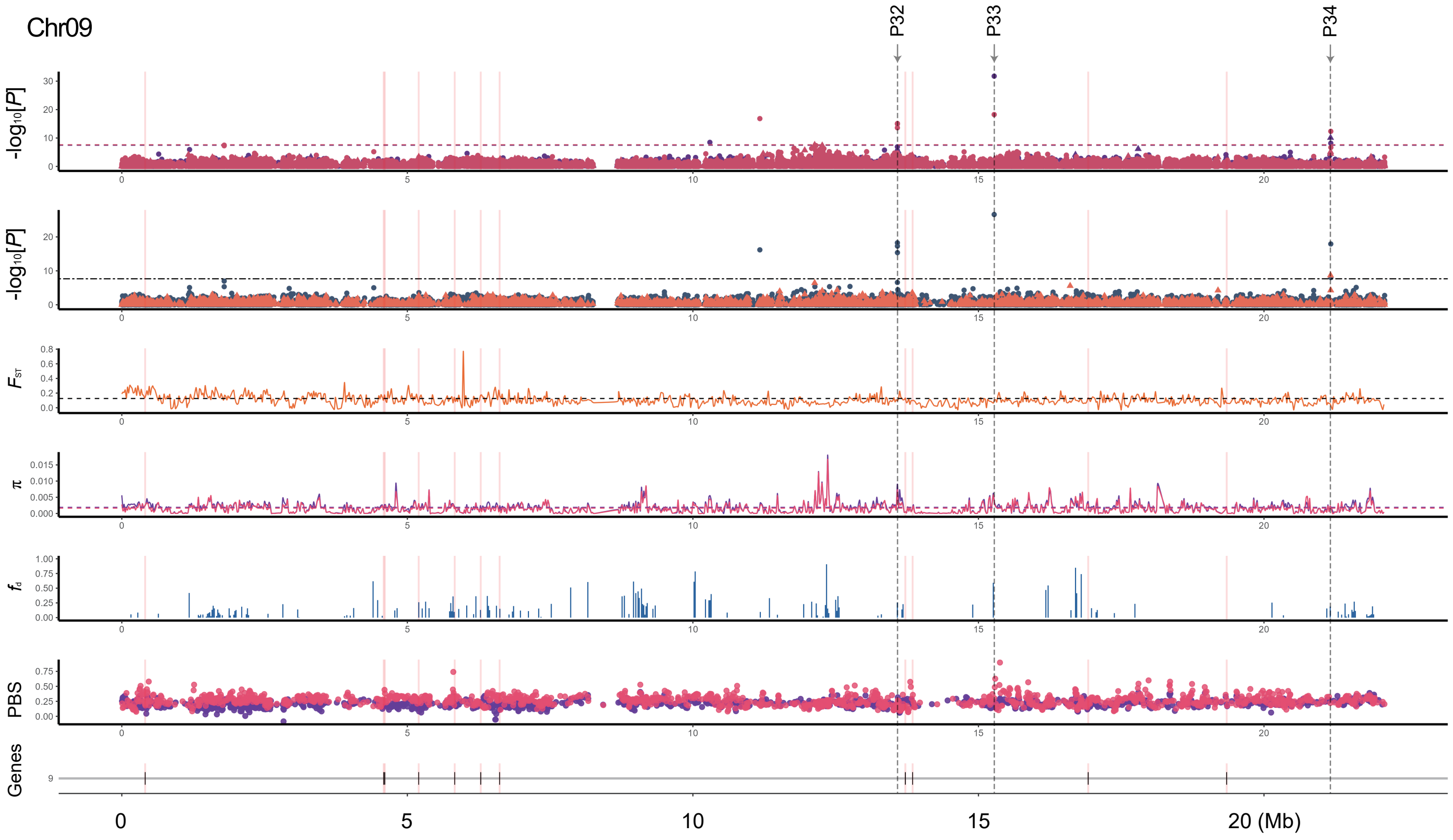
Chr09
P32
P33
P34
-log10[P]
-log10[P]
FST
π
fd
PBS
Genes
0
5
10
15
20 (Mb)

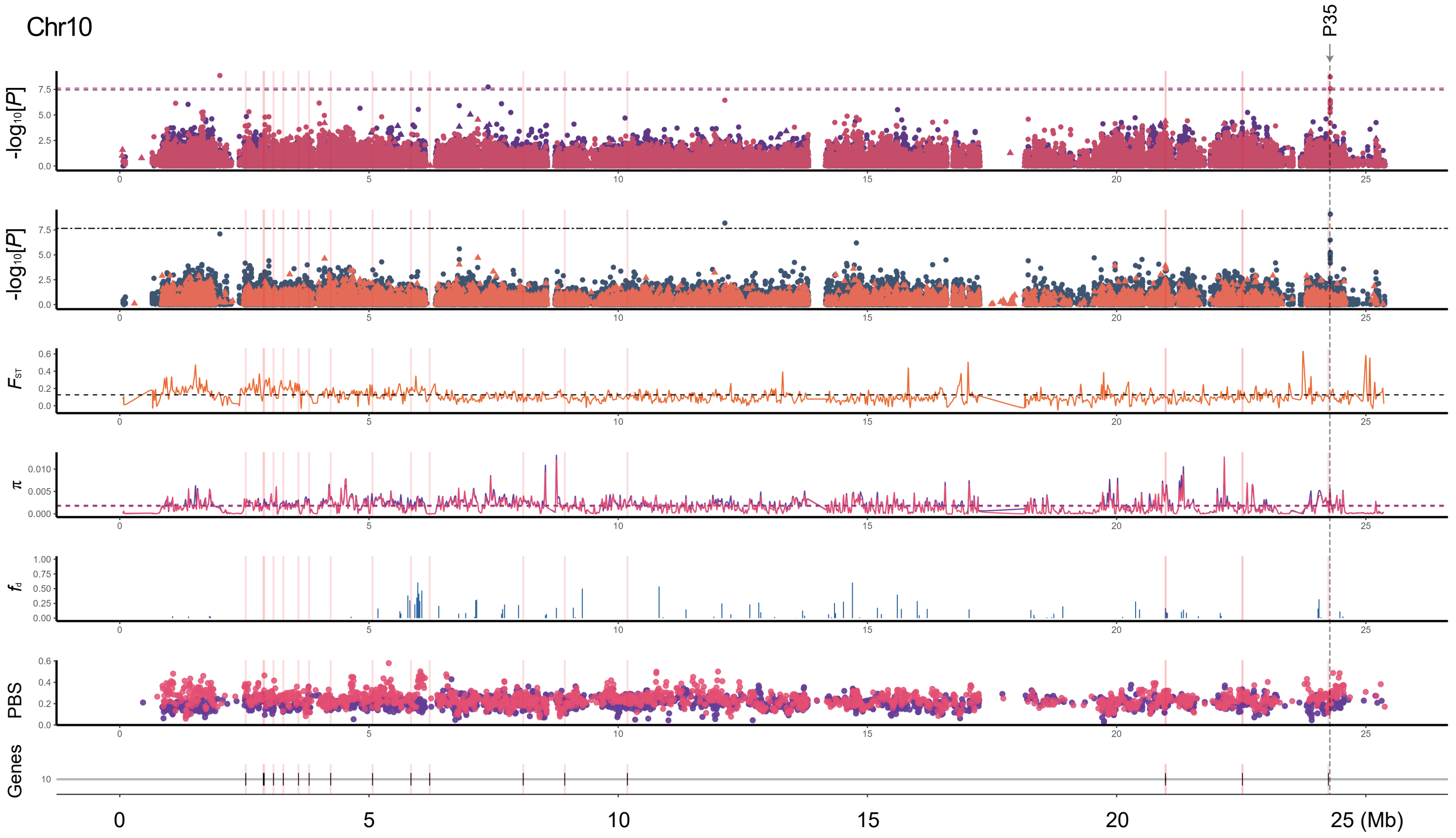
Chr10
P35
-log10[P]
-log10[P]
FST
π
fd
PBS
Genes
0
5
10
15
20
25 (Mb)

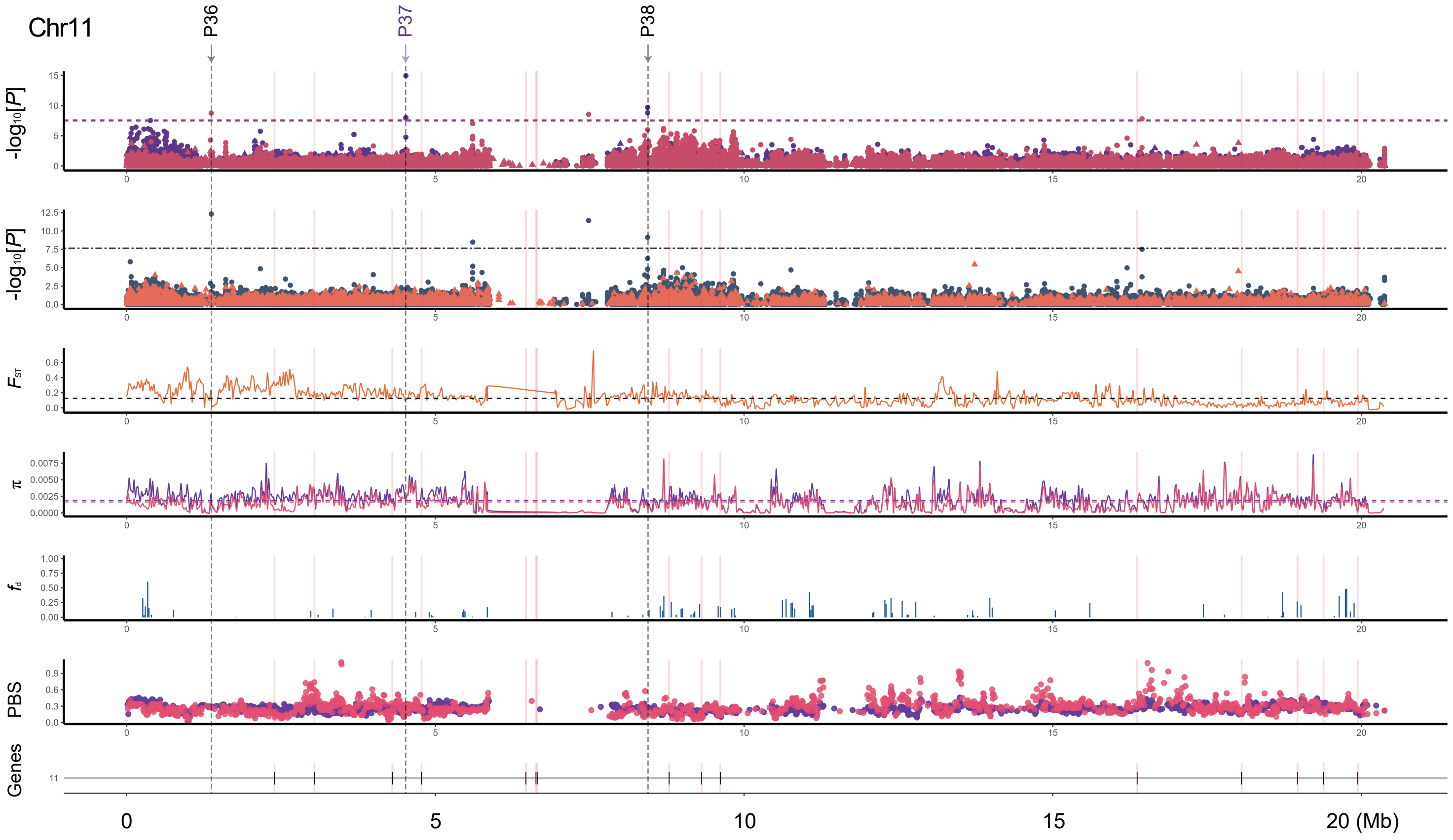
Chr11
P36
P37
P38
-log10[P]
-log10[P]
FST
π
fd
PBS
Genes
0
5
10
15
20 (Mb)

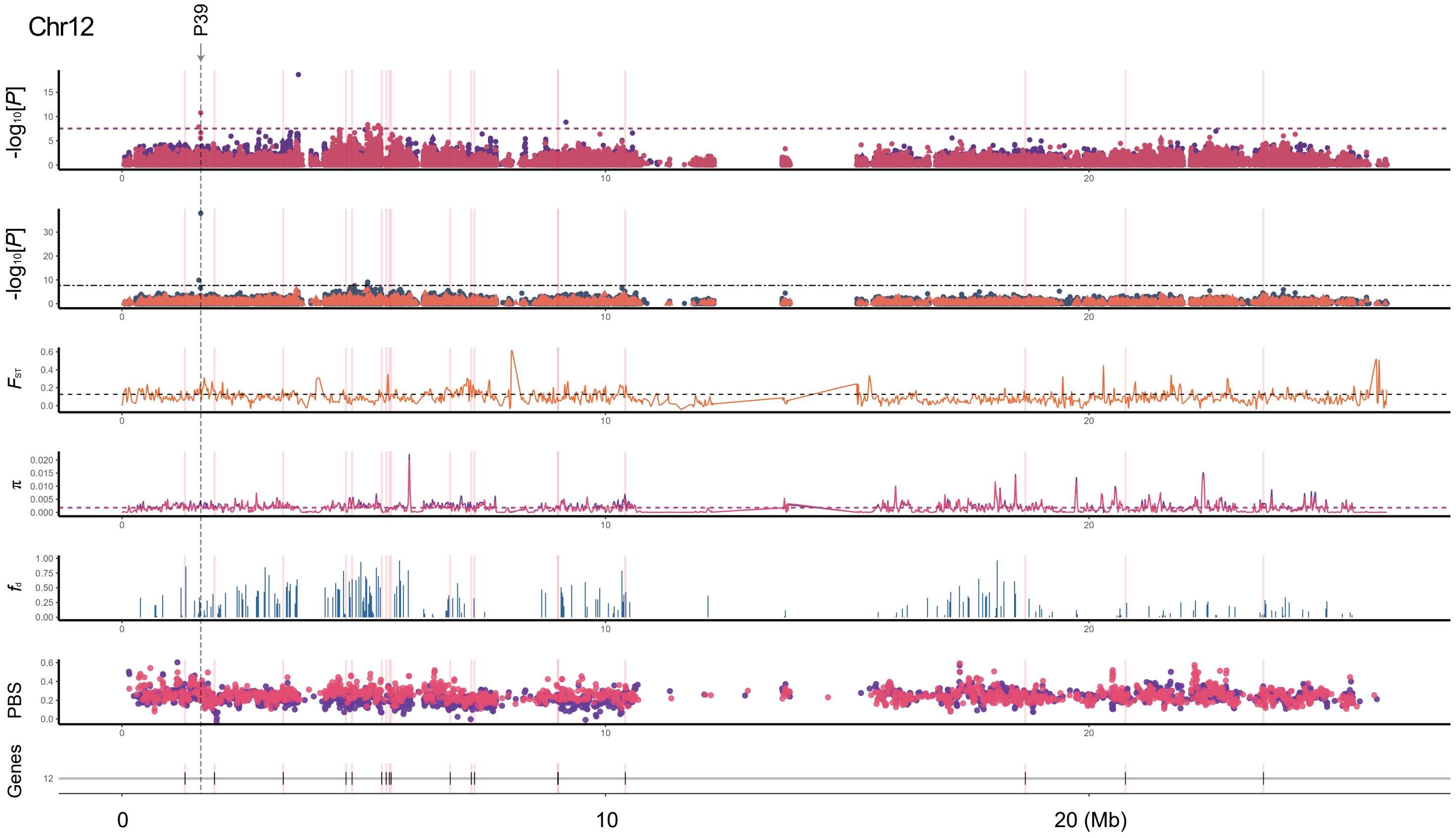
Chr12
P39
-log10[P]
-log10[P]
FST
π
fd
PBS
Genes
0
10
20 (Mb)

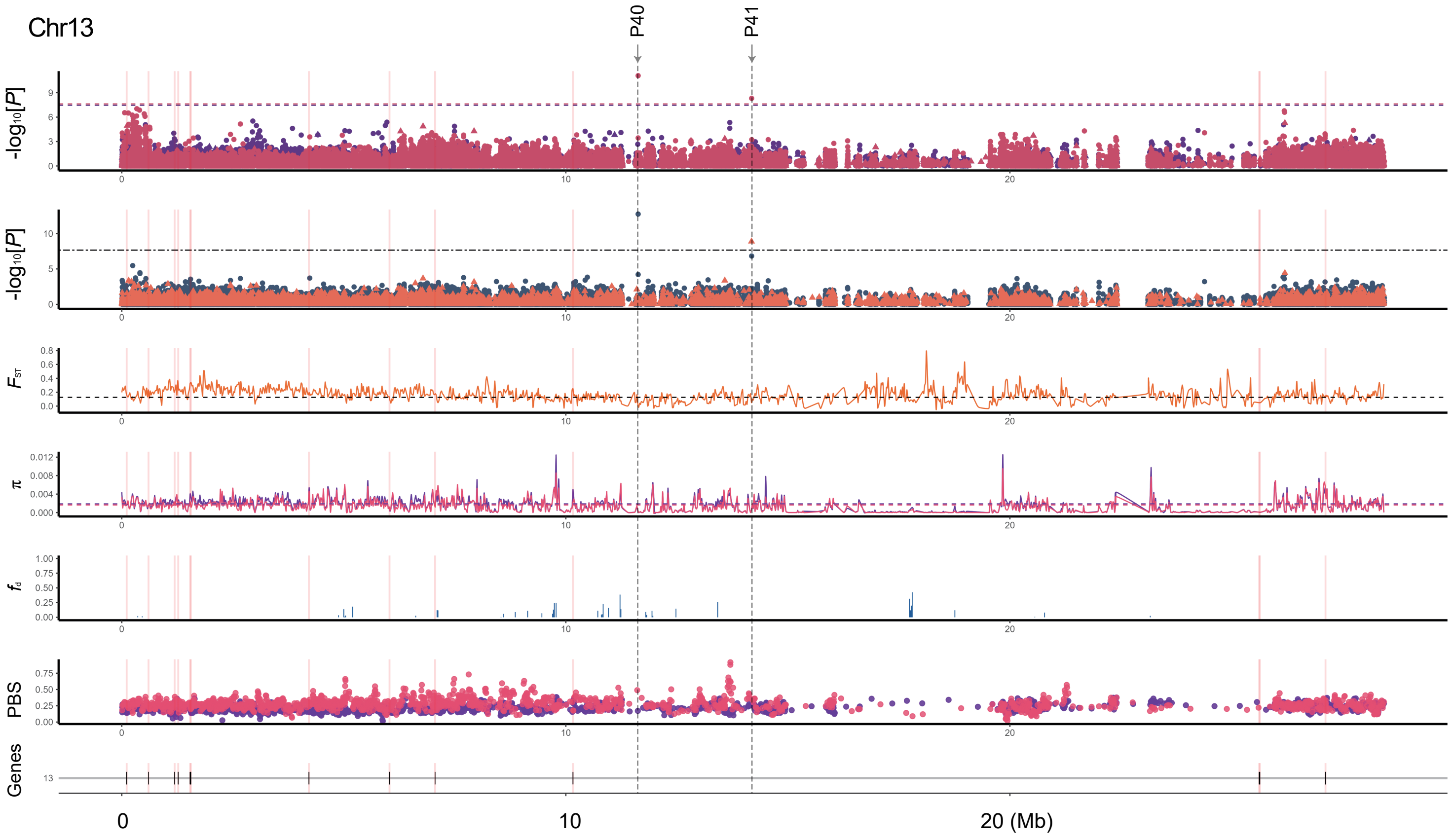
Chr13
P40
P41
-log10[P]
-log10[P]
FST
π
fd
PBS
Genes
13
0
10
20 (Mb)

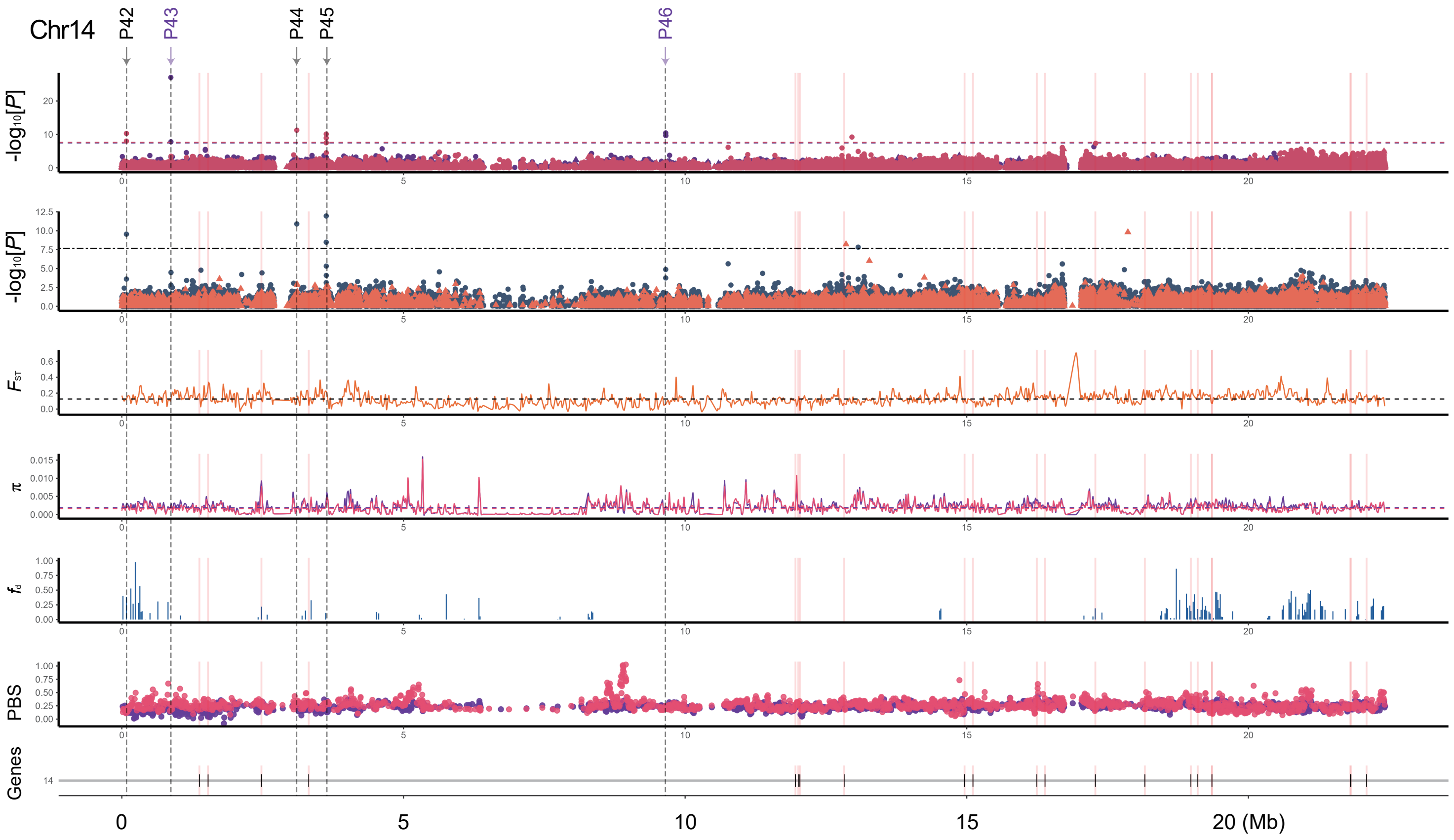
Chr14
P42
P43
P44
P45
P46
-log10[P]
-log10[P]
FST
π
fd
PBS
Genes
14
0
5
10
15
20 (Mb)

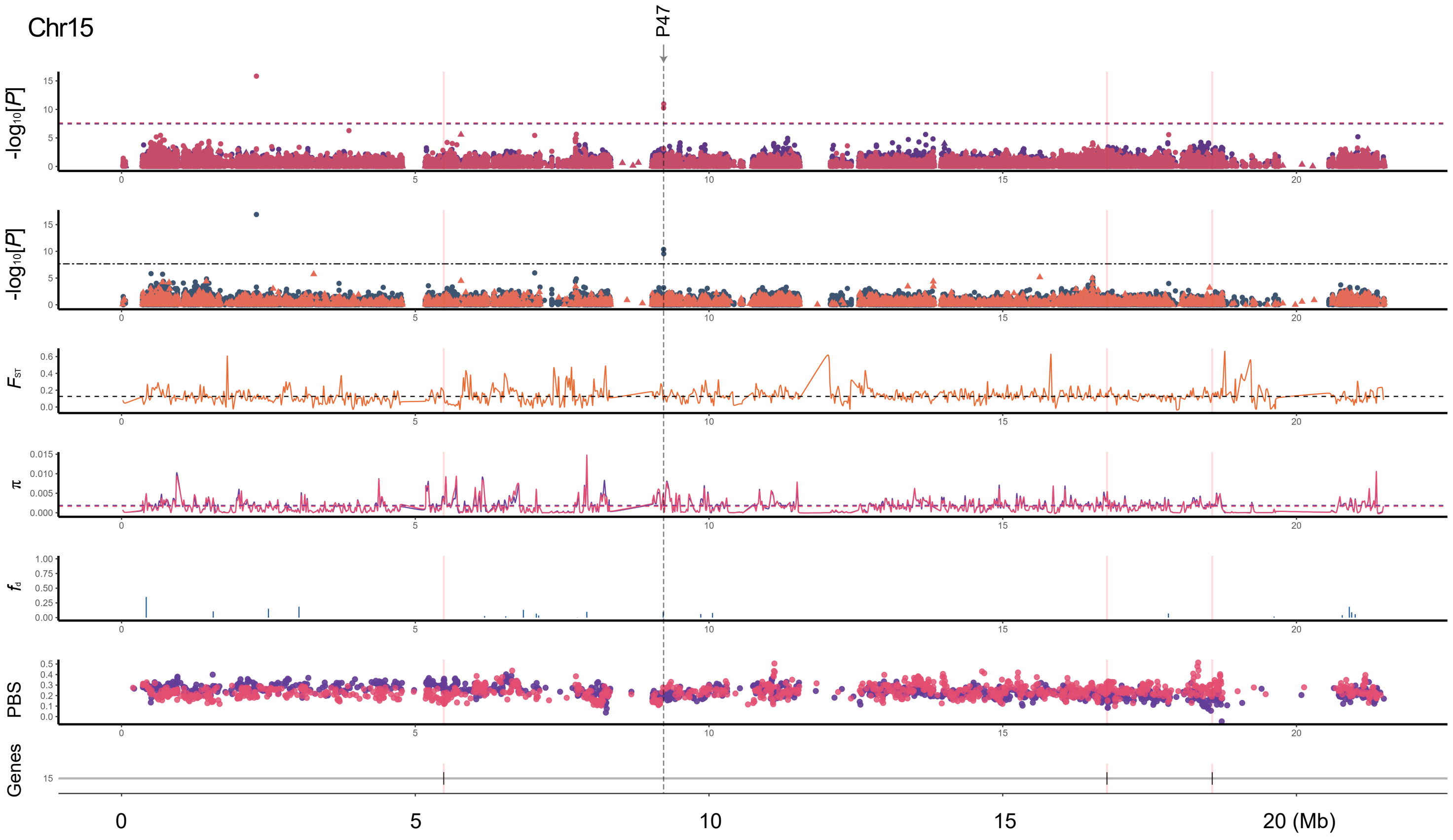
Chr15
P47
-log10[P]
FST
π
fd
PBS
Genes
0
5
10
15
20 (Mb)

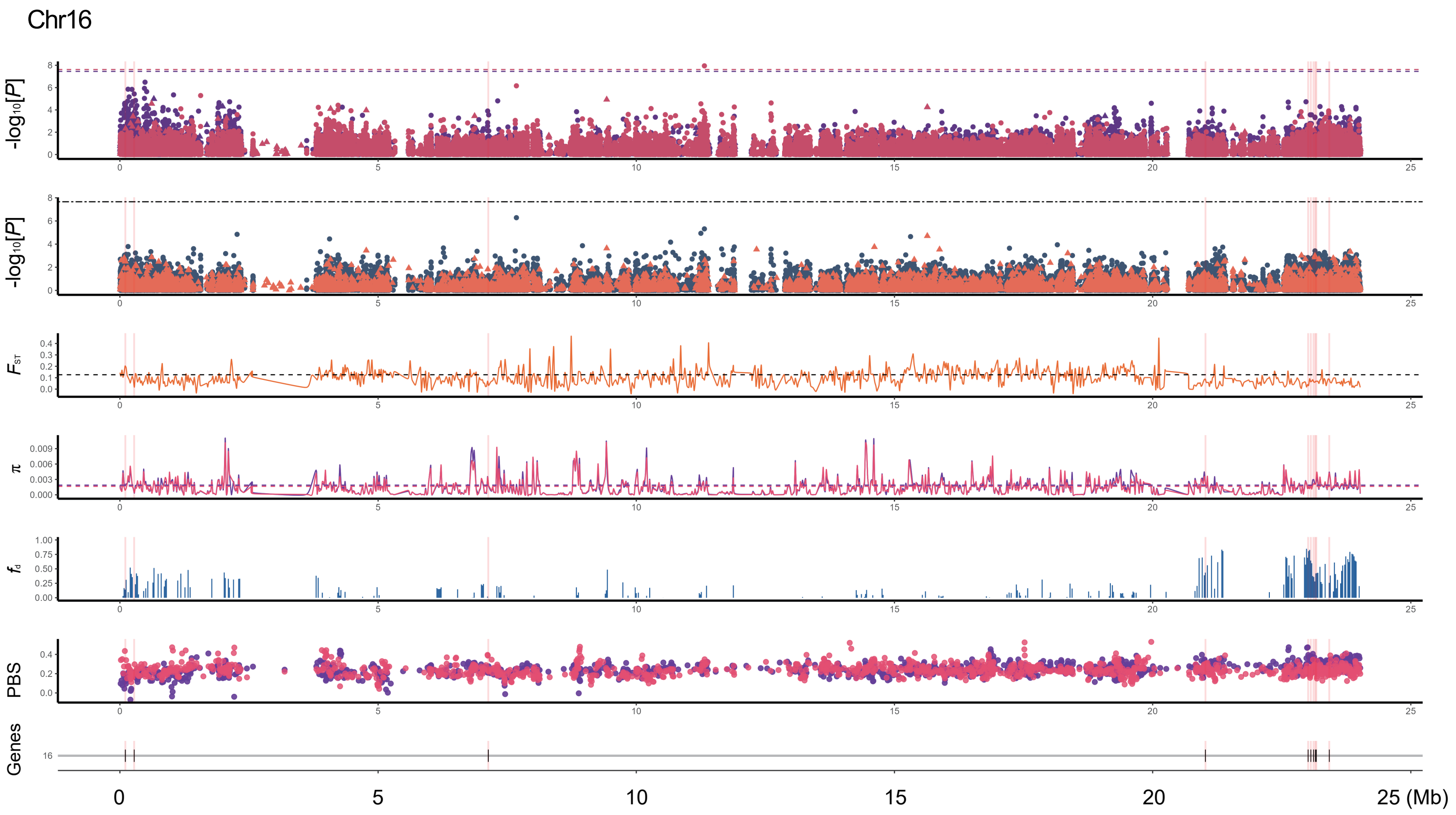
Chr16
-log10[P]
FST
π
fd
PBS
Genes
0
5
10
15
20
25 (Mb)

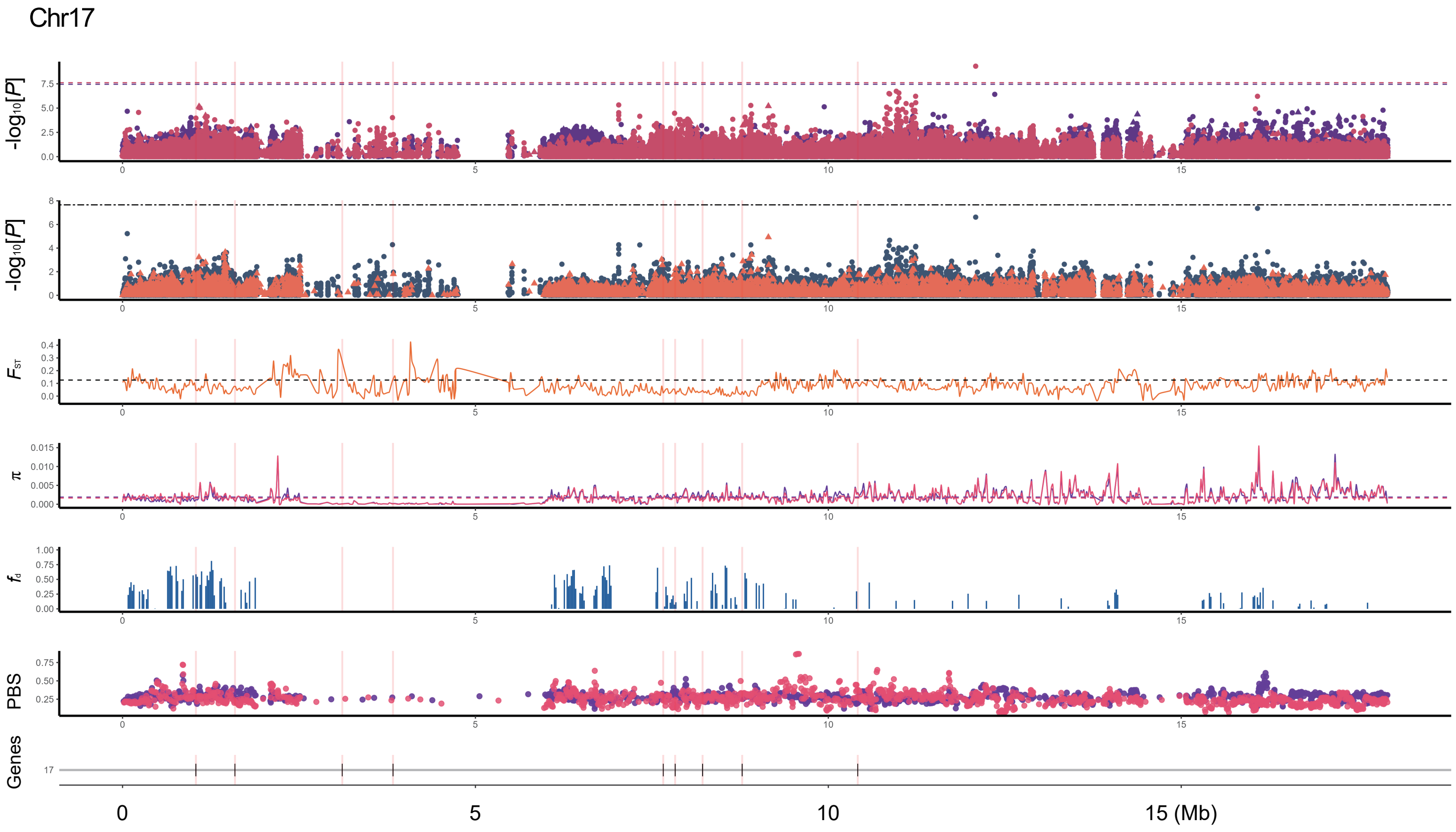
Chr17
-log10[P]
-log10[P]
FST
π
fd
PBS
Genes
17
0
5
10
15 (Mb)

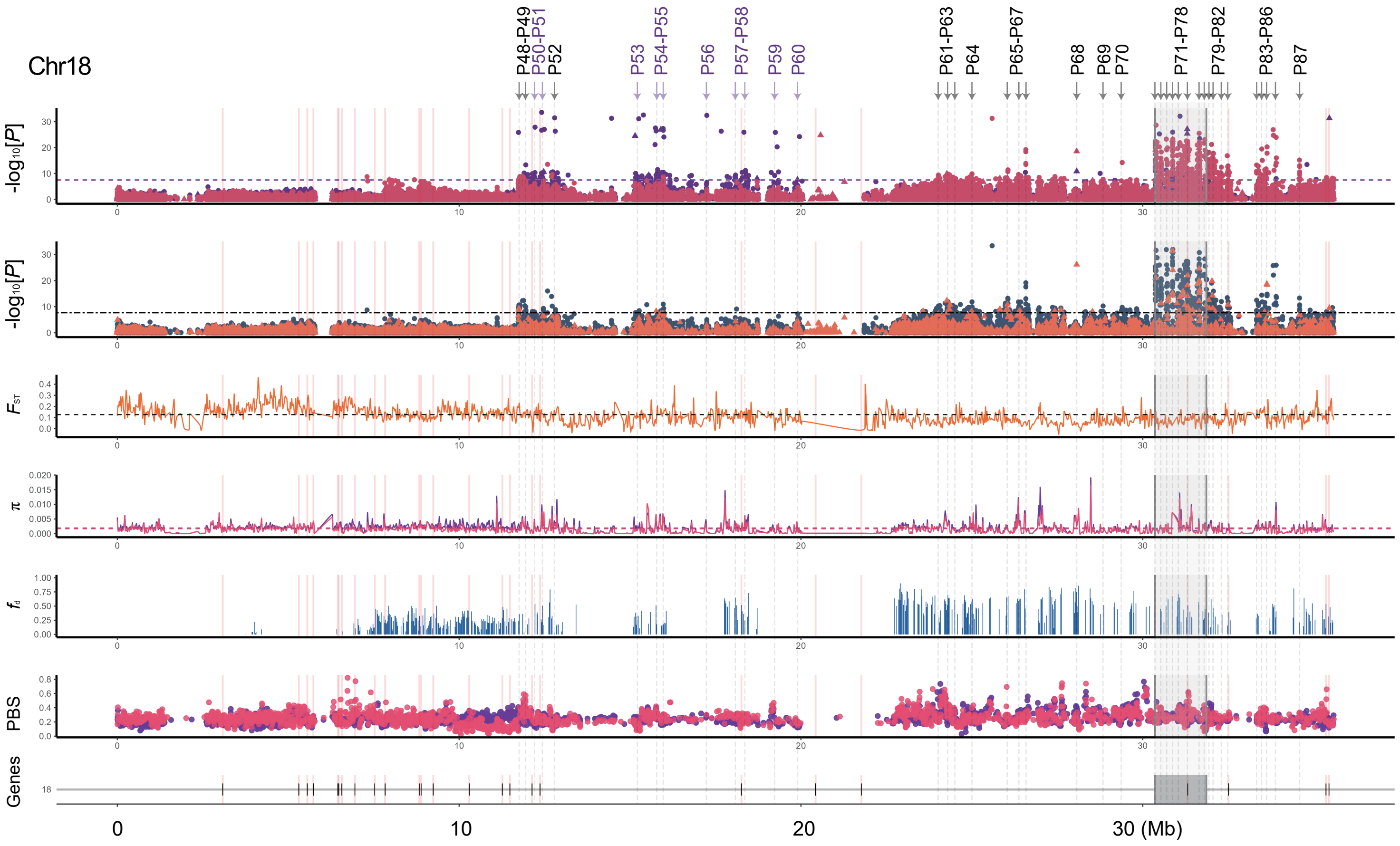
Chr18
P48-P49
P50-P51
P52
P53
P54-P55
P56
P57-P58
P59
P60
P61-P63
P64
P65-P67
P68
P69
P70
P71-P78
P79-P82
P83-P86
P87
-log10[P]
-log10[P]
FST
π
fd
PBS
Genes
18
0
10
20
30 (Mb)

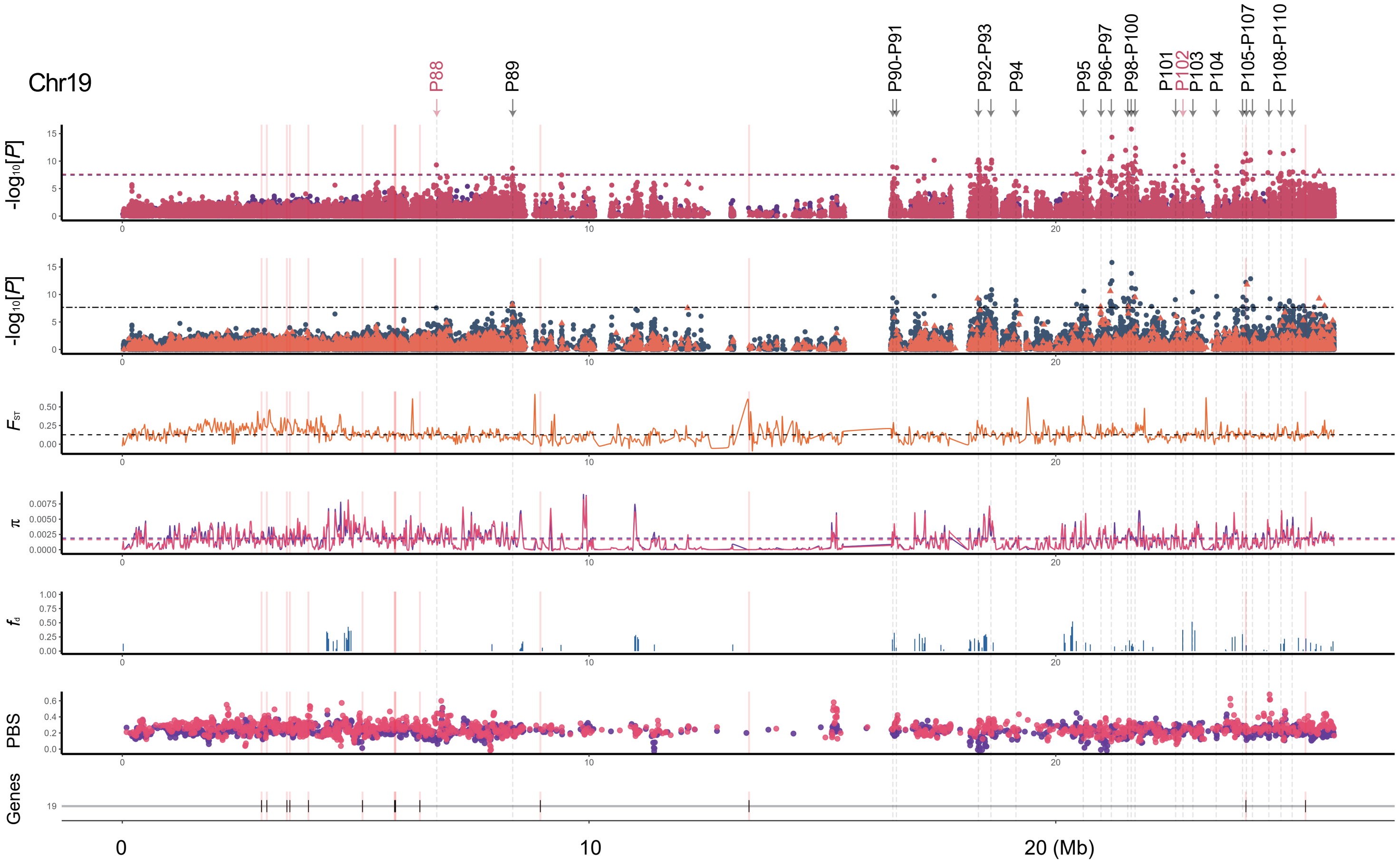


**Extended Data Fig. 7 | Visualization of whole-genome analyses.** From top to bottom, hese results included QTL peaks (red: VV, purple: VV×VL, black: consensus peaks), GWAS analyses within three populations (red: VV, purple: VV×VL, dark-blue and yellow: admixed population; points: SNPs, triangles: InDels and SVs), population analyses of fixation indices ($F_{ST}$), nucleotide diversity (π), introgression ($f_d$), and divergent selection (PBS) (refer to Supplementary Table 13), and 339 core candidate genes (refer to Supplementary Table 11).

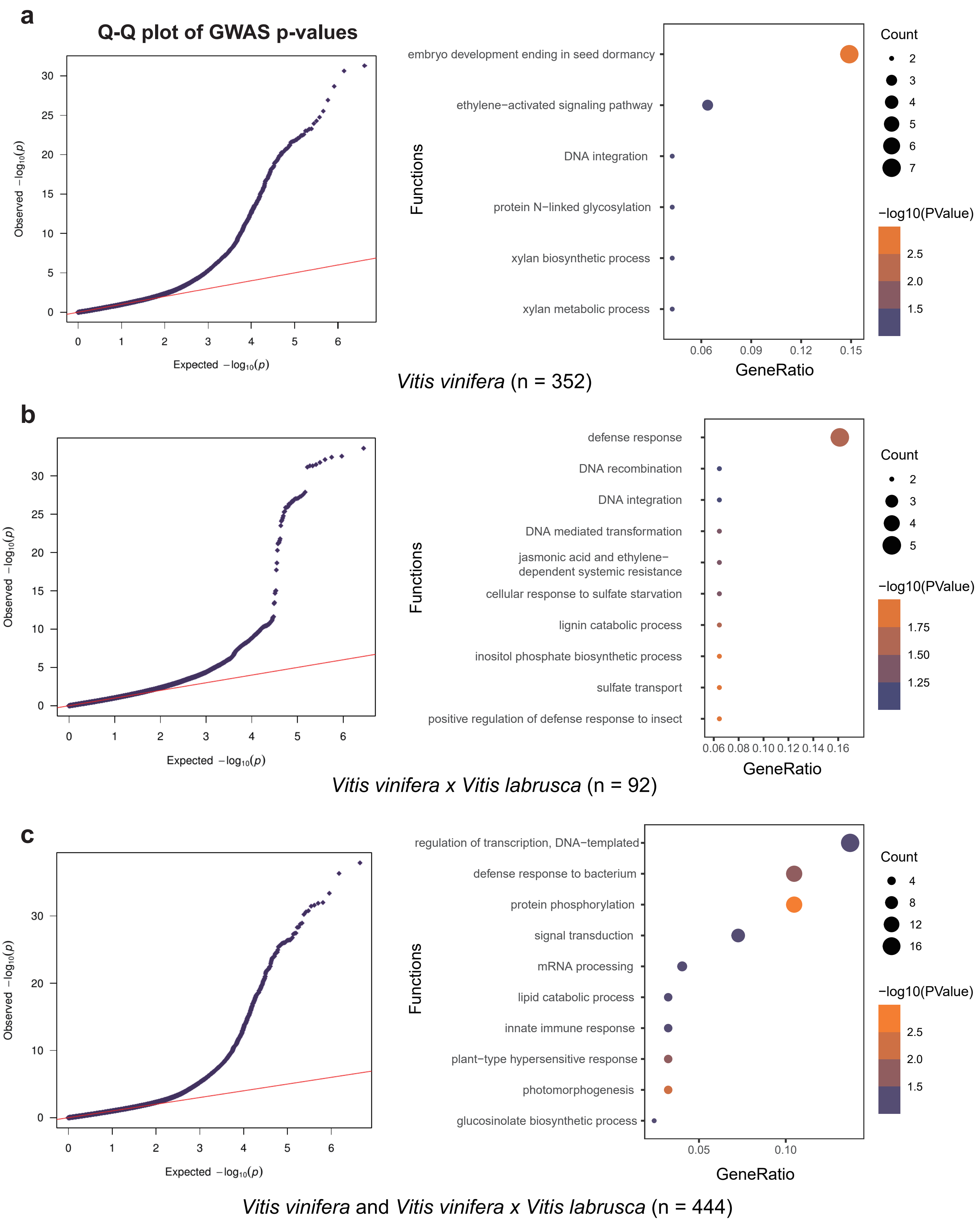


**Extended Data Fig. 8 | Quantile-Quantile (Q-Q) plot and Gene Ontology (GO) enrichment analysis.** Genome-Wide Association Study (GWAS) Q-Q plot for the three populations, along with GO enrichment analysis for biological processes in genes specific to VV×VL and VV, as well as those consensus gene for admixed populations (refer to Supplementary Table 12).

```
                    *        20         *        40         *        60         *        80         *       100
PN_T2T_G1  : ---------------------------------------------------------------------------------------------------- :    -
CShap1_G1  : ---------------------------------------------------------------------------------------------------- :    -
CShap2_G1  : ---------------------------------------------------------------------------------------------------- :    -
MHhap1_G1  : ---------------------------------------------------------------------------------------------------- :    -
MHhap2_G1  : ---------------------------------------------------------------------------------------------------- :    -
SMhap1_G1  : ---------------------------------------------------------------------------------------------------- :    -
SMhap2_G1  : ---------------------------------------------------------------------------------------------------- :    -
BCsd_G1    : ---------------------------------------------------------------------------------------------------- :    -
BCs1_G1    : ---------------------------------------------------------------------------------------------------- :    -
TShap1_G1  : ---------------------------------------------------------------------------------------------------- :    -
TShap2_G1  : ---------------------------------------------------------------------------------------------------- :    -
BMhap1_G1  : ---------------------------------------------------------------------------------------------------- :    -
BMhap2_G1  : ---------------------------------------------------------------------------------------------------- :    -
MRhap2_G1  : ---------------------------------------------------------------------------------------------------- :    -
PN_T2T_G2  : ---------------------------------------------------------------------------------------------------- :    -
CShap1_G2  : MAKRLCLLLCFLVLFQCCMGRQQTSRQQKECRISRLNAQRPSNRIQSEAGVTEVFDHNNEQFQCAGVAVVRYTIEPRGLLLPSYVNAPQLMYFVQGRGLQ :  100
CShap2_G2  : MAKRLCLLLCFLVLFQCCMGRQQTSRQQKECRISRLNAQRPSNRIQSEAGVTEVFDHNNEQFQCAGVAVVRYTIEPRGLLLPSYVNAPQLMYFVQGRGLQ :  100
MHhap1_G2  : ---------------------------------------------------------------------------------------------------- :    -
MHhap2_G2  : ---------------------------------------------------------------------------------------------------- :    -
SMhap1_G2  : ---------------------------------------------------------------------------------------------------- :    -
SMhap2_G2  : ---------------------------------------------------------------------------------------------------- :    -
BCsd_G2    : ---------------------------------------------------------------------------------------------------- :    -
BCs1_G2    : ---------------------------------------------------------------------------------------------------- :    -
TShap1_G2  : ---------------------------------------------------------------------------------------------------- :    -
TShap2_G2  : ---------------------------------------------------------------------------------------------------- :    -
BMhap1_G2  : ---------------------------------------------------------------------------------------------------- :    -
BMhap2_G2  : ---------------------------------------------------------------------------------------------------- :    -
MRhap2_G2  : ---------------------------------------------------------------------------------------------------- :    -
PN_T2T_G3  : ---------------------------------------------------------------------------------------------------- :    -
MHhap1_G3  : ---------------------------------------------------------------------------------------------------- :    -
MHhap2_G3  : ---------------------------------------------------------------------------------------------------- :    -
SMhap1_G3  : ---------------------------------------------------------------------------------------------------- :    -
SMhap2_G3  : ---------------------------------------------------------------------------------------------------- :    -
BCsd_G3    : ---------------------------------------------------------------------------------------------------- :    -
BCs1_G3    : ---------------------------------------------------------------------------------------------------- :    -
TShap1_G3  : ---------------------------------------------------------------------------------------------------- :    -
TShap2_G3  : ---------------------------------------------------------------------------------------------------- :    -
BMhap1_G3  : ---------------------------------------------------------------------------------------------------- :    -
BMhap2_G3  : ---------------------------------------------------------------------------------------------------- :    -
MRhap2_G3  : ---------------------------------------------------------------------------------------------------- :    -
             MAKRLCLLLCFLVLFQCCMGRQQTSRQQKECRISRLNAQRPSNRIQSEAGVTEVFDHNNEQFQCAGVAVVRYTIEPRGLLLPSYVNAPQLMYFVQGRGLQ

                    *       120         *       140         *       160         *       180         *       200
PN_T2T_G1  : ---------------------------------------------------------------------------------------------------- :    -
CShap1_G1  : ---------------------------------------------------------------------------------------------------- :    -
CShap2_G1  : ---------------------------------------------------------------------------------------------------- :    -
MHhap1_G1  : ---------------------------------------------------------------------------------------------------- :    -
MHhap2_G1  : ---------------------------------------------------------------------------------------------------- :    -
SMhap1_G1  : ---------------------------------------------------------------------------------------------------- :    -
SMhap2_G1  : ---------------------------------------------------------------------------------------------------- :    -
BCsd_G1    : ---------------------------------------------------------------------------------------------------- :    -
BCs1_G1    : ---------------------------------------------------------------------------------------------------- :    -
TShap1_G1  : ---------------------------------------------------------------------------------------------------- :    -
TShap2_G1  : ---------------------------------------------------------------------------------------------------- :    -
BMhap1_G1  : ---------------------------------------------------------------------------------------------------- :    -
BMhap2_G1  : ---------------------------------------------------------------------------------------------------- :    -
MRhap2_G1  : ---------------------------------------------------------------------------------------------------- :    -
PN_T2T_G2  : ---------------------------------------------------------------------------------------------------- :    -
CShap1_G2  : GIMITGCPETFQSFQESQQGQEQQEQGQQGEQGQQGQQGEQGKQGQQGQQGH---QGQQGQQFRGDQHQKIREVEEGDVFAVPVGTGHFIYNNGDRQLIV :  197
CShap2_G2  : GIMITGCPETFQSFQESQQGQEQQEQGQQGQQGEQGQQGQQGQQGQQGQQGQQGQQGQQFRGDQHQKIREVEEGDVFAVPVGTGHFIYNNGDRQLIV :  200
MHhap1_G2  : ---------------------------------------------------------------------------------------------------- :    -
MHhap2_G2  : ---------------------------------------------------------------------------------------------------- :    -
SMhap1_G2  : ---------------------------------------------------------------------------------------------------- :    -
SMhap2_G2  : ---------------------------------------------------------------------------------------------------- :    -
BCsd_G2    : ---------------------------------------------------------------------------------------------------- :    -
BCs1_G2    : ---------------------------------------------------------------------------------------------------- :    -
TShap1_G2  : ---------------------------------------------------------------------------------------------------- :    -
TShap2_G2  : ---------------------------------------------------------------------------------------------------- :    -
BMhap1_G2  : ---------------------------------------------------------------------------------------------------- :    -
BMhap2_G2  : ---------------------------------------------------------------------------------------------------- :    -
MRhap2_G2  : ---------------------------------------------------------------------------------------------------- :    -
PN_T2T_G3  : ---------------------------------------------------------------------------------------------------- :    -
MHhap1_G3  : ---------------------------------------------------------------------------------------------------- :    -
MHhap2_G3  : ---------------------------------------------------------------------------------------------------- :    -
SMhap1_G3  : ---------------------------------------------------------------------------------------------------- :    -
SMhap2_G3  : ---------------------------------------------------------------------------------------------------- :    -
BCsd_G3    : ---------------------------------------------------------------------------------------------------- :    -
BCs1_G3    : ---------------------------------------------------------------------------------------------------- :    -
TShap1_G3  : ---------------------------------------------------------------------------------------------------- :    -
TShap2_G3  : ---------------------------------------------------------------------------------------------------- :    -
BMhap1_G3  : ---------------------------------------------------------------------------------------------------- :    -
BMhap2_G3  : ---------------------------------------------------------------------------------------------------- :    -
MRhap2_G3  : ----------------MAECSQISDETNENENTQKHQEIHLEDAESRNPDDESNSHTEMKKPISPEQEYKASIQMKKSKKKALIELRCRVEDAILGNYML :   84
             GIMITGCPETFQSFQESQQGQEQQEQGQQGEQGQQGQQGQQGQQGQQGQQGQQGQQGQQFRGDQHQKIREVEEGDVFAVPVGTGHFIYNNGDRQLIV

                    *       220         *       240         *       260         *       280         *       300
PN_T2T_G1  : ---------------------------------------------------------------------------------------------------- :    -
CShap1_G1  : ---------------------------------------------------------------------------------------------------- :    -
CShap2_G1  : ---------------------------------------------------------------------------------------------------- :    -
MHhap1_G1  : ---------------------------------------------------------------------------------------------------- :    -
```

```
MHhap2_G1 : ---------------------------------------------------------------------------------------------------- :    -
SMhap1_G1 : ---------------------------------------------------------------------------------------------------- :    -
SMhap2_G1 : ---------------------------------------------------------------------------------------------------- :    -
BCsd_G1   : ---------------------------------------------------------------------------------------------------- :    -
BCsl_G1   : ---------------------------------------------------------------------------------------------------- :    -
TShap1_G1 : ---------------------------------------------------------------------------------------------------- :    -
TShap2_G1 : ---------------------------------------------------------------------------------------------------- :    -
BMhap1_G1 : ---------------------------------------------------------------------------------------------------- :    -
BMhap2_G1 : ---------------------------------------------------------------------------------------------------- :    -
MRhap2_G1 : ---------------------------------------------------------------------------------------------------- :    -
PN_T2T_G2 : ---------------------------------------------------------------------------------------------------- :    -
CShap1_G2 : VSVLDTSNDANQLDFQPRRFYLAGNPQNEFQQQQQQQQGSEGQQQQQEGGGSEGRGQESSGNNIFSGFDAQQLAEAFNVDVQIIRKLQGQNDRRGNIVRV :  297
CShap2_G2 : VSVLDTSNDANQLDFQPRRFYLAGNPQNEFQQQQQQQQGSEGQQQQQEGGGSEGRGQESSGDNIFSGFDAQQLAEAFNVDVQLIRKLQGQNDRRGNIVRV :  300
MHhap1_G2 : ---------------------------------------------------------------------------------------------------- :    -
MHhap2_G2 : ---------------------------------------------------------------------------------------------------- :    -
SMhap1_G2 : ---------------------------------------------------------------------------------------------------- :    -
SMhap2_G2 : ---------------------------------------------------------------------------------------------------- :    -
BCsd_G2   : ---------------------------------------------------------------------------------------------------- :    -
BCsl_G2   : ---------------------------------------------------------------------------------------------------- :    -
TShap1_G2 : ---------------------------------------------------------------------------------------------------- :    -
TShap2_G2 : ---------------------------------------------------------------------------------------------------- :    -
BMhap1_G2 : ---------------------------------------------------------------------------------------------------- :    -
BMhap2_G2 : ---------------------------------------------------------------------------------------------------- :    -
MRhap2_G2 : ---------------------------------------------------------------------------------------------------- :    -
PN_T2T_G3 : ---------------------------------------------------------------------------------------------------- :    -
MHhap1_G3 : ---------------------------------------------------------------------------------------------------- :    -
MHhap2_G3 : ---------------------------------------------------------------------------------------------------- :    -
SMhap1_G3 : ---------------------------------------------------------------------------------------------------- :    -
SMhap2_G3 : ---------------------------------------------------------------------------------------------------- :    -
BCsd_G3   : ---------------------------------------------------------------------------------------------------- :    -
BCsl_G3   : ---------------------------------------------------------------------------------------------------- :    -
TShap1_G3 : ---------------------------------------------------------------------------------------------------- :    -
TShap2_G3 : ---------------------------------------------------------------------------------------------------- :    -
BMhap1_G3 : ---------------------------------------------------------------------------------------------------- :    -
BMhap2_G3 : ---------------------------------------------------------------------------------------------------- :    -
MRhap2_G3 : GKPSGKVPESNKTQENLHDISLWGVPLLPSKGHEGTDIILLKFLKARDFRVSEAFNMLRRTLIWRREFKTEGILEEN------------FGPELENVVYI :  172
            VSVLDTSNDANQLDFQPRRFYLAGNPQNEFQQQQQQQQGSEGQQQQQEGGGSEGRGQESSGDNIFSGFDAQQLAEAFNVDVQIIRKLQGQNDRRGNIVRV

                    *        320         *        340         *        360         *        380         *        400
PN_T2T_G1 : ---------------------------------------------------------------------------------------------------- :    -
CShap1_G1 : ---------------------------------------------------------------------------------------------------- :    -
CShap2_G1 : ---------------------------------------------------------------------------------------------------- :    -
MHhap1_G1 : ---------------------------------------------------------------------------------------------------- :    -
MHhap2_G1 : ---------------------------------------------------------------------------------------------------- :    -
SMhap1_G1 : ---------------------------------------------------------------------------------------------------- :    -
SMhap2_G1 : ---------------------------------------------------------------------------------------------------- :    -
BCsd_G1   : ---------------------------------------------------------------------------------------------------- :    -
BCsl_G1   : ---------------------------------------------------------------------------------------------------- :    -
TShap1_G1 : ---------------------------------------------------------------------------------------------------- :    -
TShap2_G1 : ---------------------------------------------------------------------------------------------------- :    -
BMhap1_G1 : ---------------------------------------------------------------------------------------------------- :    -
BMhap2_G1 : ---------------------------------------------------------------------------------------------------- :    -
MRhap2_G1 : ---------------------------------------------------------------------------------------------------- :    -
PN_T2T_G2 : ---------------------------------------------------------------------------------------------------- :    -
CShap1_G2 : EGGLQAVLPPRGQQERGEQQQDHFHARGNGYEETICSLRLKQNIGDPWRADVYTPRGGHRSSVTGYDLPILRKVVRLSAHQGRLHQGAMVLPYYNVNAHS :  397
CShap2_G2 : EGGLQAVLPPRGQQERGEQQQDHFHARGNGYEETICSLRLKQNIGDPWRADVYTPRGGHRSSVTGYDLPILRKVVRLSAHQGRLHQGAMVLPYYNVNAHS :  400
MHhap1_G2 : ---------------------------------------------------------------------------------------------------- :    -
MHhap2_G2 : ---------------------------------------------------------------------------------------------------- :    -
SMhap1_G2 : ---------------------------------------------------------------------------------------------------- :    -
SMhap2_G2 : ---------------------------------------------------------------------------------------------------- :    -
BCsd_G2   : ---------------------------------------------------------------------------------------------------- :    -
BCsl_G2   : ---------------------------------------------------------------------------------------------------- :    -
TShap1_G2 : ---------------------------------------------------------------------------------------------------- :    -
TShap2_G2 : ---------------------------------------------------------------------------------------------------- :    -
BMhap1_G2 : ---------------------------------------------------------------------------------------------------- :    -
BMhap2_G2 : ---------------------------------------------------------------------------------------------------- :    -
MRhap2_G2 : ---------------------------------------------------------------------------------------------------- :    -
PN_T2T_G3 : ---------------------------------------------------------------------------------------------------- :    -
MHhap1_G3 : ---------------------------------------------------------------------------------------------------- :    -
MHhap2_G3 : ---------------------------------------------------------------------------------------------------- :    -
SMhap1_G3 : ---------------------------------------------------------------------------------------------------- :    -
SMhap2_G3 : ---------------------------------------------------------------------------------------------------- :    -
BCsd_G3   : ---------------------------------------------------------------------------------------------------- :    -
BCsl_G3   : ---------------------------------------------------------------------------------------------------- :    -
TShap1_G3 : ---------------------------------------------------------------------------------------------------- :    -
TShap2_G3 : ---------------------------------------------------------------------------------------------------- :    -
BMhap1_G3 : ---------------------------------------------------------------------------------------------------- :    -
BMhap2_G3 : ---------------------------------------------------------------------------------------------------- :    -
MRhap2_G3 : NSTDKEGHPLCYNVCGAFKDREFYNKTFGSEAKCEDFLRWRVQSMEKVIQNLNFTAGGVDSMVQILDLKNSPRPSNKELRLVTKKAITLLQDNYPELIFR :  272
            EGGLQAVLPPRGQQERGEQQQDHFHARGNGYEETICSLRLKQNIGDPWRADVYTPRGGHRSSVTGYDLPILRKVVRLSAHQGRLHQGAMVLPYYNVNAHS

                    *        420         *        440         *        460         *        480         *        500
PN_T2T_G1 : ---------------------------------------------------------------------------------------------------- :    -
CShap1_G1 : ---------------------------------------------------------------------------------------------------- :    -
CShap2_G1 : ---------------------------------------------------------------------------------------------------- :    -
MHhap1_G1 : ---------------------------------------------------------------------------------------------------- :    -
MHhap2_G1 : ---------------------------------------------------------------------------------------------------- :    -
SMhap1_G1 : ---------------------------------------------------------------------------------------------------- :    -
SMhap2_G1 : ---------------------------------------------------------------------------------------------------- :    -
BCsd_G1   : ---------------------------------------------------------------------------------------------------- :    -
BCsl_G1   : ---------------------------------------------------------------------------------------------------- :    -
TShap1_G1 : ---------------------------------------------------------------------------------------------------- :    -
TShap2_G1 : ---------------------------------------------------------------------------------------------------- :    -
```

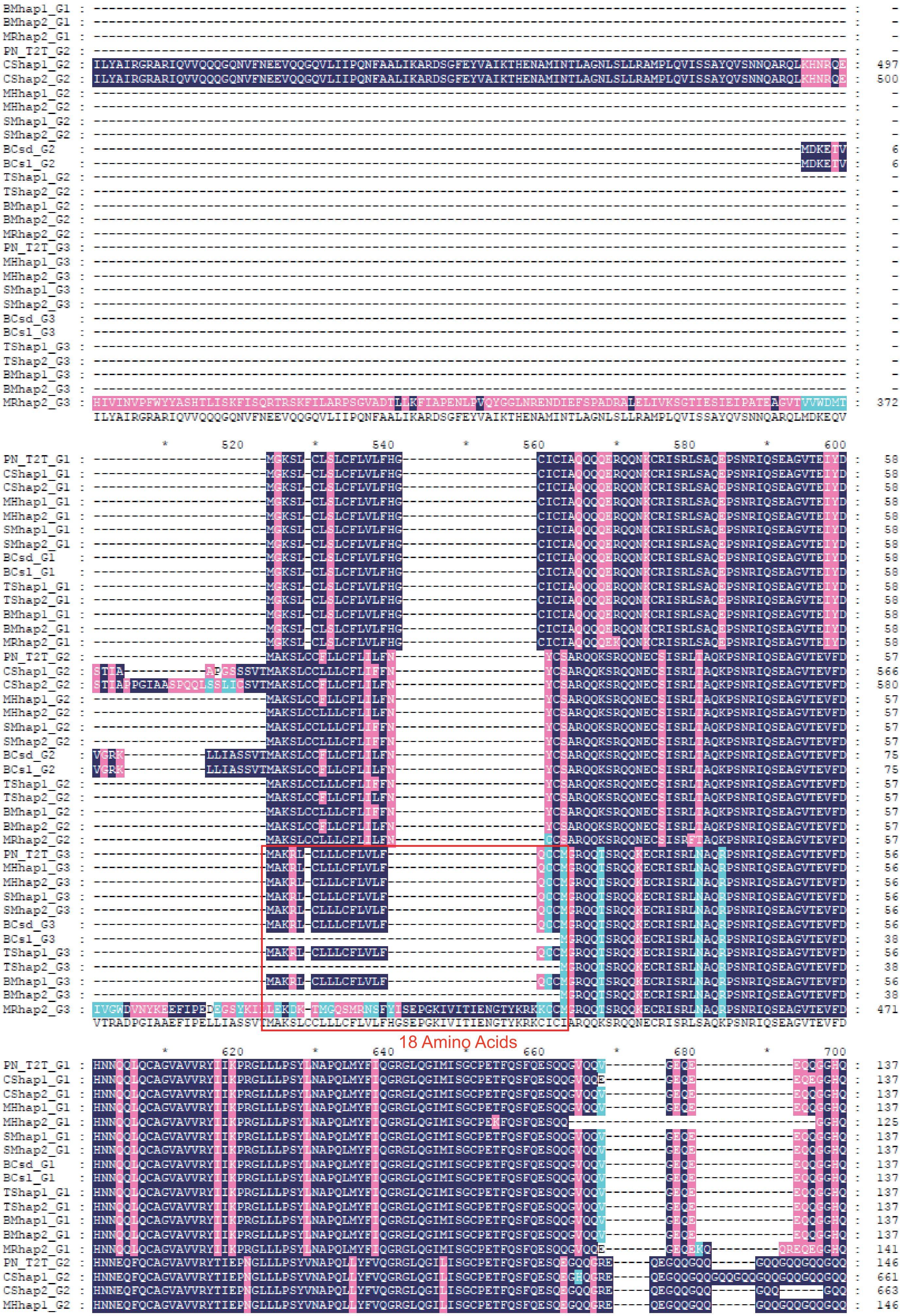
18 Amino Acids

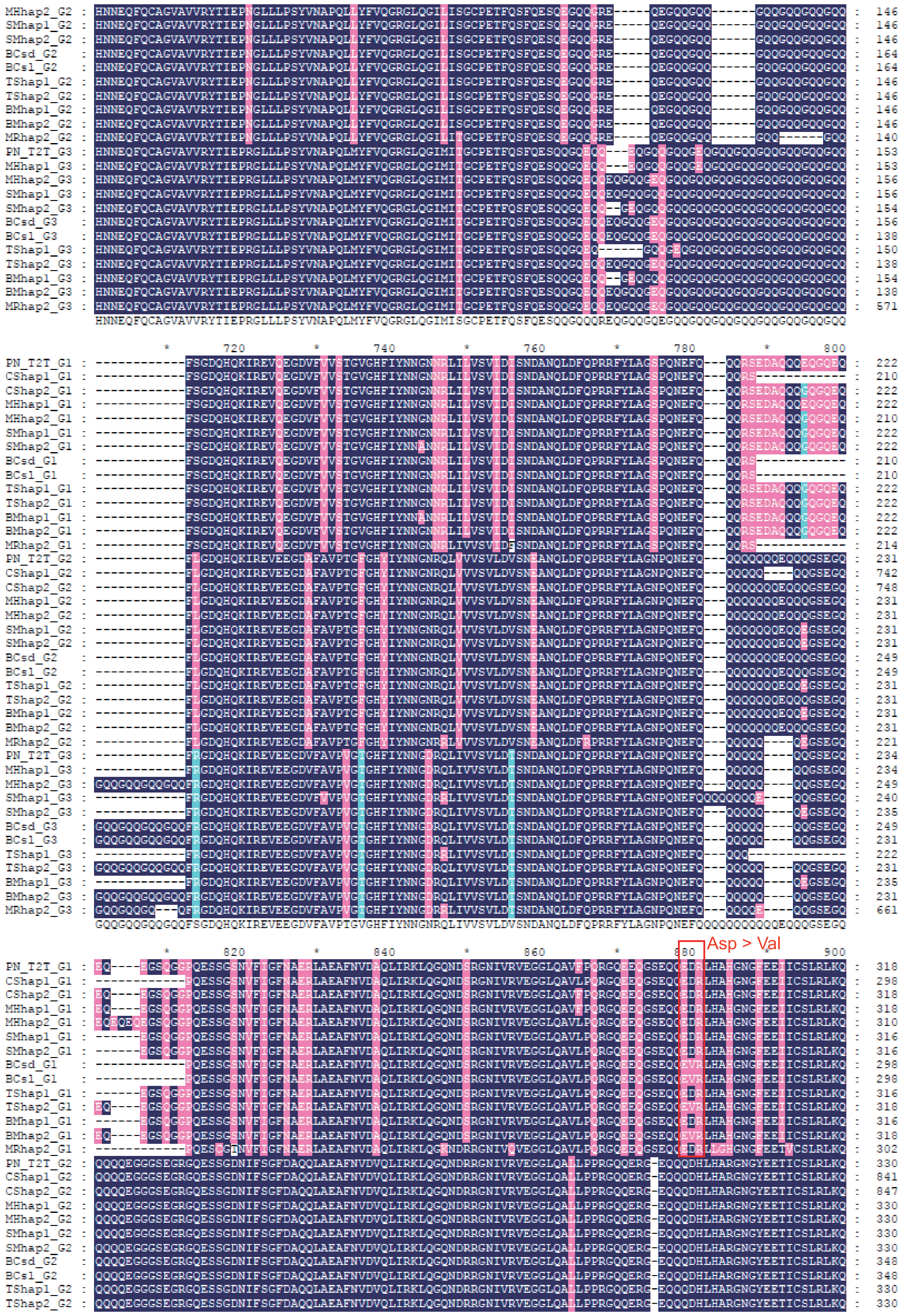

720
740
760
780
800
820
840
860
880
900
Asp > Val

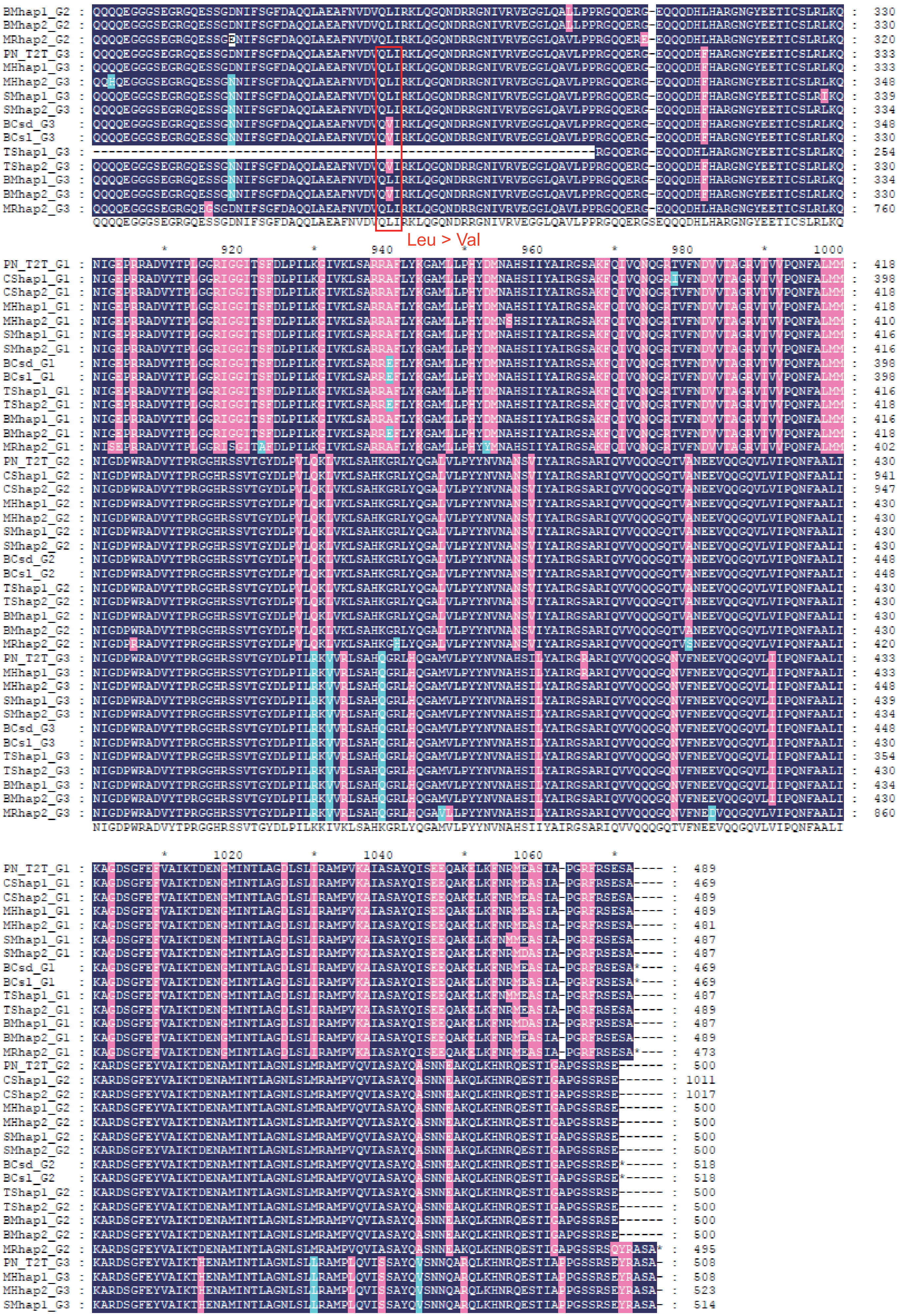
Leu > Val

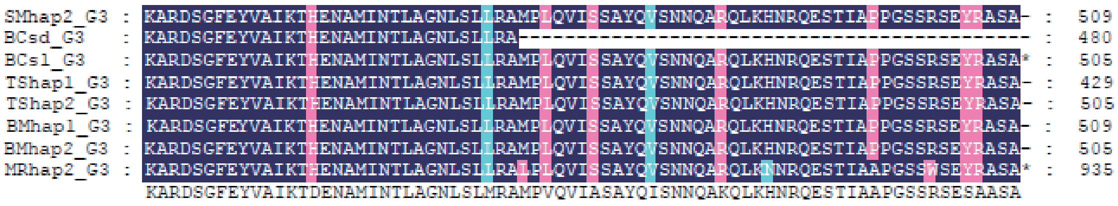


**Extended Data Fig. 9 | Sequence alignment of *11S globulin G1-G3* homologous genes from 14 grape genomes:** includes 'PN_T2T' (PN40024), 'Cabernet Sauvignon' (CS), 'Muscat Hamburg' (MH), 'Shine Muscat' (SM), 'Black Corinth Seeded' (BCsd), 'Black Corinth Seedless' (BCsl), 'Thompson Seedless' (TS), 'Black Monukka' (BM), and 'Muscadinia Rotundifolia' (MR).

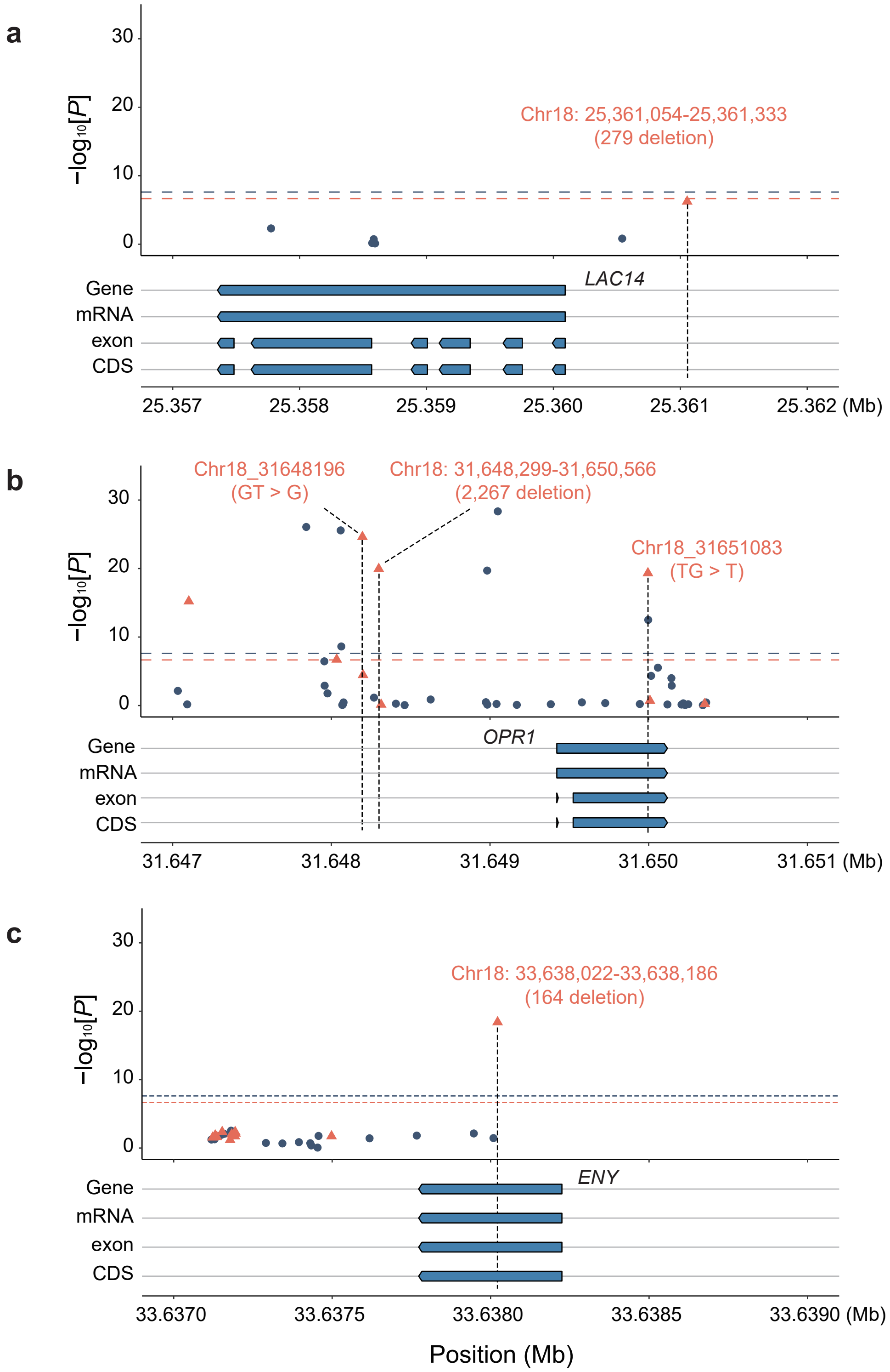


**Extended Data Fig. 10 | Three candidate genes related to seedlessness in Chr18. a-c**, Candidate genes were identified through GWAS analysis using the admixed population, with a significant threshold of 7.61 for SNPs and 6.66 for SVs (and InDels).

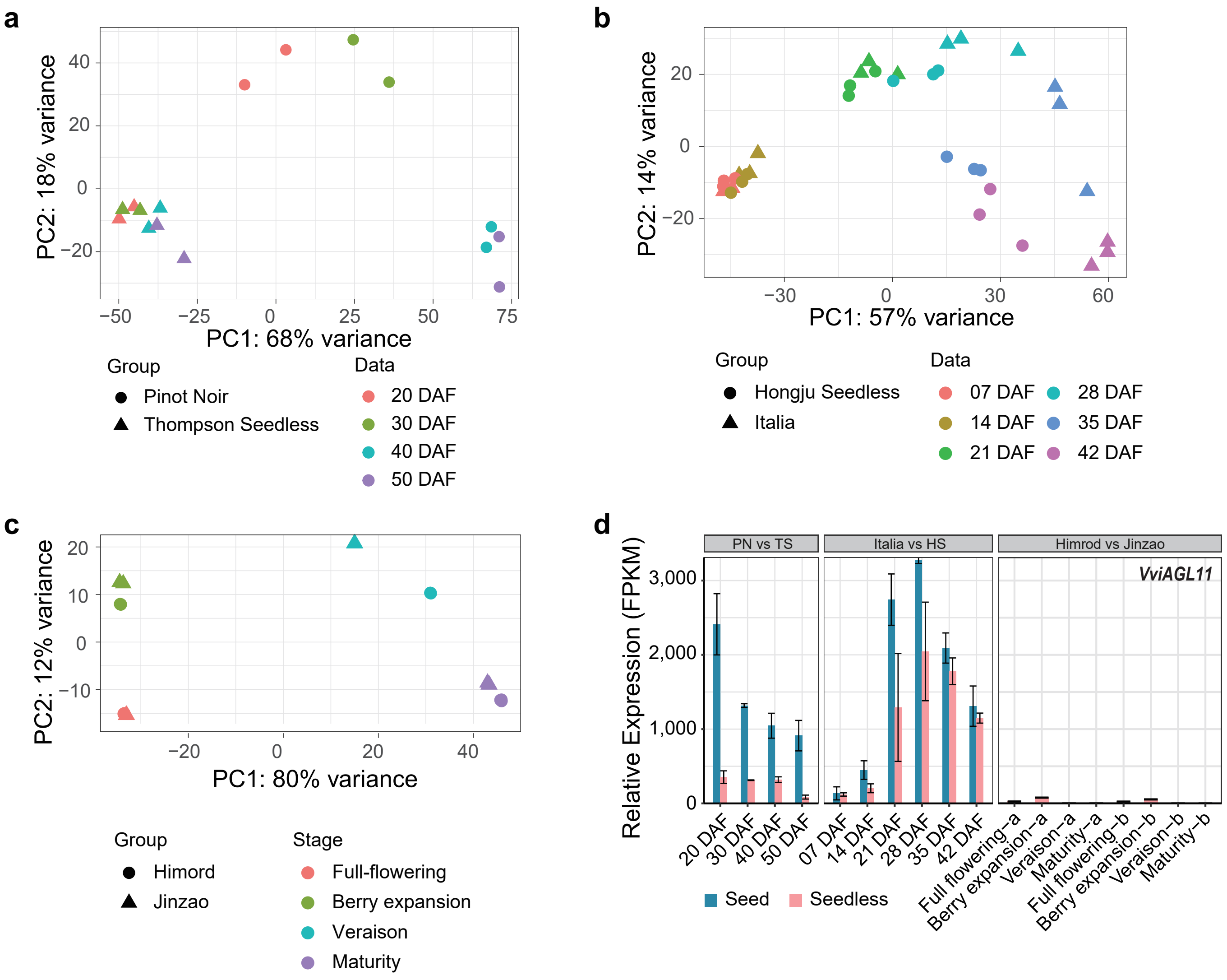


**Extended Data Fig. 11 | Principal Component Analysis (PCA) analysis of the three transcriptomic datasets, and relative expression values (FPKM) for *VviAGL11*.**

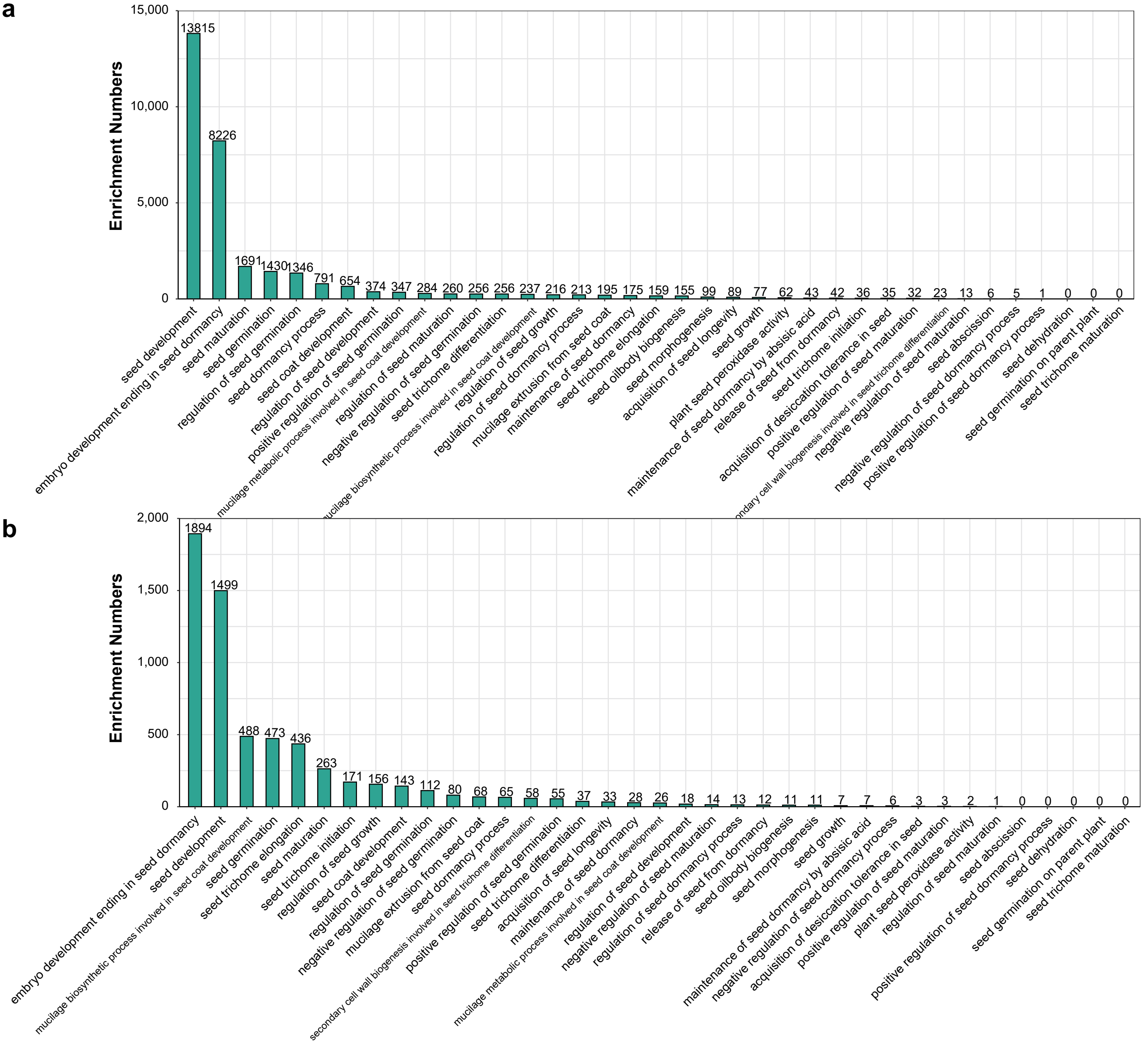


**Extended Data Fig. 12 | Enrichment analysis of GO homologous genes related to grape seed development. a**, Enrichment analysis of 14,650 seed development genes from the GO database. **b**, Enrichment analysis of 6,529 GO homologous genes associated with grape seed development.

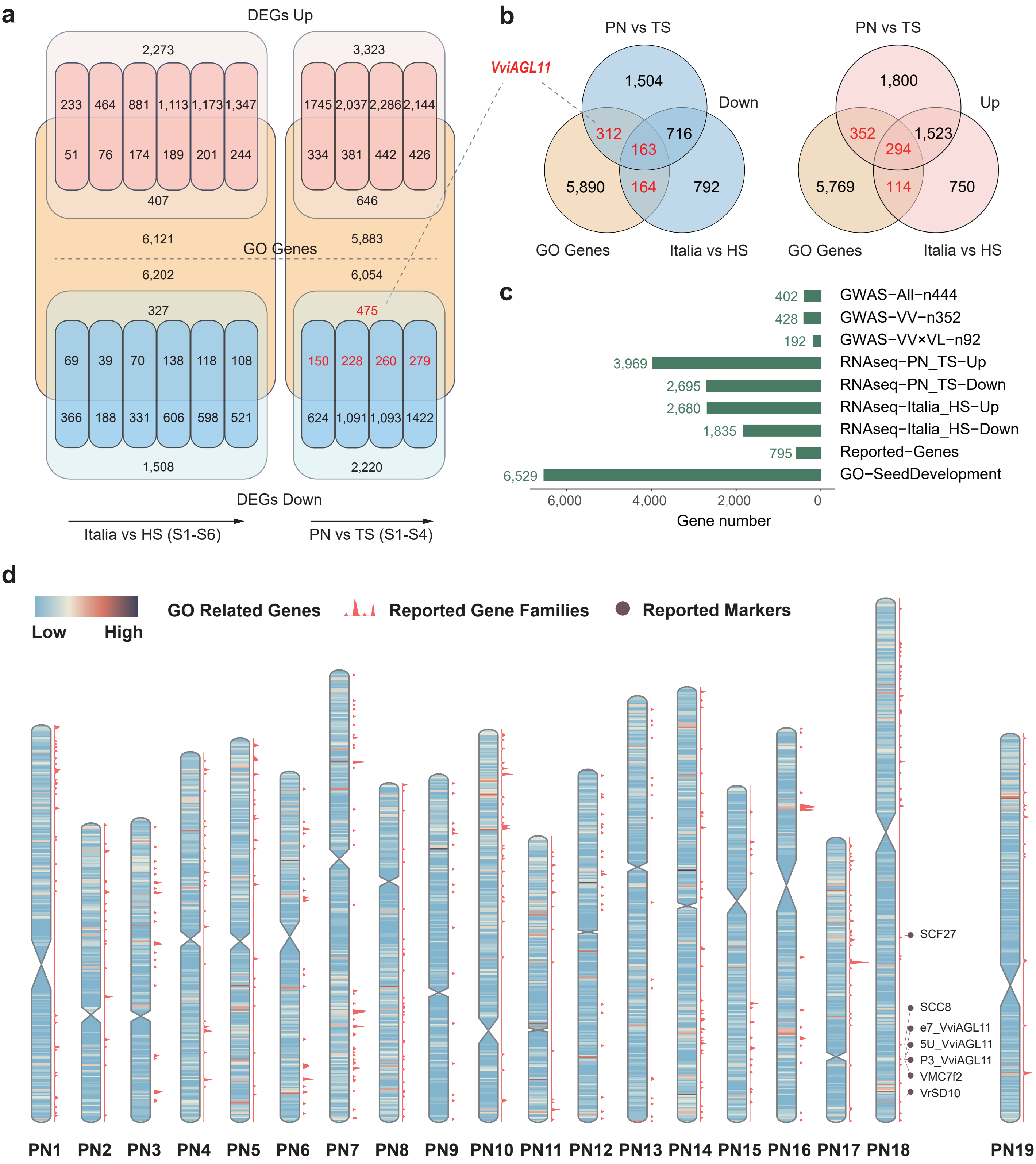


**Extended Data Fig. 13 | Visualization of multiple seed development-related datasets. a** and **b**, Results from two independent transcriptomic analyses, 'Italia' vs 'Hongju Seedless' (HS) and 'Pinot Noir' (PN) vs 'Thompson Seedless' (TS), overlapping with GO homologous gene. Red indicates up-regulated DEGs, blue represents down-regulated DEGs, and yellow denotes GO homologous genes associated with seedlessness. **c**, Results from integrative genomic analyses: GWAS, transcriptomics, reported genes mapping, and GO homologous genes. Green bars represent the total number of genes identified through this approach. **d**, Visualization of three datasets: GO homologous genes, reported gene families, and reported molecular markers.

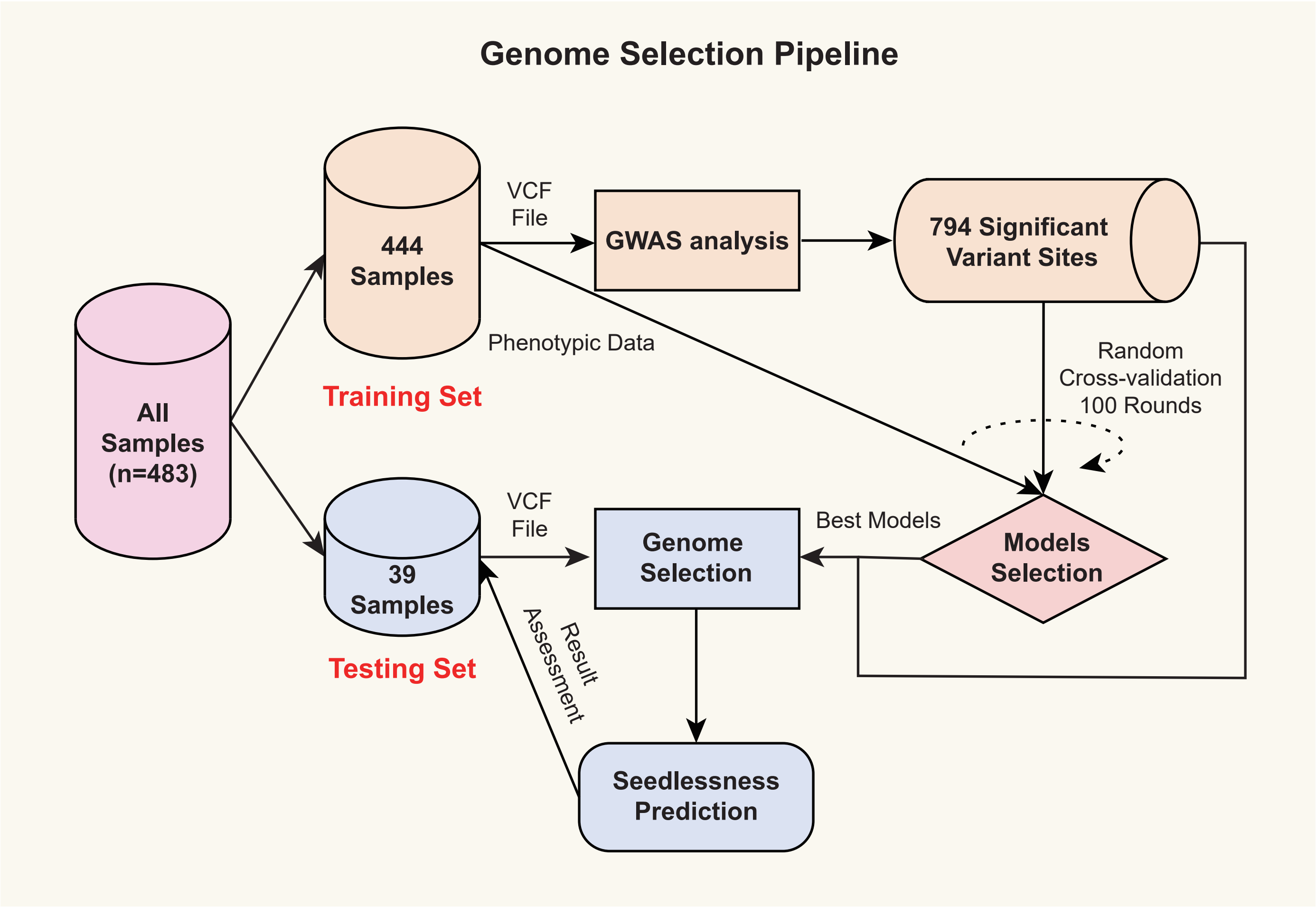


**Extended Data Fig. 14 | Genome selection workflow. The 794 significant variants extracted from GWAS results, including 77 InDels and 717 SNPs.** More detail code can be found on our lab GitHub@zhouyflab.
