## Supplementary figures and images for "Integrative genomics reveals the polygenic basis of seedlessness in grapevine"

### Figure 1-6

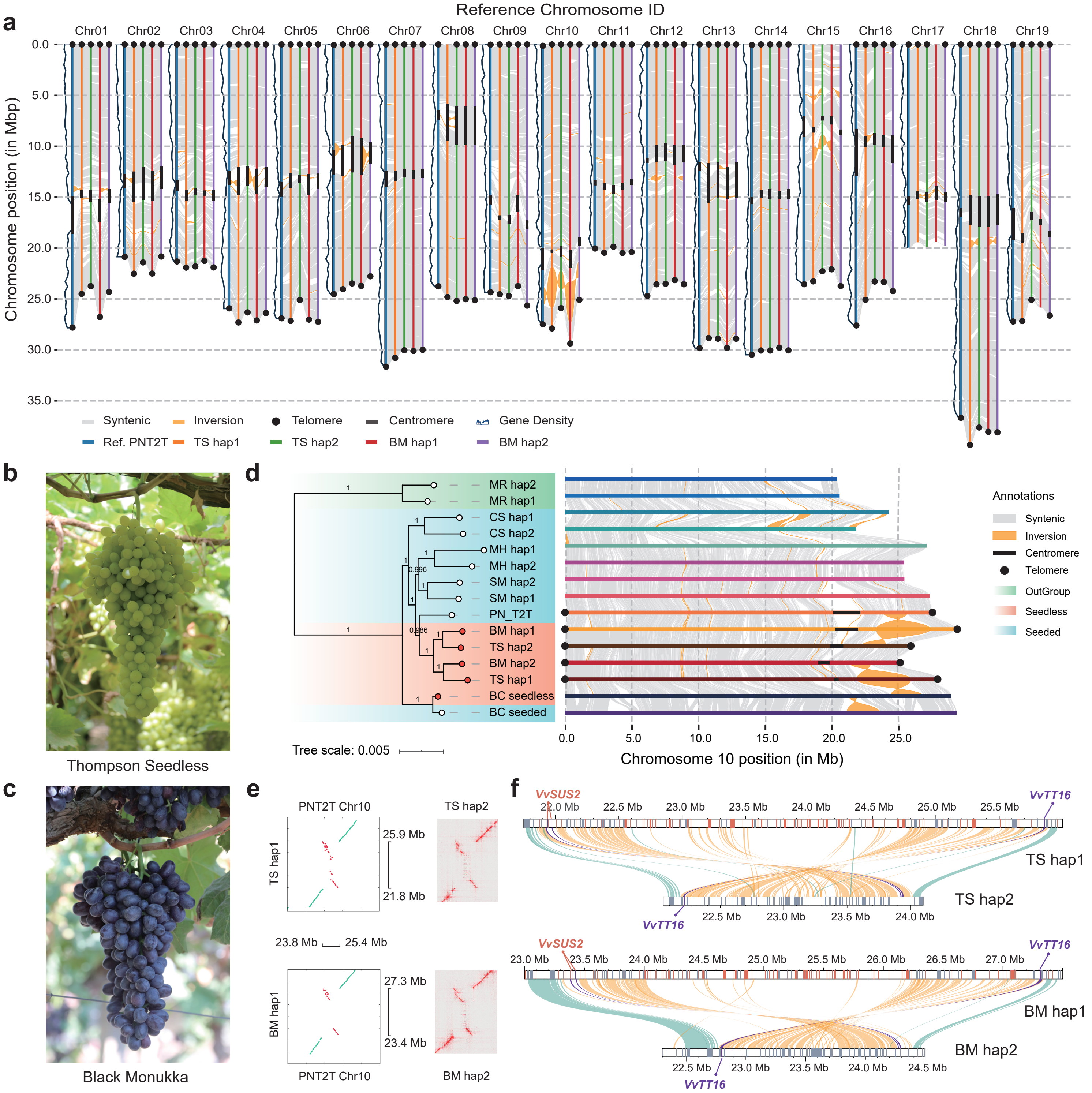

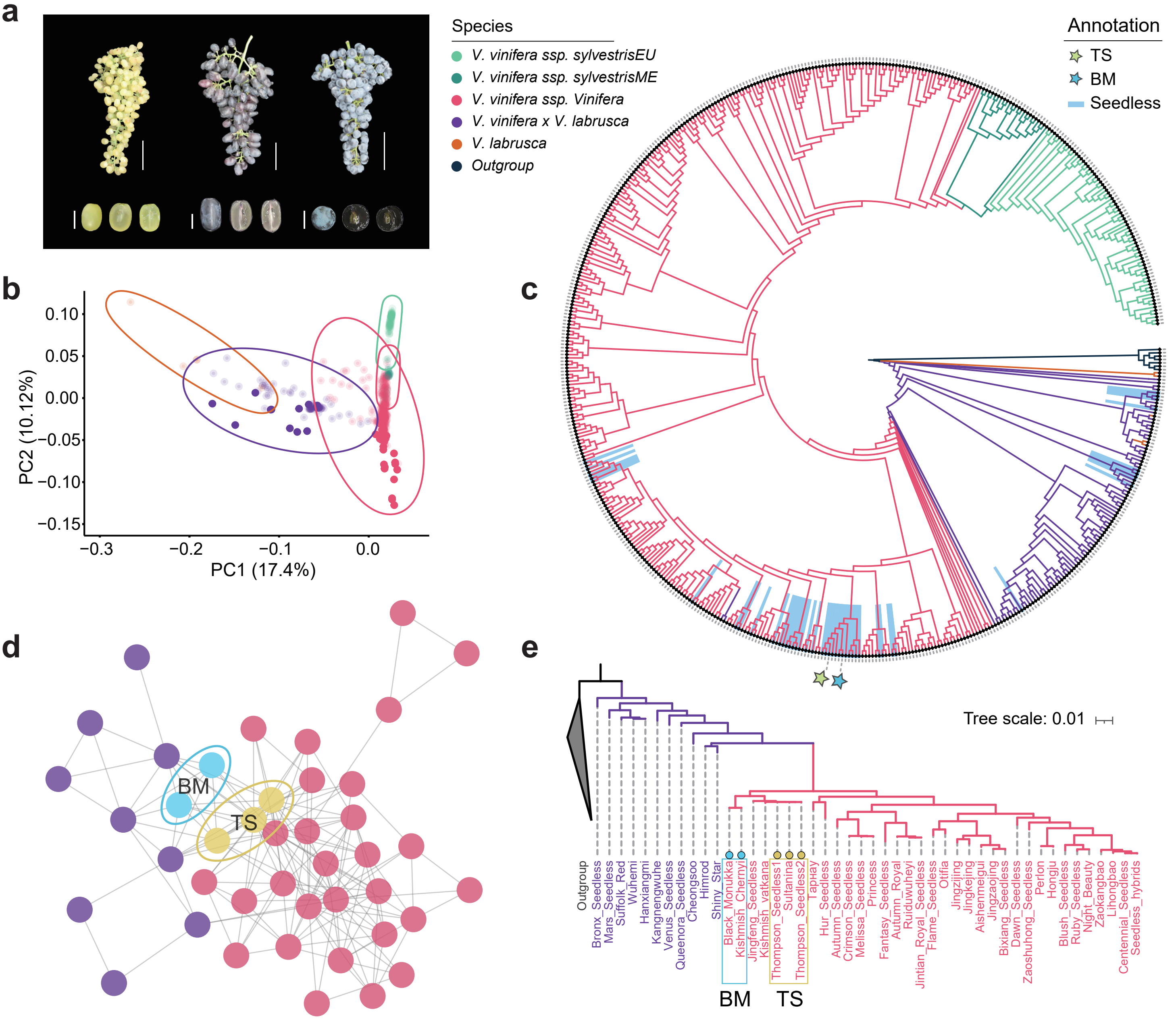

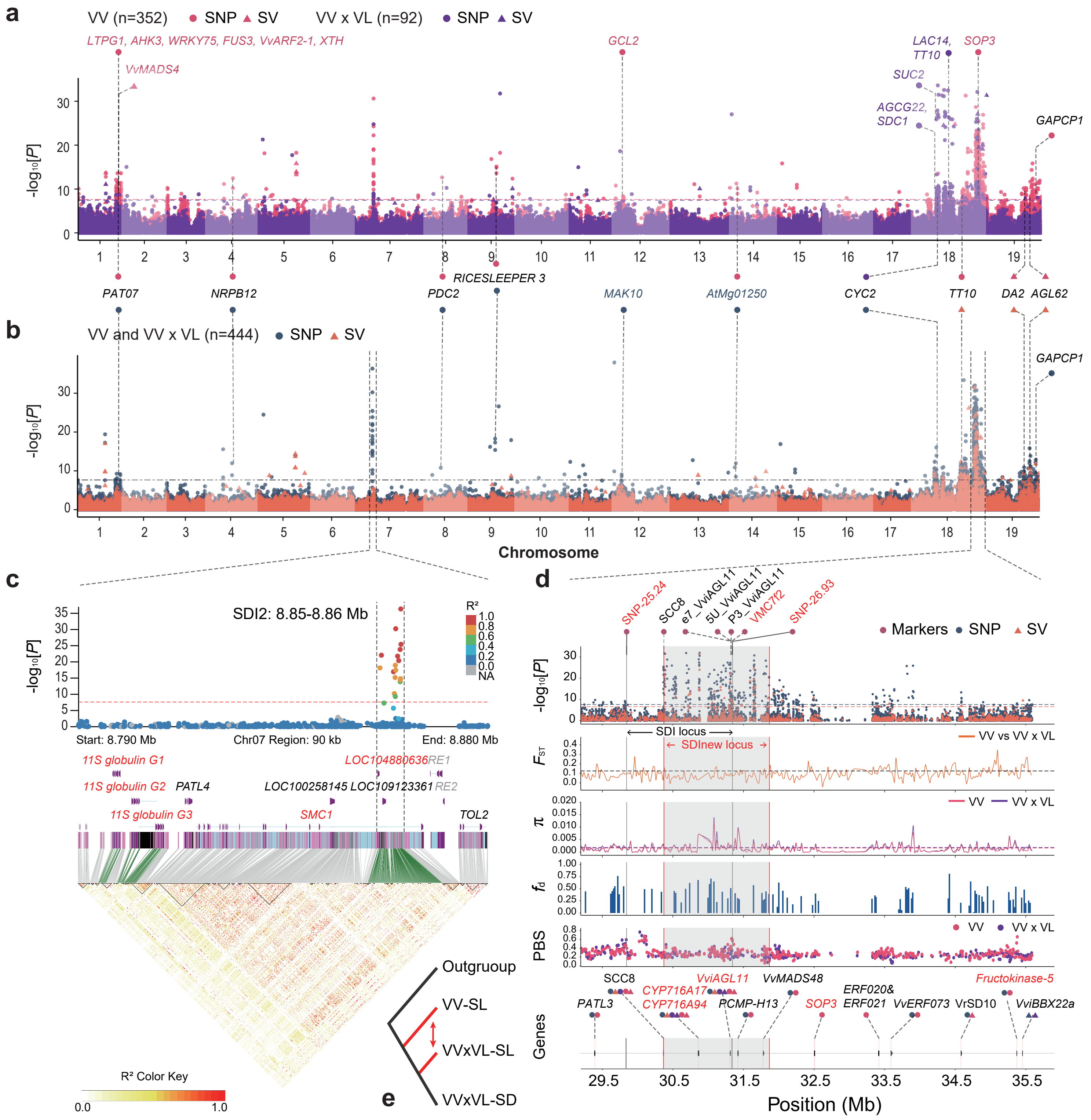

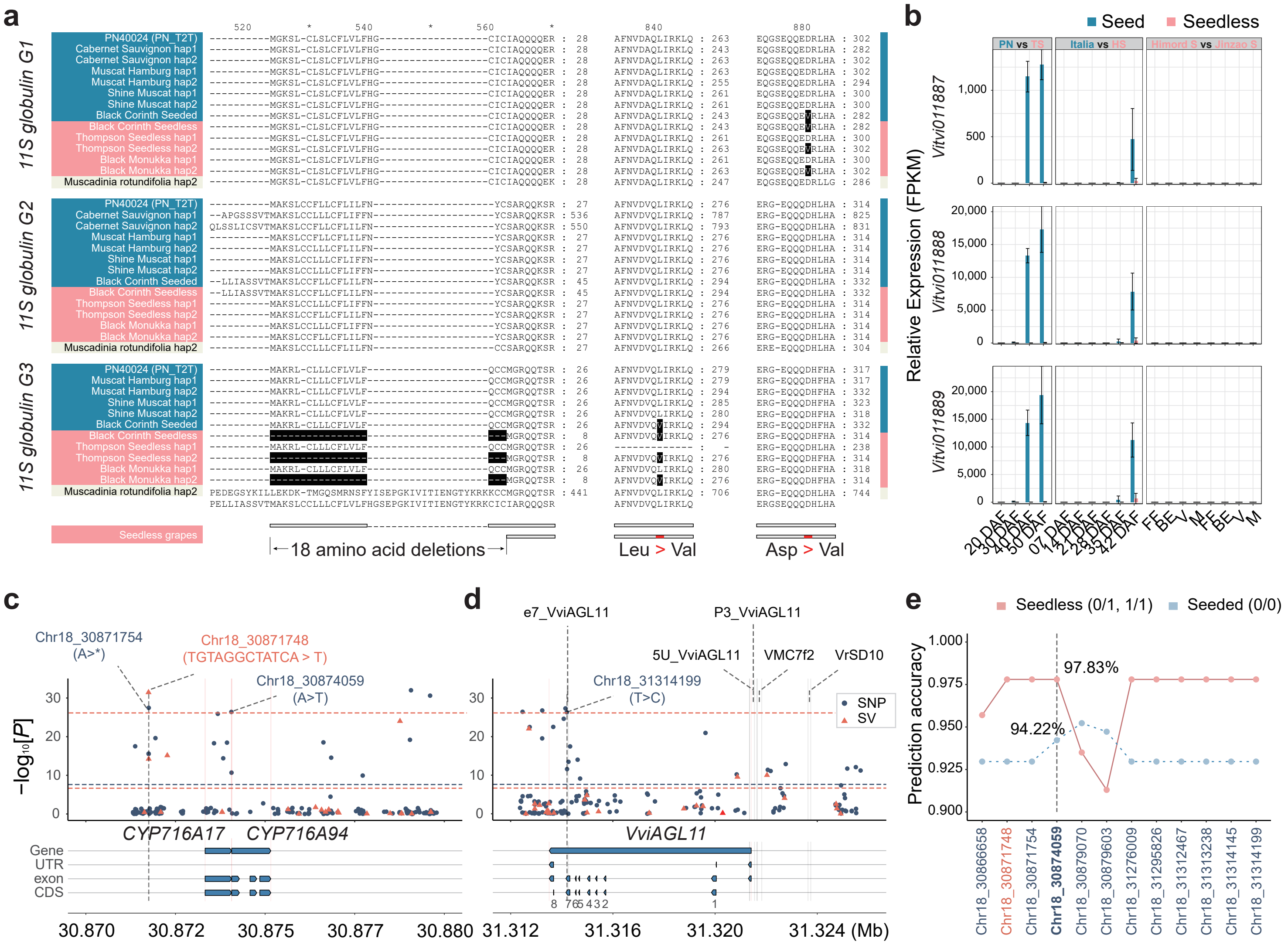

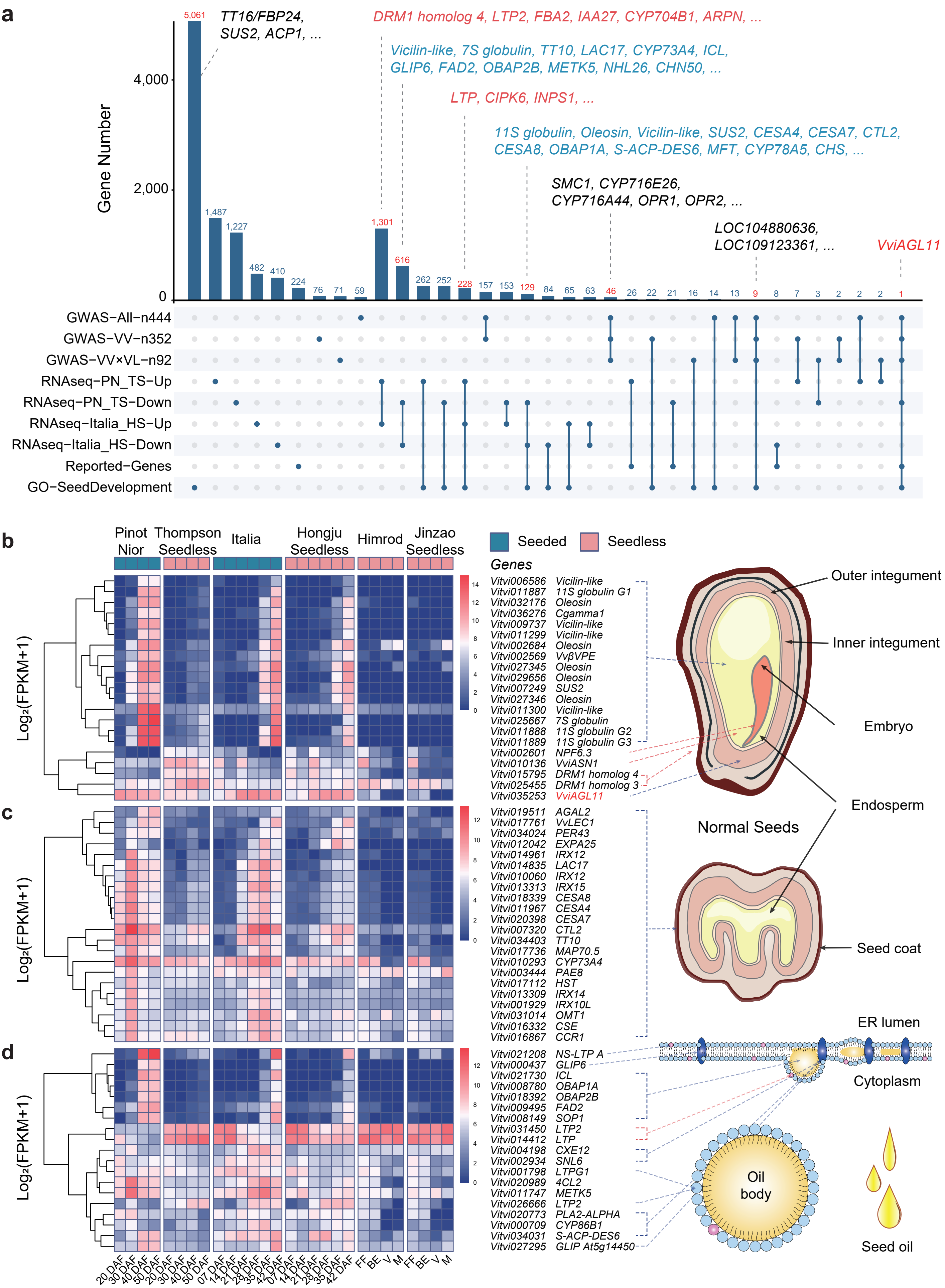

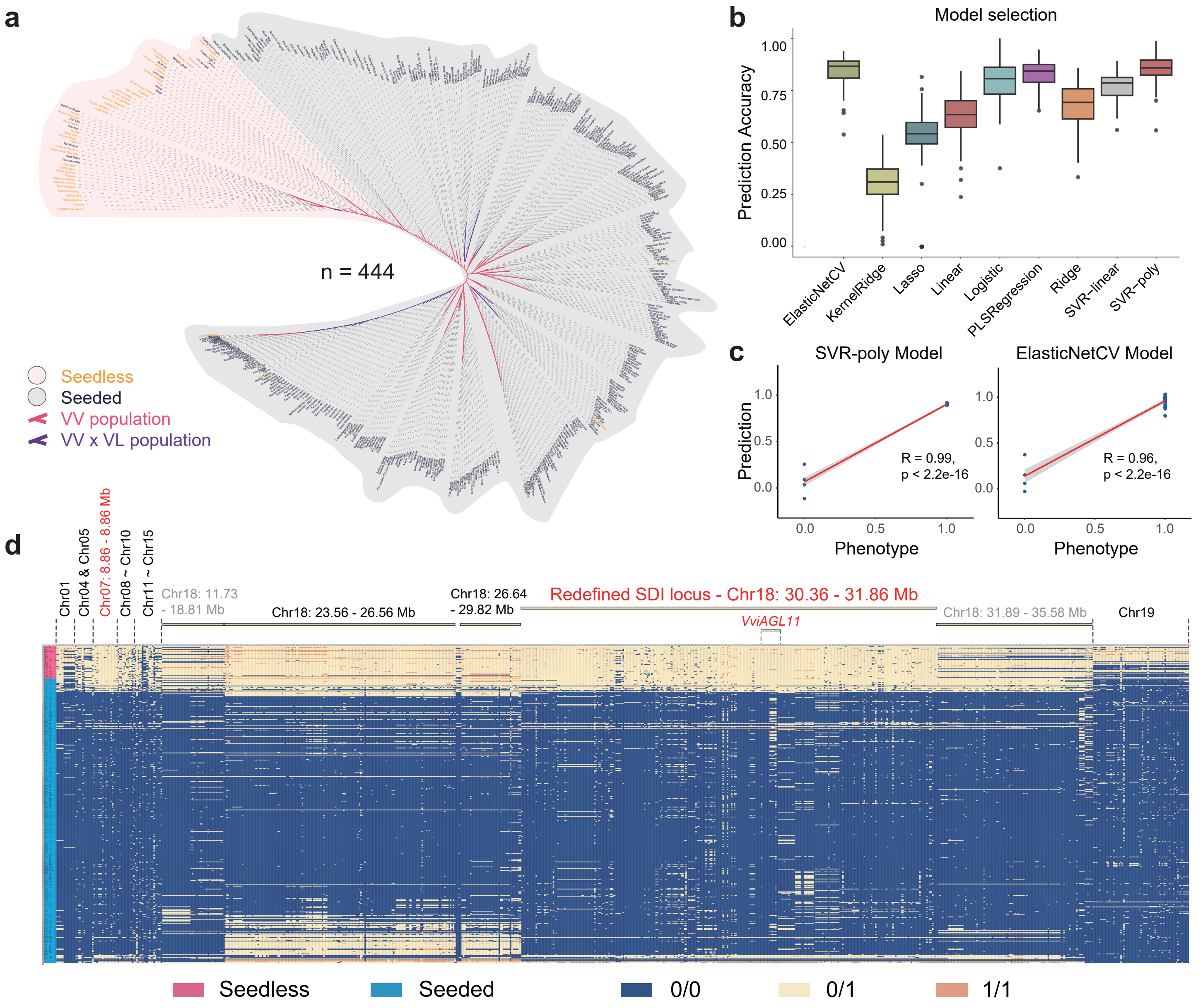
